# Getting to the core of the matter – Assessing the role of replication in metabarcoding-based *seda*DNA

**DOI:** 10.64898/2026.08.24.746646

**Authors:** Elena Baños, Clara Ras Segura, Erik J. De Boer, Andrew B. Cundy, Xavier Turon Barrera, Sandra Nogué, Luke E. Holman, Marc Rius

## Abstract

Replication is central to most experimental and sampling designs, increasing inferential power and capturing fine-scale data heterogeneity. However, its importance remains poorly evaluated in some ecological and evolutionary settings. This is the case of metabarcoding studies using DNA recovered from sedimentary archives, in which biological signals integrate ecological information through depositional and burial processes, yet are commonly inferred from a single sediment core per site.

Here, we evaluated the effect of different types of replication using sedimentary DNA metabarcoding data from two genetic markers (mitochondrial COI and nuclear 18S) using a nested sampling design. The design included three intertidal sites, three spatially separated sediment cores per site (biological replicates), two sediment horizons per core, and eight PCR (technical) replicates per sediment sample. Variance partitioning showed that site identity and sediment age group together explained >70% of the variation in beta diversity, indicating that among-site spatial and stratigraphic differences were the dominant drivers of community composition. PERMANOVA likewise identified non-significant effects of biological replication. Among PCR replicates from the same sediment sample, richness varied substantially, whereas Shannon diversity was more consistent.

Despite this variability, differences in community composition among technical replicates remained smaller than those associated with biological replication or site identity, indicating a limited influence on broader ecological patterns. Community composition was highly similar among replicate cores within sites, consistent with stratigraphic coherence.

These results indicate limited within-site heterogeneity and suggest that, under stratigraphically coherent conditions, increasing biological replication may provide little additional information, whereas enhancing technical replication and stratigraphic resolution can improve ecological inference from sedimentary DNA metabarcoding datasets.

## 1. Introduction

Experimental and sampling designs rely on appropriate replication to ensure robust ecological and evolutionary inference (Ficetola et al., 2015; Quinn & Keough, 2023). Adequate replication reduces false positives, increases statistical power, and ensures that observed patterns are not artefacts of specific samples or conditions (Fraser et al., 2020). Reproducibility problems often arise not from analytical methods but from inadequate experimental design, particularly when levels of replication are poorly defined or conflated (Underwood, 1997). Such shortcomings can lead to pseudoreplication, inflated degrees of freedom, and unreliable variance estimates, ultimately weakening ecological inference (Hurlbert, 1984; Marshall, 2024). Within this context, understanding how different levels of replication contribute to the reliability of biodiversity estimates and community patterns is critical (Mata et al., 2019; Zinger et al., 2019).

Within molecular ecology, replication is a key determinant of the robustness and reproducibility of environmental DNA (eDNA) and metabarcoding datasets (van der Loos & Nijland, 2021; Zarcero et al., 2024). Environmental DNA refers to genetic material recovered directly from environmental substrates such as water, sediment, soil, or air, and may include both intracellular and extracellular DNA originating from multiple organisms. Replication strategy can strongly influence estimates of alpha diversity and the consistency of the community composition across PCR and sample replicates (Guri et al., 2024; Jensen et al., 2024). Replication may operate at two main levels: biological (i.e., field sampling) replicates, often involving the collection of multiple sediment cores from a single sampling site, to capture spatial and temporal ecological heterogeneity (Beentjes et al., 2019; Hestetun et al., 2021; Mata et al., 2019); and technical replicates, which quantify laboratory-level variability introduced during DNA extraction, amplification, and sequencing (Shirazi et al., 2021). Although both replication levels are expected to shape metabarcoding datasets, their relative contributions are known to vary with marker choice, substrate, and sampling design (Prosser, 2010; Zinger et al., 2019). Despite recognition of the importance of different types of replication, little is known yet about their effects in metabarcoding-based studies using DNA recovered from sedimentary archives.

Sedimentary ancient DNA (*seda*DNA) represents a temporally structured subset of environmental DNA preserved within sedimentary archives (Capo et al., 2021; Ficetola et al., 2018; Parducci et al., 2017). Depending on depositional and preservation conditions, *seda*DNA records may span timescales ranging from recent historical deposits to sediments that are thousands of years old (Heintzman et al., 2023). *Seda*DNA has become increasingly important for reconstructing past biological communities because it can complement traditional palaeoecological proxies such as pollen, diatoms, and macrofossils (Capo et al., 2021; Nogué et al., 2021). DNA fragments originating from multiple organisms become incorporated into sedimentary deposits and preserved through burial processes, providing information on past assemblages across a wide range of taxa and environments (Baños et al., 2025; Foster et al., 2020; Garcés-Pastor et al., 2022; Holman et al., 2025). However, the taxonomic resolution and ecological interpretation of these records may vary depending on marker choice, reference database completeness, and preservation conditions (Freeman et al., 2023; Garcés-Pastor et al., 2023; Heintzman et al., 2023; Parducci et al., 2017). As *seda*DNA applications continue to expand across ecological and palaeoenvironmental research, increasing attention has been given to methodological consistency, reproducibility, and explicit treatment of analytical uncertainty (Capo et al., 2021; Chen & Ficetola, 2020; Heintzman et al., 2023).

The field of *seda*DNA encompasses a wide range of methodological approaches. Shotgun-based palaeogenomic *seda*DNA studies commonly assess molecular authenticity through characteristic fragment length distributions and post-mortem nucleotide damage patterns (Everett & Cribdon, 2023; Zimmermann et al., 2023), whereas metabarcoding-based *seda*DNA studies are primarily designed to recover broad ecological community signals rather than authenticate individual ancient molecules (Holman et al., 2025; Zimmermann et al., 2024). Consequently, these metabarcoding-based *seda*DNA studies are particularly useful for comparative ecological analyses, although their interpretation should acknowledge limitations associated with PCR amplification biases and differential DNA preservation (Elbrecht & Leese, 2015; Shaffer et al., 2025).

In aquatic environments, eDNA studies often show strong fine-scale spatial variability driven by environmental heterogeneity and DNA transport, making biological replication essential (Andruszkiewicz et al., 2017; Hestetun et al., 2021). In these systems, field replication often explains more variation than technical replication (Mata et al., 2019). Sedimentary archives, however, may integrate biological signals through depositional and burial processes across broader spatial and temporal scales (Alsos et al., 2018; Harrison et al., 2019; Heintzman et al., 2023; O’Reilly-Berkeley et al., 2026). Consequently, the degree of small-scale spatial heterogeneity in sedimentary DNA assemblages may differ from that observed in aquatic eDNA, depending on the depositional environment, sediment characteristics, and temporal integration of biological signals (Ataman et al., 2025; Capo et al., 2021; Holman et al., 2019).

Here, we explicitly test the relative importance of both biological and technical replication in metabarcoding-based *seda*DNA datasets derived from dated sediment archives. Using mitochondrial and nuclear markers to analyse samples collected at multiple sampling sites, we assess whether replicate sediment cores collected within sites yield distinguishable estimates of richness, Shannon diversity and community composition, and evaluate the contribution of technical replication to these patterns. Site identity was included as a broader spatial context within which biological and technical replication effects were evaluated, allowing local variability among cores to be distinguished from larger-scale spatial differences among sampling locations. By partitioning variance among biological replicates, technical replicates, site identity, and sediment age group, we provide empirical evidence of the relative contribution of each replication type and propose a framework to optimise the sampling design and strengthen ecological inference in metabarcoding studies using DNA recovered from sedimentary archives.

## 2. Materials and Methods

### 2.1. Study sites and field sampling

We collected sediment cores from three intertidal saltmarsh sites [Cicero (CCO), Colindres (CLR), and Santoña (STO)] within the Santoña estuarine system on the northeast Atlantic coast (Fig. 1, Table S1) in November 2023. This wetland is characterised by fine-grained sediments with high organic matter content under tidal influence, which may influence DNA retention and preservation processes (Baños et al., 2025; Campbell et al., 2025; Foster et al., 2020; Irabien et al., 2008). At each sampling site, we used a 50 cm Russian-type peat corer to collect three sediment cores (i.e., biological replicates) randomly placed approximately 50 cm apart from each other. Sediment sections were transferred to semi-cylindrical PVC tubes (5 cm internal diameter), transported to research facilities at 4 °C within 72 h of collection. We then processed the cores in a dedicated clean DNA laboratory (PCR-free, positive-pressure facility) that was specifically designed to minimise contamination. Standard clean-lab procedures were followed throughout sample processing, including the use of dedicated protective clothing and spatial separation between subsampling and pre-PCR activities. Work surfaces and instruments were decontaminated with 10% commercial bleach (∼0.5–0.6% sodium hypochlorite), rinsed with ultrapure water, and cleaned with 70% ethanol prior to core processing.

**Fig. 1.**
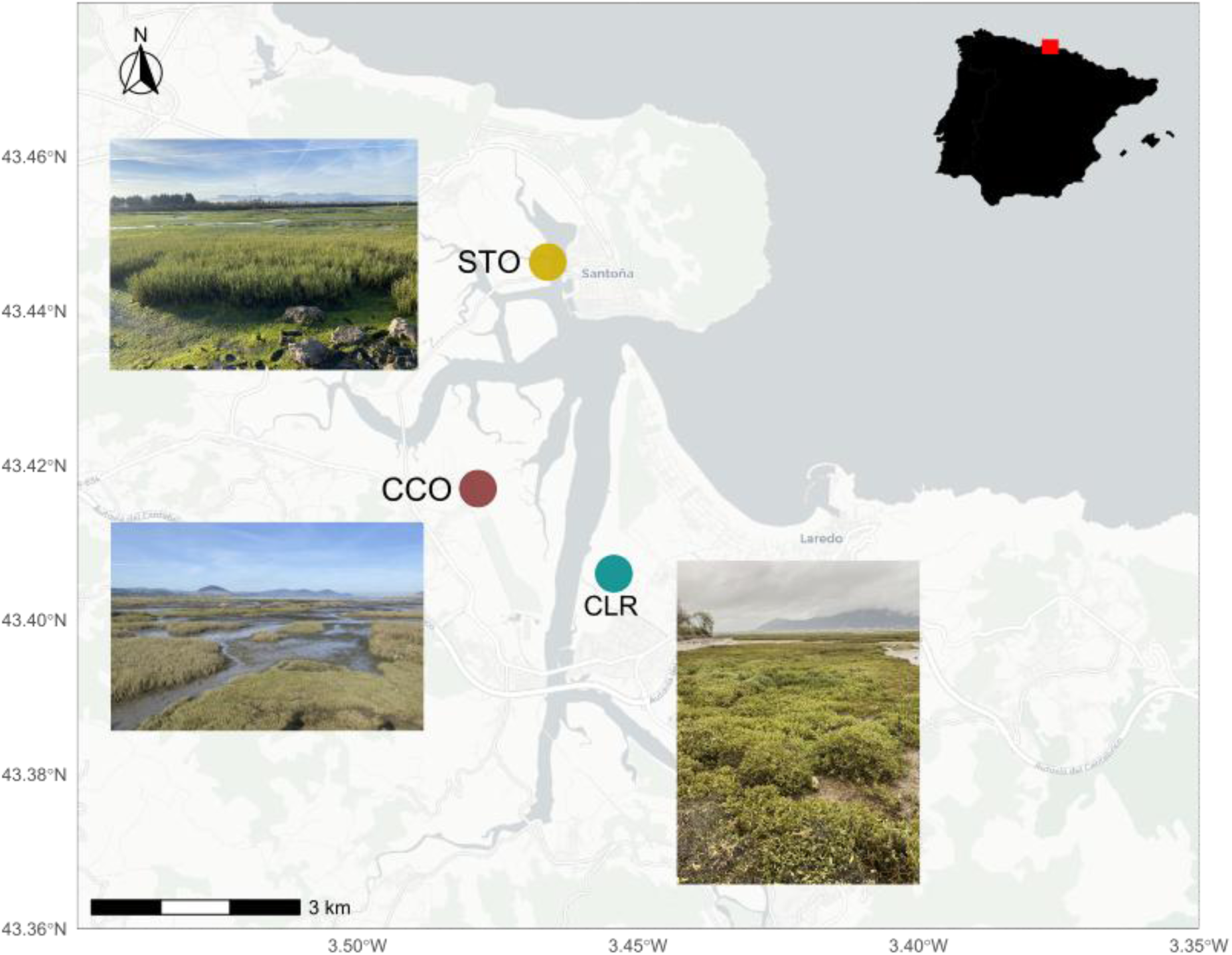
Geographic location of the three intertidal study sites [Cicero (CCO), Colindres (CLR) and Santoña (STO)] within the Santoña estuary. Photographs illustrate the general environmental characteristics of the sampled intertidal sites. The inset indicates the broader regional context of the Iberian Peninsula, while the main panel shows a site-level overview of the sampling sites.

Subsampling of the cores was carried out using sterile tools after removal of the outer sediment layer, and 2 to 5 g of sediment per horizon were transferred to 20 mL sterile containers. Gloves were changed frequently and new sterile materials were used for each subsample. Each core was subsampled at two depth horizons representing recent and old sediment layers, respectively: STO (5 and 50 cm), CCO (5 and 45 cm), and CLR (4 and 45 cm). All subsamples were stored at −20 °C until DNA extraction

### 2.2. Dating and X-ray fluorescence core scanning

The chronological context for the sediment profiles was established using radiometric dating of one sediment core per site, corresponding to the same shallow and deep horizons selected for metabarcoding-based *seda*DNA analyses. The selected core was chosen based on visual stratigraphic integrity and preservation, was stratigraphically consistent with the other two replicate cores at each site, and was used to distinguish recent and old sediment intervals.

The dried sediment samples were radiometrically dated via ^210^Pb through a proxy method using measurement of its granddaughter radionuclide ^210^Po by alpha spectrometry. The method followed Flynn (1968), in which an aliquot of sediment was spiked with ^209^Po as a chemical yield tracer and digested using aqua regia. Polonium was subsequently autodeposited onto a cleaned silver disc and measured using Ortec Alpha Octete spectrometers equipped with PIPS detectors to determine ^210^Po activity. The analytical detection limit was 0.1 Bq kg⁻¹.

Sediment accumulation rates were estimated using the Constant Flux: Constant Sedimentation (CF:CS) or ’Simple’ model (Robbins, 1978), based on a linear regression between the natural logarithm of the unsupported ^210^Pb activity (^210^Pb_ex_) and core depth. Supported ^210^Pb activities were estimated from the mean activity measured at the base of each core, where ^210^Pb activity approaches constant background values (i.e., supported activities) in older sediments (Chepstow-Lusty et al., 2007).

Elemental composition was analysed for each biological replicate using a Niton™ XL3p+ handheld X-ray fluorescence scanner (Thermo Fisher Scientific, USA). Freeze-dried, homogenised samples (∼0.5 g) were measured for 180 s at 1-cm intervals and elemental concentrations were reported as proportional values (%) for elements detected above a threshold of 0.001%. This non-destructive method allows elemental characterisation of sediment cores (Croudace et al., 2019). Elemental profiles (Fe, Rb, Ti, Zr) were expressed in parts per million (ppm) and normalised to total scatter (Fisher et al., 2025; Kylander et al., 2011) and were z-standardised within each core. The pairwise Pearson correlation coefficients (r) between the cores were calculated within the sites using shared depth intervals, ensuring that the comparisons were restricted to stratigraphically equivalent sections despite differences in total core length or sediment recovery. Summary metrics (median, minimum, maximum r) were used to quantify reproducibility.

### 2.3. DNA extraction

DNA was extracted from two depth horizons for each biological replicate, resulting in six sediment samples per site (three cores × two depths) using the DNeasy PowerMax Soil Kit (QIAGEN, Hilden, Germany), following the manufacturer’s protocol with a final elution volume of 1.5 mL. We performed all DNA extractions in the same clean DNA laboratory described above under strict contamination control procedures. Contamination was monitored using negative controls at each step: three subsampling blanks consisting of sterile sediment processed alongside samples, three extraction blanks containing sterile sediment, and PCR blanks (n = 8 per plate; two plates per marker, COI and 18S). All blanks (n = 22 per marker) were carried through the entire workflow and sequenced along with the samples. DNA concentration was quantified using a Qubit High Sensitivity Assay Kit (Thermo Fisher Scientific), and extracts were stored at −20 °C.

### 2.4. Primer selection and library preparation

Two metabarcoding regions were targeted: a 313-bp fragment of the mitochondrial cytochrome c oxidase I (COI) gene using the Leray-XT primer pair (mlCOIintF-XT / jgHCO2198) (Wangensteen et al., 2018); and the V7 region of the nuclear 18S rRNA gene (∼110 bp) with 18S_allshort primers (Guardiola et al., 2015) (Table S2). Although the COI fragment is longer than those commonly used in studies targeting highly degraded DNA (Dabney et al., 2013), it was selected because of its broad taxonomic coverage and its widespread use in ecological monitoring (Atienza et al., 2020). Given that this study employed metabarcoding-based *seda*DNA of relatively recent historical sediments rather than ancient DNA authentication, maximising taxonomic resolution was prioritised over recovering the shortest possible DNA fragments. The sediment horizons analysed in this study correspond predominantly to historical timescales (the last ∼500 years), where extreme DNA fragmentation characteristic of older sedimentary archives may be less pronounced. All PCR products were dual indexed following a twin tagging approach (Bohmann et al., 2022; Kircher et al., 2012), with 8-bp barcodes (minimum Hamming distance ≥3) attached to the 5’ ends of both primers. Identical tag sequences were used on forward and reverse primers (mirrored tags) to minimise tag-jump artefacts and misassignments during library construction (Schnell et al., 2015).

For each genetic marker, eight independent PCR replicates (that is, technical replicates) were performed per sample in 40 µL reactions containing 4 µL of *seda*DNA template, 20 µL AmpliTaq Gold 360 Master Mix (Thermo Fisher Scientific), 11.68 µL laboratory water, 0.32 µL BSA (20 mg/mL; Fisher Scientific) and 2 µL of each primer (5 µM). The thermal cycling conditions consisted of 35 cycles after initial denaturation at 95 °C for 10 min, with marker-specific extension times (COI: 94 °C 1 min, 45 °C 1 min, 72 °C 1 min; 18S: 95 °C 30 s, 45 °C 30 s, 72 °C 30 s), and a final extension at 72 °C for 5 min.

PCR products were purified using the MinElute PCR Purification Kit (QIAGEN, Hilden, Germany), amplicon concentrations were quantified with Qubit, normalised to equimolar concentrations, and pooled. In total, 144 samples per marker were generated (3 sites × 3 cores × 2 depths × 8 PCR replicates). These, together with the corresponding negative controls, were distributed across two sequencing libraries. Each library contained 96 samples, including biological samples and the corresponding negative controls. Libraries were prepared using the TruSeq PCR-free kit (Illumina) with bead ratios of 0.13 (COI) and 0.16 (18S), pooled, and sequenced on an Illumina NovaSeq 6000 platform (COI: 2 × 250 bp; 18S: 2 × 150 bp). Due to the low DNA concentration of the negative controls, blanks were included in sequencing libraries without equimolar normalisation and added at the same volume (20 µL) as biological samples.

### 2.5. Bioinformatics

Raw paired-end reads were demultiplexed using Cutadapt v2.3 (Martin, 2011) with the following specifications: sample-specific dual barcodes, combined tag–primer sequences (∼36 bp) and allowing up to 10% mismatches (--error rate 0.1; no indels). Reads were separated into sense and antisense orientations (--pair adapters, --pair filter=both) and processed independently. Twin-tagged amplicons were recovered in both sense and antisense orientations after demultiplexing. The two orientation-specific read sets were processed independently throughout the DADA2 v1.12 workflow (Callahan et al., 2016). Reads were quality filtered (maxEE = 1, truncQ = 2), trimmed (COI: 250/230 bp; 18S: 120/110 bp), denoised, merged (minimum overlap 18 bp) and chimera screened (*removeBimeraDenovo*). The antisense reads were reverse-complemented and merged with the sense reads to generate the final sequence table.

Amplicon sequence variants (ASV) were curated using LULU (Frøslev et al., 2017) implemented through a custom R function (ApplyLulu, doi:10.5281/zenodo.4671710), with minimum_match = 98. Co-occurrence and similarity-based curation was performed using *vsearch* (Rognes et al., 2016) in self-matching mode (--usearch_global, --self), with a minimum identity threshold of 0.84 (--id 0.84, --iddef 1), a minimum query coverage of 0.9 (--query_cov 0.9), and a maximum of ten hits per query (--maxhits 10).

The taxonomic assignment of both markers was performed using BLAST+ v2.14.1 (Camacho et al., 2009) against the NCBI *nt* database (March 2024). Up to 300 hits per query were retained (-num_alignments 300). The resulting BLAST outputs were parsed in R using a custom script (ParseTaxonomy, doi:10.5281/zenodo.4671710), applying identity and coverage thresholds and resolving multiple matches using a lowest common ancestor approach. The high-confidence thresholds for species-level assignment were defined as >97% identity and >85% coverage for COI and 100% identity with >98% coverage for 18S. Records labelled as uncultured, environmental, construct, clone, or sp. were removed prior to downstream analyses. ASVs showing multiple equivalent high-confidence matches were retained and resolved using a Lowest Common Ancestor (LCA) approach.

The ASV table was filtered by removing observations with <3 reads and ASVs occurring in a single PCR replicate. Potential contamination was addressed using a maximum threshold approach based on negative controls. For each ASV, the highest read count observed across all negative controls was subtracted from the corresponding ASV counts across samples. This conservative procedure was applied prior to downstream analyses and therefore affected all subsequent diversity estimates, including richness calculations. The approach assumes that low-level contamination patterns represented in negative controls are broadly comparable across samples and was intended to minimise the influence of background contamination and index cross-talk while retaining biologically meaningful low-abundance signals. ASVs were retained for community analyses regardless of their level of taxonomic assignment.

### 2.6. Statistical analyses and data interpretation

All analyses were performed in R v4.3.1 (R Core Team, 2025). Community diversity and structure were evaluated using the vegan package (v2.7-2) (Oksanen et al., 2020). To minimise biases associated with sequencing depth variation, samples with fewer than 1,000 reads were excluded from downstream analyses. Sampling completeness was further evaluated using coverage-based analyses implemented in iNEXT (v3.0.2) (Hsieh et al., 2016). Sample coverage values were consistently close to 1 across PCR replicates for both genetic markers, and estimated asymptotic richness closely matched observed richness, indicating that sequencing depth was sufficient to recover the detectable diversity within individual PCR replicates. As a sensitivity analysis, richness was additionally estimated after sample-size rarefaction using a common sequencing depth for each marker (20,662 reads for 18S and 10,540 reads for COI, corresponding to the minimum retained library size). Richness estimates were calculated as the mean of 100 independent rarefaction replicates per PCR replicate. The mixed-effects richness models were then repeated using these rarefied richness estimates to evaluate whether sequencing-depth standardisation affected the biological interpretation of the results.

We tested the role of technical replication at the PCR level, with multiple independent PCRs generated from the same DNA extract for each sample. Other potential sources of technical variation, such as extraction replicates or differences between sequencing runs, were not explicitly evaluated.

Alpha diversity was calculated using both the Shannon diversity index and ASV richness, calculated at the level of individual technical replicates. Shannon diversity was calculated from relative abundance data, while richness was defined as the number of detected ASVs per replicate (i.e., presence–absence), and was therefore not affected by relative abundance normalisation. These metrics were calculated directly from the sequencing data. To test the role of technical and biological replication, linear mixed-effects models (lme4 v1.1-38) were fitted including the site and sediment age group as fixed effects. Technical replication was modelled as a source of variability within the sample rather than as a fixed effect, reflecting the hierarchical structure of the data. Sample identity (Site:Core_Rep:AgeGroup) was included as a random intercept, with PCR replicates nested within samples: *Diversity ∼ Site + AgeGroup + (1 | Site:Core_Rep:AgeGroup)*. Variance components associated with the random effect at the sample level were used to quantify variability between samples, while intraclass correlation coefficients (ICC) were used to estimate repeatability among PCR replicates. ICC values were calculated from fitted mixed-effects models using the icc() function of the *performance* package, based on the estimated variance components of the random effects relative to total model variance.

To examine biological replication across markers, diversity metrics were analysed in two complementary modelling approaches. PCR replicates were retained as individual observations rather than averaged or collapsed prior to modelling, allowing technical variability to remain explicitly represented in the hierarchical structure of the data. First, to test whether biological replicates (cores) differed within each site, sediment age group, and marker, the biological replicate was included as a fixed effect nested within each Marker × Site × AgeGroup combination. This allowed direct pairwise comparisons among cores within each sample context using the model structure: *Diversity ∼ Marker * Site * AgeGroup + Marker:Site:Core_Rep:AgeGroup*. Second, to assess broader ecological patterns, Core_Rep was included as a random effect nested within the site: *Diversity ∼ Marker * Site * AgeGroup + (1 |Site:Core_Rep).* This framework allowed variability among PCR replicates to be distinguished from broader variation among sediment cores.

Shannon diversity was modelled using linear mixed models (lmer), whereas richness, treated as count data, was modelled using negative binomial mixed models (glmer.nb) to account for overdispersion. Unlike Gaussian mixed models, negative binomial models do not estimate residual variance as a separate variance component because variability is represented through the mean–variance relationship and dispersion parameter of the distribution. Post hoc comparisons were made using emmeans with Tukey correction.

Beta diversity was analysed using Bray–Curtis dissimilarity matrices from relative abundance data, calculated using the vegdist function of the *vegan* package (v2.7-2) in R, and visualised with non-metric multidimensional scaling (NMDS) based on community profiles averaged across PCR replicates to reduce stochastic variation associated with amplification and better represent the underlying biological signal. This approach is commonly used in *seda*DNA studies, where technical replication is used to minimise analytical noise prior to ecological interpretation. In addition, complementary analyses using non-averaged data quantified technical variability through within-sample dissimilarities and dispersion analyses (betadisper). These complementary analyses were not intended to infer ecological structure but to quantify the relative contribution of PCR-level stochasticity and among core ecological variability.

Distance-based redundancy analysis (db-RDA) was performed using Bray–Curtis dissimilarity matrices calculated from relative abundance data, consistent with beta diversity analyses, with the site and sediment age group as constraints (capscale, *vegan*). Taxonomic composition was summarised by plotting read relative abundance of major taxa (i.e., kingdom and phyla) using stacked barplots. The community composition was further tested using PERMANOVA (adonis2, 999 permutations), including site identity, sediment age group, and biological replicate as factors. Marginal sums of squares were used (by = “margin”) so that each term was evaluated after accounting for all other terms in the model. The homogeneity of the dispersion was assessed using betadisper and permutest. The dispersion was evaluated separately for each factor and marker. The proportion of variance explained by each factor was visualised using bar graphs representing the percentage of explained variation associated with each factor for each marker gene.

A joint species distribution model (jSDM) was fitted to partition community variation into fixed effects (site identity, sediment age group) and random effect (sediment core). The model was implemented using the Hierarchical Modelling of Species Communities (HMSC) framework with the *Hmsc* package in R (Tikhonov et al., 2020). Models were fitted separately for four datasets: complete eukaryotic COI, complete eukaryotic 18S, metazoan-only COI, and metazoan-only 18S. For each dataset, the input consisted of a phylum-level presence– absence matrix derived from the relative-abundance table after technical PCR replicates had been collapsed at the sediment-sample level. Presence–absence values were generated by converting all positive relative-abundance values to 1 and absences to 0. Site identity, sediment age group, and their interaction were included as predictors. Random effects were specified for site and core identity, with core identity defined as the interaction between site and core replicate. Model performance was evaluated using AUC, RMSE, and Tjur’s R². The relative contribution of fixed and random effects was assessed by variance partitioning, allowing quantification of the proportion of explained variation attributable to site, sediment age group and sampling structure. All jSDM outputs are provided in the Supplementary Material.

Data visualisation was performed using *ggplot2* (Wickham, 2011), and data handling was performed using *phyloseq* (McMurdie & Holmes, 2013). Maps were generated using the *marmap* package v1.1.10 (Pante & Simon-Bouhet, 2013).

## 3. Results

### 3.1. Dating and geochemical correlation

Radiometric dating based on excess or unsupported ^210^Pb activities indicated recent sediment deposition at all sites, although accumulation rates and temporal resolution differed markedly between them (Supplementary Data 1). ^210^Pb declined to supported levels at ∼49 cm (CLR), 48 cm (CCO), and ∼10 cm (STO), defining the depth range over which recent sediment accumulation rates were estimated. Extrapolation of these accumulation rates indicated that the analysed horizons span approximately the last 500 years. The estimated sediment accumulation rates were 0.63 cm yr^-1^ at CLR, 0.53 cm yr^-1^ at CCO, and substantially lower at STO (∼0.1 cm yr^-1^) (Table S3; Fig. S1). The slower accumulation rate observed at STO is consistent with the restricted occurrence of unsupported ^210^Pb in the uppermost part of the core, reflecting a condensed sediment sequence.

On the basis of these rates, sampled horizons represent two depositional periods. Surface sediments corresponded to recent decades (1974 at STO; 2014 at CCO; 2017 at CLR), while deeper layers corresponded to older deposits. However, the absolute ages differed between sites: CCO and CLR corresponded to the mid-20th century (1939 and 1952), while STO represented a much older interval (∼16^th^ century CE, based on extrapolation of ^210^Pb data). Therefore, ’old’ sediments reflect site-specific depositional histories rather than equivalent time periods.

Standardised XRF profiles revealed consistent stratigraphic patterns among biological replicates within each site (Fig. S2; Supplementary Data 2). Pairwise correlations between replicate cores showed generally moderate to strong agreement for lithogenic and detrital elements (Ti, Rb, Zr) (Table S4), particularly at CLR where the median r values were approximately 0.75 or higher for Rb, Ti and Zr. At CCO, Rb showed strong coherence, whereas Fe was more variable. In STO, Ti showed strong agreement (median r = 0.74), while Fe and Zr were more variable.

### 3.2. Sequencing output and data overview

High-throughput sequencing yielded 101,370 ASVs for COI and 57,415 for 18S. After quality filtering and taxonomic curation, 65,466 (COI) and 51,263 (18S) ASVs were retained. The mean sequencing depth per PCR replicate was 226,613 reads (SD = 309,124) for COI and 468,673 (SD = 801,189) for 18S. Rarefaction curves and sequencing-depth diagnostics are provided in the Supplementary Material (Fig. S3). Coverage-based analyses using iNEXT indicated that sequencing effort was sufficient across all PCR replicates for both markers. Sample coverage was consistently close to 1, and estimated asymptotic richness was identical to observed richness in all PCR replicates, indicating that sequencing depth was sufficient to recover the detectable diversity within individual PCR replicates despite differences in sequencing depth (Fig. S3). Additional sensitivity analyses based on sample-size rarefaction yielded highly consistent results. Although rarefaction reduced absolute richness values, particularly for COI, the overall patterns among PCR replicates and the outcomes of the mixed-effects richness models remained consistent with the original analyses, indicating that the biological conclusions were robust to sequencing-depth variation (Fig. S13).

Negative controls showed zero or minimal read counts (mean ± SD: 1,909 ± 3,085 for COI and 11,773 ± 25,102 for 18S). Of the 22 negative controls analysed per marker, 10 (COI) and 9 (18S) yielded zero reads.

### 3.3. Taxonomic composition

The taxonomic composition was broadly similar across sites in terms of the major taxonomic groups recovered (Fig. 2). Both COI and 18S recovered Metazoa, Plantae, Protista, and Fungi, although their relative abundance profiles differed substantially between markers. COI was characterised by a higher proportion of unassigned ASVs, whereas 18S recovered a broader range of assigned metazoan and non-metazoan taxa. Absolute read-count summaries supporting these patterns are provided in Table S5. Among the major taxonomic groups recovered, Protista represented the largest component of the 18S dataset (5,070,100 reads; 40.55%), followed by unassigned sequences (3,737,679 reads; 29.90%), Fungi (1,564,356 reads; 12.51%), Metazoa (1,088,146 reads; 8.70%), and Plantae (1,042,361 reads; 8.34%). In contrast, COI showed a higher proportion of unassigned sequences (435,752 reads; 53.08%), followed by Metazoa (200,713 reads; 24.45%), Protista (93,794 reads; 11.43%), Fungi (81,266 reads; 9.90%), and Plantae (9,350 reads; 1.14%). Surface layers (4–5 cm) showed higher proportions of metazoan and protistan ASVs, while deeper layers (45–50 cm) were relatively enriched in plants.

**Fig. 2.**
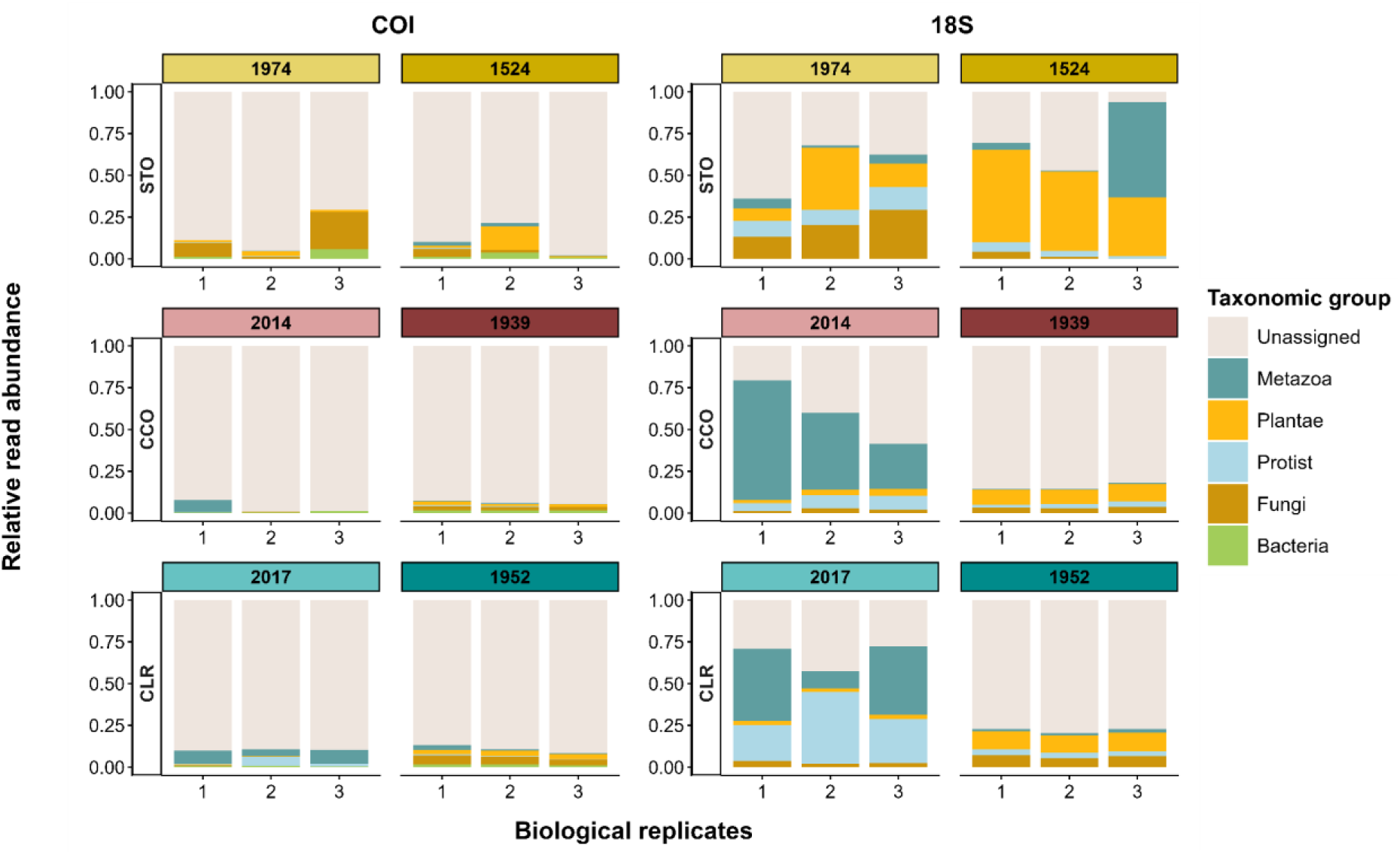
Relative abundance of major taxonomic groups recovered across the three intertidal sites (STO, CCO and CLR; see text) based on metabarcoding-based sedaDNA metabarcoding. Bars represent the mean relative abundance of ASVs per biological replicate (sediment core), obtained after combining the eight technical PCR replicates. Facets indicate sediment age group and genetic marker (cytochrome c oxidase I [COI] and the V7 region of 18S rRNA).

Biological replicates within each site displayed similar kingdom-level profiles, indicating low intrasite variability relative to differences among sites and sediment age groups, with the exception of STO, where variability was higher.

When the analysis was restricted to Metazoa, taxonomic composition remained broadly consistent across both sites and sediment age groups (Fig. S4), and composition was dominated by Annelida, Nematoda, Arthropoda and Mollusca. However, the relative abundances of these major phyla varied among biological replicates in several samples, indicating some degree of heterogeneity (Fig. S4).

### 3.4. Alpha diversity

Technical-replicate variability was greater for ASV richness than for Shannon diversity (Table S6; Fig. 3). Shannon diversity showed moderate to high repeatability (ICC = 0.43–0.69), while ASV richness showed consistently lower repeatability (ICC = 0.18–0.31), indicating greater variability between technical replicates. This pattern was consistent across markers and taxonomic subsets. Variability among PCR replicates primarily reflected random variation among the eight PCR replicates generated from the same sediment sample rather than consistent systematic shifts among replicates. For richness, the model did not directly estimate residual technical variance because richness was modelled using a negative binomial distribution; however, visual inspection of Fig. 3 indicated appreciable variability among technical replicates within some samples.

**Fig. 3.**
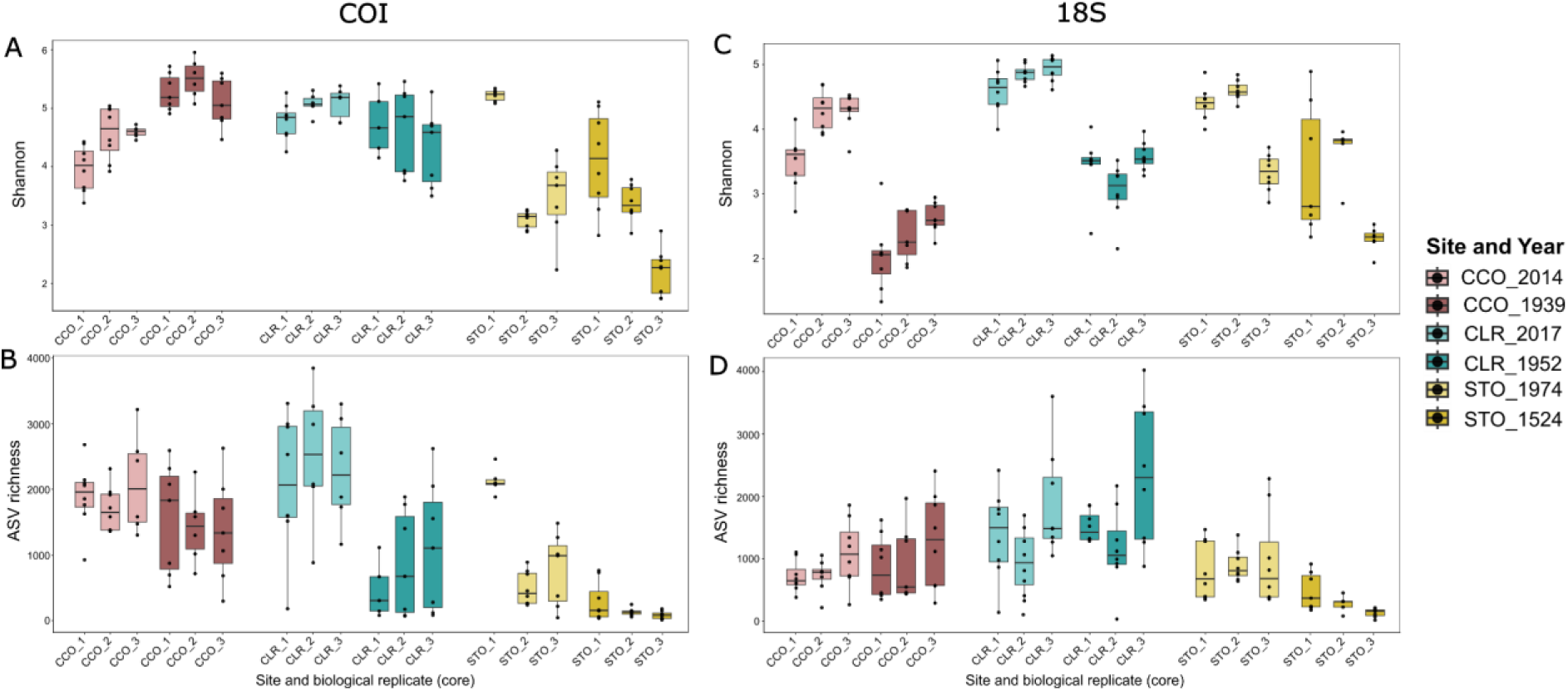
Boxplots showing variation in (A) Shannon diversity for COI, (B) ASV richness for COI, (C) Shannon diversity for 18S, and (D) ASV richness for 18S among biological replicates (cores) across sites and sediment age groups. Each box represents the distribution of technical PCR replicates for a given biological replicate and sediment age group, overlaid with individual observations. Colours indicate site and sediment age group, with lighter shades representing recent sediments and darker shades representing older sediments.

After accounting for technical variability, core-level biological replication had a limited effect on alpha diversity in the entire eukaryotic community (Table S7; Fig. 3), with significant among-core differences concentrated mainly at STO. This pattern depended on the diversity metric. Shannon diversity for both genetic markers showed no differences between biological replicates within CCO and CLR, whereas significant differences were detected at STO in both recent and old sediments. ASV richness showed even fewer significant contrasts among biological replicates for both markers, with only isolated differences detected at STO. These results indicate generally low within-site heterogeneity among biological replicates, with localised variability at STO, particularly for Shannon diversity. In contrast, site-level differences were more pronounced (Table S9; Figs. 2–4). For COI, Shannon diversity differed between sites only in old sediments, where model estimates indicated significantly higher Shannon diversity at STO than at CCO and CLR (Table S9). ASV richness showed a clearer spatial structure, with STO consistently exhibiting a higher richness than the other sites. For 18S, the site effects were weaker for Shannon diversity but remained evident for richness in old sediments (Table S9). Overall, alpha diversity was more strongly structured by site identity than by biological replication, particularly for richness.

**Fig. 4.**
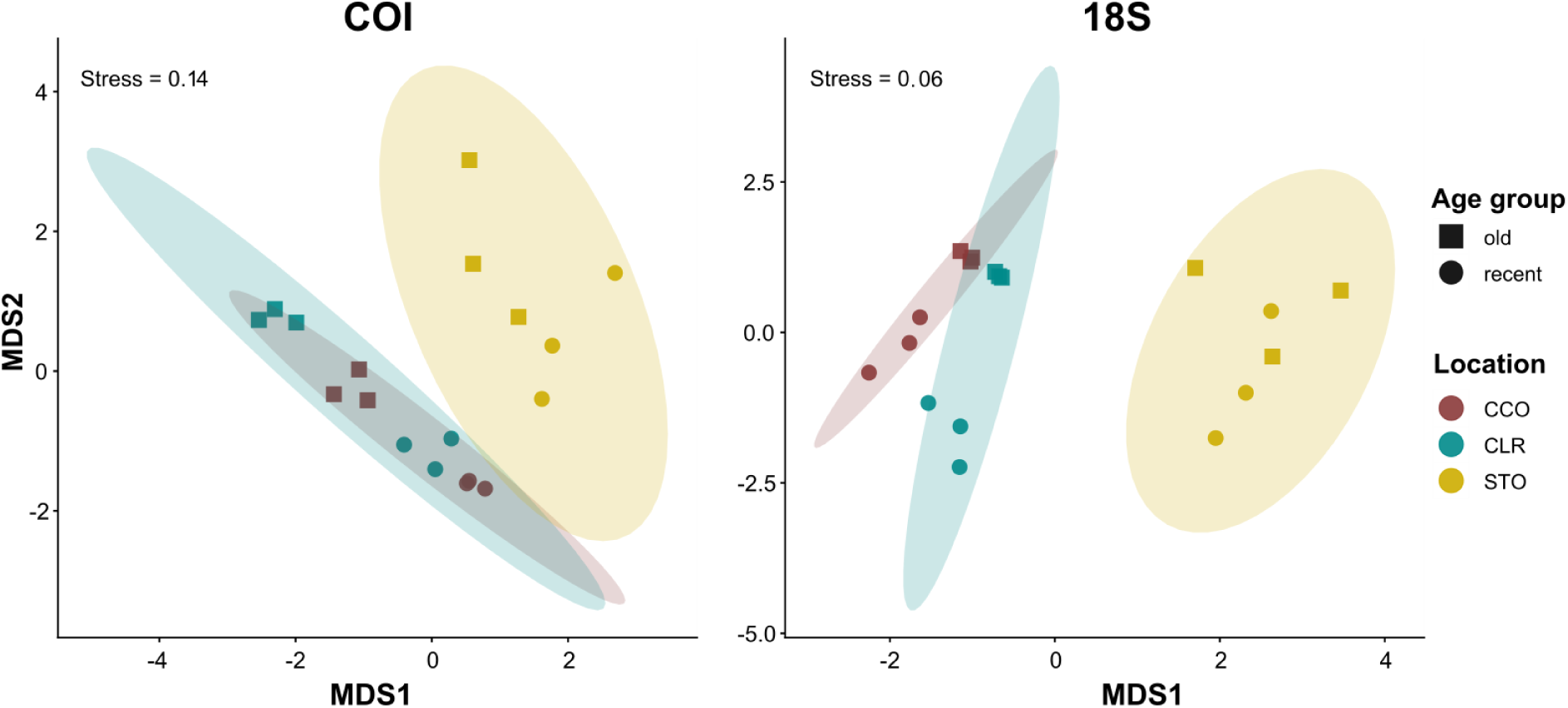
NMDS ordination of eukaryotic communities. Non-metric multidimensional scaling (NMDS) ordination of COI- and 18S-based metabarcoding-based sedaDNA eukaryotic community composition across the three intertidal sites. The plot shows axes 1 and 2 of the two-dimensional NMDS solution based on Bray–Curtis dissimilarities of ASVs relative abundances (stress value indicated in each panel). Points are coloured by site (CCO, CLR and STO; see text) and shaped according to sediment age group (recent versus old), while shaded ellipses delimit the dispersion of biological replicates (cores) within each site.

Within the metazoan subset, biological replication had little influence on alpha diversity, with significant differences again mostly restricted to STO (Table S8; Fig. S5). Shannon diversity based on 18S showed occasional differences among the cores in STO, whereas the richness remained largely consistent among cores for both markers, apart from two significant 18S contrasts at STO. The effects at the site were more consistent (Table S10; Figs. S4–S5), with STO generally showing higher diversity than CCO and CLR for both markers, especially for richness. These patterns were consistent with those observed in the entire eukaryotic dataset, indicating that the limited effect of biological replication and the stronger influence of the site are robust across taxonomic subsets.

### 3.5. Community structure and beta diversity

NMDS analyses revealed a pronounced spatial and stratigraphic structure of the metabarcoding-based *seda*DNA communities for both the COI and the 18S markers (Fig. 4). Within each site, biological replicates were clustered tightly with minimal dispersion, demonstrating highly similar community composition among cores and generally low within-site heterogeneity relative to broader community patterns (Fig. 4). Across both markers, community composition showed site-related structure, mainly driven by the separation of STO from CCO and CLR, whereas the latter partially overlapped. Sediment age group represented a secondary gradient, partially separating recent and old samples within sites (Fig. 4).

To evaluate the relative contribution of technical and biological variation, additional analyses were performed that included non-collapsed technical replicates (Figs. S6–S7). Analyses including individual PCR replicates (Fig. S6) showed greater dispersion than averaged datasets, but this variation remained largely confined within the clusters defined by site and sediment age group. Bray–Curtis dissimilarities among PCR replicates derived from the same sediment sample were generally lower than, or comparable to, dissimilarities among biological replicates within sites. This pattern suggests that stochastic variation associated with PCR amplification contributes mainly to variability within sediment samples, whereas differences among cores contributed additional variability at the local scale (Fig. S7).

NMDS ordinations restricted to metazoan taxa had low stress values (COI: stress = 0.120; 18S: stress = 0.084) and showed a broadly similar but weaker community structure compared to the entire eukaryotic community (Fig. S8). For both marker datasets, site-level separation was reduced, with greater overlap between sites. Age-related patterns were also less pronounced, and metazoan samples exhibited a tighter clustering overall. Despite this, the biological replicates again clustered closely within the sites, indicating a consistent community composition among cores. The jSDM supported these patterns (Table S11; Fig. S9). The site explained the largest proportion of variance (63–77%), followed by the sediment age group (14–28%), while random effects associated with biological replication were consistently low (<6%) (Table S11). This pattern was consistent across markers and community subsets, confirming that spatial differences between sites dominate the community structure. Model performance was high (AUC: 0.929–0.959; Tjur’s R²: 0.220–0.277) (Table S11). Similar trends were observed at the phylum level (Fig. S10; Supplementary Data 3).

Our db-RDA further confirmed the significant effects of the site and sediment age group for both markers (Fig. S11). For COI, both the site (F = 3.33, p = 0.001) and the sediment age group (F = 3.86, p = 0.001) were significant and similar results were obtained for 18S (site: F = 5.01, p = 0.001; sediment age group: F = 5.18, p = 0.001). The constrained axes were significant for both markers (COI: all axes, p = 0.001; 18S: axes 1 and 2, p = 0.001, axis 3, p = 0.002), indicating that these effects were distributed over multiple dimensions of community variation. In agreement with NMDS, the effects of the site were stronger than the effects of age, and the differences between the biological replicates were minimal.

PERMANOVA analysis confirmed that community composition was primarily structured by site and sediment age group, with negligible contributions from biological replication (Table S12; Fig. 5). For COI, the site (R² = 0.272, p = 0.001) and the sediment age group (R² = 0.157, p = 0.001) were significant, while biological replication was not (R² = 0.032, p = 0.712). Similar results were obtained for 18S (site: R² = 0.345, p = 0.001; sediment age group: R² = 0.177, p = 0.002; biological replication: R² = 0.029, p = 0.516). When restricted to metazoans, biological replication remained non-significant for both markers, while the effects of the site and sediment age group were weaker or marker dependent (Table S12; Fig. S12).

**Fig. 5.**
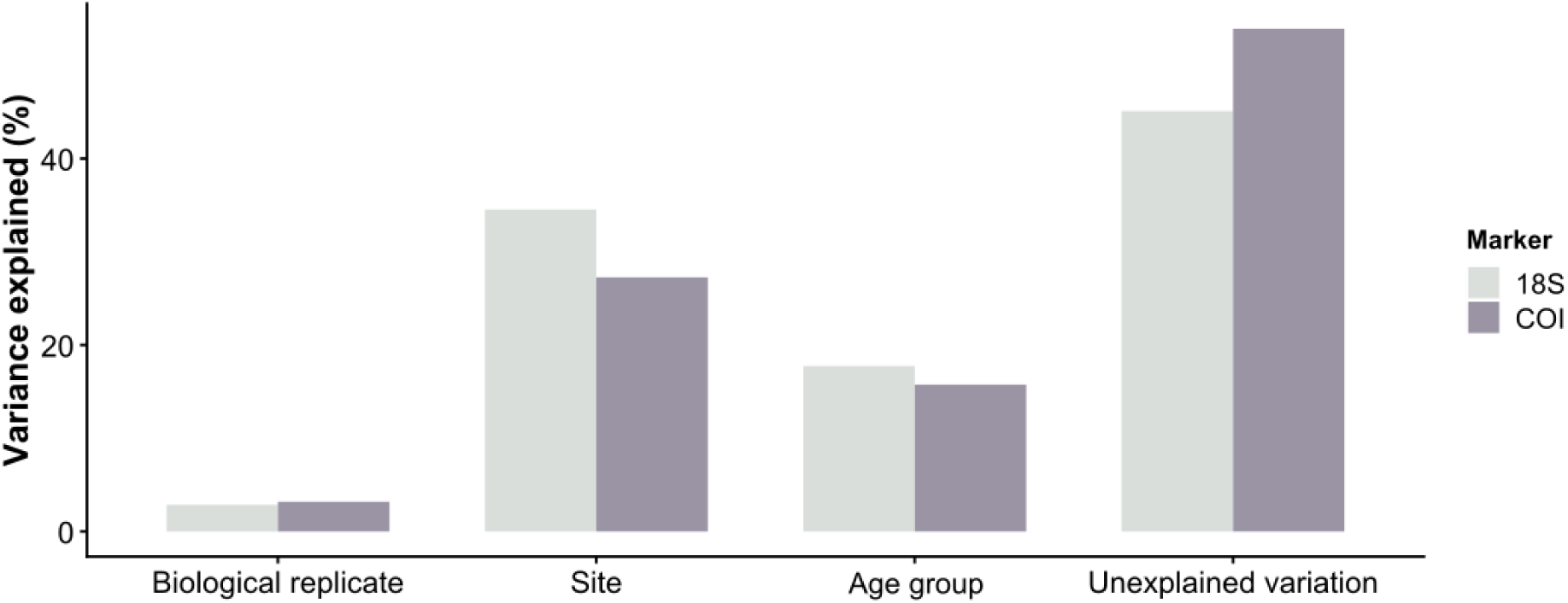
Bar plots showing marginal R² values from PERMANOVA models (adonis2, 999 permutations) illustrating the relative contribution of site, sediment age group and biological replication to Bray–Curtis dissimilarities in eukaryotic community composition for the COI and 18S datasets. Marginal R² values represent the unique contribution of each factor after accounting for the others and are intended for comparative interpretation among predictors rather than as additive components of the total explained variance. The unexplained variation corresponds to residual community dissimilarity not captured by the tested factors and may arise from unmeasured environmental gradients, fine-scale spatial heterogeneity, stochastic biological processes, temporal integration of sediments and methodological noise inherent to metabarcoding-based sedaDNA.

The homogeneity tests for the multivariate dispersion indicated that, in the entire eukaryotic dataset, part of the PERMANOVA signal may reflect differences in dispersion (Table S13). In contrast, no dispersion differences were detected in the metazoan subset, suggesting that the PERMANOVA results in this subset reflect differences in detected community composition rather than differences in dispersion.

## 4. Discussion

Across all sampling sites (CCO, CLR and STO) and genetic markers (COI and 18S), core-level biological replication contributed relatively little to variation in detected metabarcoding-based *seda*DNA community composition compared with site identity and sediment age group. This suggests that, within the sampled sites, replicate sediment cores captured broadly coherent stratigraphic community signals, whereas among-site and stratigraphic differences represented the dominant sources of ecological variation. This pattern was supported by PERMANOVA (Table S12; Fig. 5), where biological replicates explained a small and non-significant fraction of the variation, while site identity and, in most cases, sediment age group accounted for substantially larger proportions of variance. NMDS ordinations supported this result, with biological replicates clustering tightly within sites in the NMDS space (Fig. 4; Fig. S8), indicating high similarity among cores. Although some significant differences among cores were detected in alpha diversity (Table S7; Fig. 3), these effects were largely localised at STO and were less consistent than broader site- and age-related patterns.

This limited contribution of biological replication should be interpreted in the context of metabarcoding analyses in which diversity estimates were derived from multiple PCR replicates per sample and technical variation was explicitly accounted for, allowing variability within sediment samples to be distinguished from differences among sediment cores (Bulcke et al., 2021; Shirazi et al., 2021). This is particularly relevant for richness, which showed low repeatability across PCR replicates (Table S6), potentially reflecting stochastic detection of low abundance taxa. Rarefaction and coverage analyses indicated that sequencing effort was generally sufficient to approach asymptotic diversity across samples (Fig. S3). Repeating the richness analyses after sequencing-depth standardisation produced the same overall biological interpretation, indicating that the observed patterns were robust to variation in sequencing depth. Although sequencing depth contributed to absolute richness estimates, it did not alter the relative ecological patterns or the conclusions regarding the effects of site, sediment age, and technical replication. However, agreement between observed and rarefied richness (Fig. S13) was higher for the 18S marker (ρ = 0.967) than for COI (ρ = 0.741), reflecting a greater effect of sequencing-depth standardisation on COI richness estimates. Nevertheless, this difference did not affect the overall ecological interpretation, as the same relative effects of site, sediment age, and technical replication were recovered following sequencing-depth standardisation. Within this context, our results indicate that metabarcoding-based *seda*DNA signals are highly consistent among nearby cores and are primarily structured by broader spatial and stratigraphic gradients. Some aquatic eDNA studies have reported strong local heterogeneity and pronounced fine-scale spatial variation (Lamy et al., 2021), whereas others have identified stronger site-level structuring with limited variability among nearby replicates (Beentjes et al., 2019). Our results are more consistent with the latter pattern and may reflect the integrative nature of sedimentary DNA archives, where ecological signals are accumulated and mixed through sediment deposition, resuspension, and other post-depositional processes, while still preserving sufficient stratigraphic structure to resolve temporal ecological patterns (Capo et al., 2021; Nguyen et al., 2023; Pearman et al., 2021).

### 4.1. Technical replication and metric-specific sensitivity

The influence of technical replication on metabarcoding-based diversity estimates depended primarily on the diversity metric considered rather than on the genetic marker itself, although richness estimates derived from COI were somewhat more affected by sequencing depth than those from 18S (Clarke et al., 2017; van der Loos & Nijland, 2021). Although overall alpha-diversity estimates differed between genetic markers, the contrasting responses of richness and Shannon diversity indicate that metric choice had a greater influence on sensitivity to technical replication than marker choice. Technical replication had a much stronger effect on ASV richness than on Shannon diversity, consistently across both the complete eukaryotic community and the metazoan subset (Table S6). This pattern reflected stochastic variation within samples rather than systematic differences among PCR replicates, likely driven by inconsistent detection of rare taxa. Consequently, richness showed low repeatability across PCR replicates, consistent with its sensitivity to stochastic amplification and sequencing effects affecting low-abundance taxa (Shirazi et al., 2021). In contrast, Shannon diversity showed moderate to high repeatability and was generally robust to technical replication, indicating more consistent estimates among PCR replicates. This suggests that abundance-weighted metrics are more resilient to stochastic detection of rare ASVs but may be influenced by biases associated with primer performance, template competition, and amplification efficiency (Casey et al., 2021; Giebner et al., 2020; Leite et al., 2021; Zinger et al., 2019).

Overall, alpha diversity estimates are not uniformly robust to technical replication, but depend strongly on the diversity metric considered: Shannon diversity was comparatively stable, while richness was more sensitive to technical variability and should be interpreted with greater caution (Table S6).

### 4.2. Biological replication and small-scale lateral heterogeneity

Strong stratigraphic agreement among spatially separated sediment cores was supported by geochemical data, providing external validation of lateral continuity within sites (Hennekam & de Lange, 2012; Kern et al., 2019). Standardised XRF profiles showed parallel depth trends across biological replicates (Fig. S2), particularly for lithogenic elements (Ti, Rb, Zr), commonly used as indicators of sediment provenance (Croudace et al., 2019; Rothwell & Croudace, 2015). Minor discrepancies were observed in redox-sensitive elements (e.g. Fe), but they probably reflect small-scale and variable diagenetic effects rather than stratigraphic offsets (Beam et al., 2018; Tribovillard et al., 2006). Overall, the geochemical consistency supports the interpretation that low biological variance observed among cores is unlikely to be driven primarily by stratigraphic mismatch or sampling artefacts.

Consistent with this framework, core-level biological replication had limited effects on alpha diversity overall, with localised heterogeneity observed mainly at STO (Table S7; Fig. 3). This pattern suggests that, although metabarcoding-based *seda*DNA signals were generally coherent within sites, local environmental conditions may still influence small-scale variability in detected community composition. Most pairwise comparisons were non-significant, indicating a high degree of lateral consistency within sites. Deviations were mainly observed at STO, where differences between cores were more frequent, particularly for Shannon diversity and, to a lesser extent, richness. This pattern may reflect locally elevated heterogeneity (Bowen et al., 2012; Hestetun et al., 2021), potentially related to the relatively low sediment accumulation rate observed at this site, which could amplify small-scale spatial variability due to reduced and more heterogeneous sediment inputs. However, more work is needed to confirm this interpretation.

Patterns from beta diversity analyses suggest that metabarcoding-based *seda*DNA assemblages were largely laterally consistent at the spatial scale examined (∼50 cm), with only localised deviations from this pattern. This interpretation was supported by PERMANOVA analyses, which indicated that biological replication explained relatively little variation in community composition compared with site identity and sediment age group (Fig. 5; Table S12). Similarly, ordination approaches (NMDS and db-RDA) showed strong clustering of samples within sites and broader separation associated with site identity and sediment age group (Fig. 4; Fig. S11). Under comparable environmental conditions, these findings suggest that increasing the number of closely spaced replicate cores within sites may provide limited additional information relative to other components of sampling design. Instead, greater gains in ecological resolution may be achieved by prioritising broader spatial coverage across sites, increased stratigraphic resolution, and sufficient PCR replication to account for technical variability. Nevertheless, localised heterogeneity observed at STO indicates that some level of within-site biological replication may still be valuable in environmentally heterogeneous settings.

This pattern suggests that DNA recovered from sedimentary archives integrates ecological signals beyond the scale of individual cores. Such spatial averaging aligns with previous work (Pearman et al., 2021) showing that *seda*DNA records reflect temporally and spatially integrated signals influenced by sediment mixing, lateral transport, and post-depositional molecular movement (Giguet-Covex et al., 2019; Harrison et al., 2019; Parducci et al., 2017; Turner et al., 2015). However, this interpretation depends on adequate control of analytical uncertainty. Under limited technical replication, stochastic amplification can inflate apparent differences between cores, particularly for metrics sensitive to rare taxa (Fig. S6–S7). Analyses including non-collapsed PCR replicates showed increased within-sample dispersion, whereas averaging across replicates reduced this technical variability in beta diversity space and clarified the tight clustering of biological replicates observed in the main analyses. In general, small-scale biological heterogeneity was detectable but became a minor component of variation once analytical noise was accounted for.

### 4.3. Ecological drivers of the structure of the metabarcoding-based *seda*DNA community: dominance of the site and sediment age group

Given the limited contribution of within-site variability, metabarcoding-based *seda*DNA communities appeared to be structured primarily by broader spatial and stratigraphic gradients. Across all analyses, site identity explained a larger proportion of community variation than biological replication, while sediment age group represented a secondary but consistent source of variation (Table S11; Table S12; Fig. 4; Fig. S9).

Site-level differences likely reflect a combination of environmental and depositional characteristics specific to each location. Differences among sites may arise from local habitat characteristics, surrounding vegetation, sediment source, hydrodynamic conditions, and connectivity with adjacent environments, all of which can influence both the origin of DNA entering the sediment and its subsequent accumulation and preservation. Such processes can generate distinct site-level community signatures even across relatively short spatial distances.

Sediment age group represented a secondary but consistent gradient separating recent and older samples within sites (Table S12; Fig. S11). This vertical pattern may reflect ecological turnover through time, but could also result from stratigraphic filtering and age-dependent differences in DNA preservation and persistence within sediments. Metabarcoding-based *seda*DNA records integrate both biological and depositional processes, making it likely that the observed vertical structure reflects a combination of ecological change and taphonomic influences rather than ecological turnover alone (Capo et al., 2021; Parducci et al., 2017; Pedersen et al., 2016).

The consistency of these patterns across both COI and 18S, despite differences in taxonomic scope and amplification behaviour, suggests that the observed gradients likely reflect broader environmental structure rather than marker-specific amplification artefacts (Atienza et al., 2020; Giebner et al., 2020). Together, these findings indicate that metabarcoding-based *seda*DNA variation in the studied system was primarily structured by environmental and stratigraphic context, whereas variation among nearby sediment cores represented only a minor component of total community variation. This is consistent with previous studies using DNA recovered from sedimentary archives, which have reported stronger structuring by depositional setting and temporal position than by fine-scale local variability (Pearman et al., 2021).

### 4.4. Implications and limitations of replication in metabarcoding-based *seda*DNA study design

Our results indicate that, under stratigraphically coherent conditions, the benefits of increasing biological replication at very local scales diminish once analytical variability is adequately controlled. In contrast, some degree of technical replication is necessary to reduce within-sample variability, particularly for richness-based metrics, and to distinguish PCR-level stochasticity from ecological differences among samples. An additional limitation is that technical replication was evaluated only at the PCR level. Other analytical sources of variation, including independent DNA extraction replicates, subsampling within sediment layers (i.e., different portions of the same stratigraphic horizon), and sequencing runs, were not explicitly evaluated and may also contribute to uncertainty in metabarcoding-based *seda*DNA studies. However, this does not imply that biological replication can be omitted. Without within-site replication, it is not possible to quantify local variability or distinguish it from differences among sites, limiting ecological inference at broader spatial scales (Hurlbert, 1984; Marshall, 2024). In this sense, the main implication of our study is not that biological replication is unnecessary, but rather that its value depends on the spatial scale of the question and the sedimentological context.

Under stratigraphically coherent conditions, a limited number of sediment cores per site may be sufficient to capture local ecological structure, although this depends on the sampling location within the depositional environment. Sedimentary DNA signals can vary across sediment surfaces, with more representative and spatially integrated signals often associated with depositional centres (Giguet-Covex et al., 2019). In our study, the lower heterogeneity observed among the cores at CCO and CLR compared to STO is consistent with this pattern, as these sites showed higher sediment accumulation rates and therefore likely greater signal integration. This highlights that core placement, rather than replication alone, is a key determinant of local species richness. In well-preserved sedimentary archives, prioritising stratigraphic resolution, spatial coverage, or analytical robustness may therefore yield greater gains than increasing local replication (Heintzman et al., 2023). In systems with higher spatial heterogeneity, increased biological replication may still be beneficial. In some depositional settings, however, the processes generating local heterogeneity (e.g. sediment mixing, hydrodynamic disturbance, or variable sediment accumulation) may also complicate chronological control and palaeoecological interpretation.

In our study, the temporal resolution of ’old’ sediment layers differed between sites due to contrasting accumulation rates, with STO representing a substantially older depositional period than CCO and CLR. This was considered when interpreting temporal patterns, as age-group differences partially reflected absolute age differences. Finally, the patterns reported here are scale- and context-dependent. The observed lateral consistency may not apply to other systems with higher disturbance, complex hydrodynamics, or greater variability in the microhabitat. Future studies should explore how sedimentary processes and spatial scale influence optimal replication strategies in other environments.

### 4.5. Conclusions

Our study shows that, under stratigraphically coherent conditions, biological replication contributed comparatively little to metabarcoding-based *seda*DNA analyses. Across multiple intertidal wetland sites, spatially separated sediment cores yielded consistent patterns of alpha diversity and community structure, with only minor local deviations. In contrast, site identity and sediment age group were the dominant drivers of variation, as consistently demonstrated in ordination, PERMANOVA, and hierarchical modelling approaches. These findings indicate that, when major sources of analytical variability are explicitly accounted for, research effort may be more effectively directed towards spatial coverage, stratigraphic resolution, sequencing depth, and technical replication than increasing biological replication within sites. However, biological replication in the form of multiple sediment cores should not be omitted, as a minimum level of within-site replication remains necessary to quantify local variability and support robust comparisons among sites. More broadly, our findings highlight the need to align replication strategies with the sedimentological context and study objectives, while explicitly accounting for multiple sources of variability, including analytical stochasticity and spatial heterogeneity, in metabarcoding studies using DNA recovered from sedimentary archives.

## Acknowledgments

This work was supported by the TEMPOINVASIONS grants TED2021-132228B-C21 and TED2021-132228B-C22, both funded by MCIN/AEI/10.13039/501100011033 and the European Union (“NextGenerationEU”/PRTR). We are also grateful to the Spanish Government for providing financial support with the BlueDNA (PID2023-146307OB) grant (MICIU/AEI/10.13039/501100011033 and ERDF/EU ). Pawel Gaca and Madeleine Cobbold at GAU-Radioanalytical, University of Southampton, are thanked for conducting ^210^Pb dating analysis. Marc Rius and Xavier Turon are members of the research group SGR2021-00405 and Sandra Nogué is member of the research group SGR 01333, both funded by the Generalitat de Catalunya. Luke E. Holman was supported by the European Research Council through the SeaChange project (Grant agreement No. 856488).

## Data availability statement

The raw sedimentary DNA metabarcoding data can be found at the NCBI Sequence Read Archive under BioProject PRJNA1443734. Raw data, datasets, supplementary data, scripts and analytical workflows used in this study are publicly available at GitHub under DOI https://doi.org/10.5281/zenodo.21283114.

## Benefit-sharing statement

This study did not involve the use of genetic resources subject to access and benefit-sharing regulations, nor did it involve Indigenous peoples or local communities. Therefore, no specific benefit-sharing agreements were required.

## Author contributions

Elena Baños, Marc Rius, Xavier Turon, Andrew B. Cundy and Erik de Boer conducted the fieldwork and collected the samples. Dating analyses were carried out by Andrew B. Cundy and Elena Baños and interpreted by Andrew B. Cundy, Sandra Nogué and Elena Baños. Elena Baños performed the laboratory work and conducted the bioinformatic analyses of metabarcoding datasets. XRF data were processed by Erik de Boer and analysed by Sandra Nogué and Elena Baños. Data analyses were conducted by Elena Baños and Clara Ras, and statistical interpretation was carried out by Elena Baños and Luke E. Holman. Elena Baños wrote the manuscript with major contributions from Marc Rius and Luke E. Holman. All authors contributed to manuscript revision and approved the final version.

## Ethics statement

Ethical approval was not required for this study because it did not involve human participants, live animals, or protected species.

## Data availability statement

The raw metabarcoding-based *seda*DNA data can be found at the NCBI Sequence Read Archive under BioProject PRJNA1443734. Raw data, datasets, supplementary data, scripts and analytical workflows used in this study are publicly available at GitHub under DOI https://doi.org/10.5281/zenodo.21283114.

## Supplementary data

**Supplementary data 1:** Radiometric dating data, including raw ^²¹⁰^Pb measurements and age-depth model outputs (CF:CS model).

**Supplementary data 2:** X-ray fluorescence (XRF) data, including raw elemental counts and normalised geochemical profiles.

**Supplementary data 3:** Full statistical outputs from hierarchical modelling of species communities (jSDM), including model diagnostics and variance partitioning results.

## Supplementary tables

**Table S1:** Sampling locations and core characteristics for metabarcoding-based *seda*DNA analyses. Geographic coordinates (UTM), site names, and total core depths are reported for each sampling location.

| Core ID | Location | Latitude | Longitude | Core depth<br>(cm) |
| --- | --- | --- | --- | --- |
| STO | Santoña | 43°26'46.90"N | 3°27'58.62"W | 100 |
| CCO | Cicero | 43°25'01.13"N | 3°28'40.92"W | 50 |
| CLR | Colindres | 43°24'03.02"N | 3°27'14.80"W | 50 |

**Table S2:**
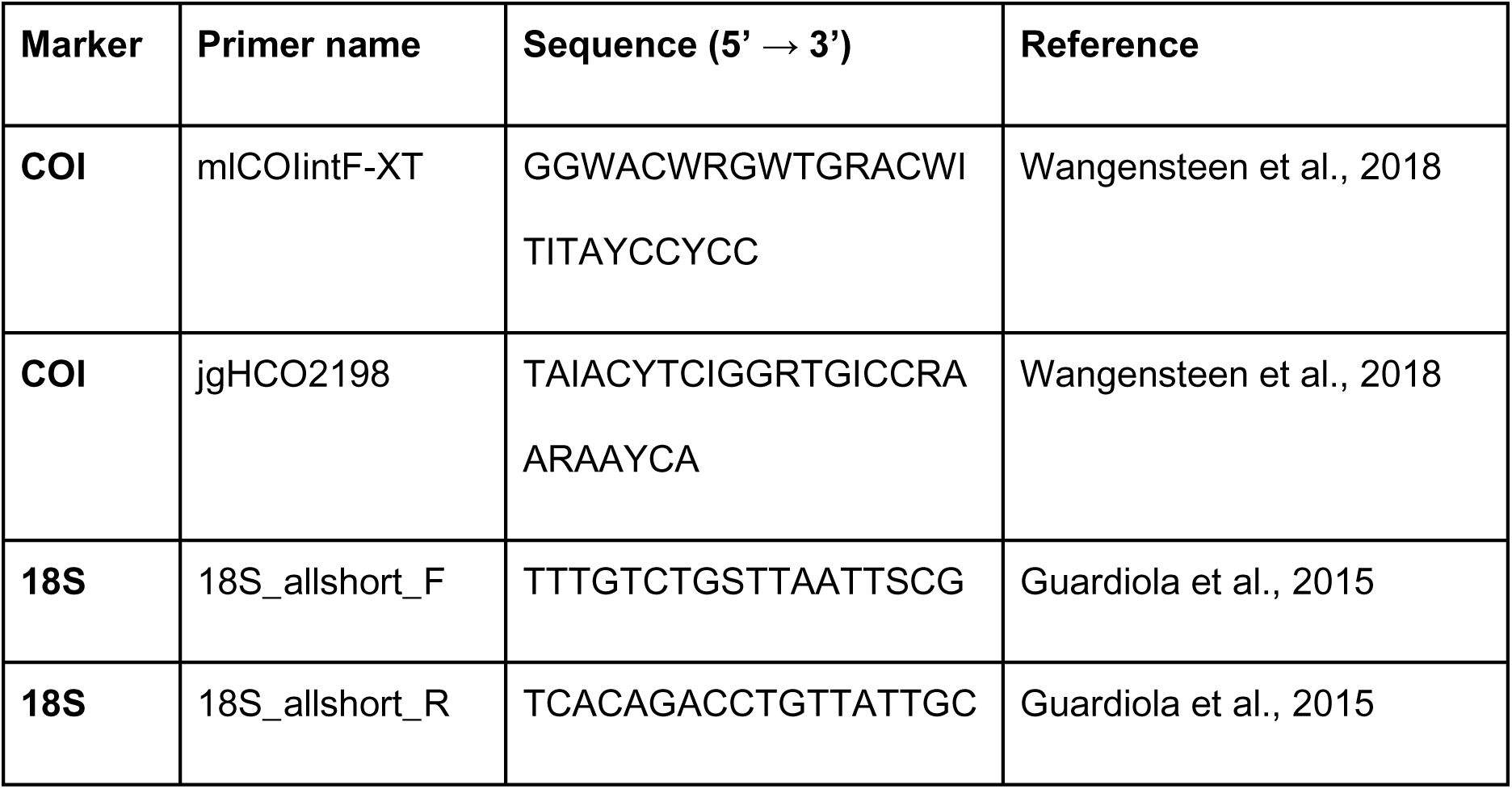
Primer sequences (5′–3′) used to amplify the COI (Leray-XT) and 18S (V7 region) markers in this study. Primers were dual-indexed using mirrored 8-bp tags to minimise tag-jumping artefacts.

**Table S3:** Sediment accumulation rates derived from excess ^210^Pb activities using the “Simple” Model of ^210^Pb dating (Robbins, 1978) for each sediment core. The table shows the estimated sediment accumulation rate (SAR) together with the associated 2σ uncertainty range derived from the regression of ln-transformed excess ^210^Pb activity against depth. The depth at which supported ^210^Pb activity is reached is also indicated.

| Core | Site | Depth of supported $^{210}\text{Pb}$ (cm) | SAR (cm yr<sup>-1</sup>) | $2\sigma$ range |
| --- | --- | --- | --- | --- |
| STO | Santoña | 10 | 0.1 | - |
| CCO | Cicero | 48 | 0.53 | 0.42 - 0.73 |
| CLR | Colindres | 49 | 0.63 | 0.50 - 0.87 |

**Table S4:** Pearson correlation coefficients (r) for selected lithogenic elements (Fe, Rb, Ti, Zr) between biological replicates within each site, calculated on standardised (z-scored) XRF profiles and restricted to depths shared between cores. Correlation strength is reported descriptively as weak (r < 0.30), moderate (0.30–0.59) and strong (≥ 0.60) to facilitate interpretation.

| Site | Element | Core1 vs.<br>Core2 (r) | Core1 vs.<br>Core3 (r) | Core2 vs.<br>Core3 (r) | Correlation |
| --- | --- | --- | --- | --- | --- |
| CCO | Fe | 0.293 | 0.038 | 0.005 | Weak |
| CCO | Rb | 0.724 | 0.626 | 0.855 | Strong |
| CCO | Ti | 0.429 | 0.411 | 0.615 | Moderate |
| CCO | Zr | 0.684 | 0.276 | 0.694 | Strong |
| CLR | Fe | 0.661 | 0.558 | 0.800 | Strong |
| CLR | Rb | 0.954 | 0.856 | 0.968 | Strong |
| CLR | Ti | 0.831 | 0.756 | 0.867 | Strong |
| CLR | Zr | 0.747 | 0.264 | 0.836 | Strong |
| STO | Fe | 0.763 | -0.135 | 0.064 | Weak |
| STO | Rb | 0.286 | 0.425 | 0.848 | Moderate |
| STO | Ti | 0.737 | 0.736 | 0.844 | Strong |
| STO | Zr | 0.488 | 0.346 | 0.824 | Moderate |

**Table S5:** Absolute read counts and relative proportions of major taxonomic groups recovered across COI and 18S metabarcoding datasets. Read counts are presented prior to relative-abundance transformation and summarised across all retained sediment samples. Relative proportions indicate the contribution of each major taxonomic group within each marker dataset.

| Marker | Kingdom | Total read counts | Percentage |
| --- | --- | --- | --- |
| COI | Unassigned | 435752 | 53.08 |
|  | Metazoa | 200713 | 24.45 |
|  | Protist | 93794 | 11.43 |
|  | Fungi | 81265 | 9.90 |
|  | Plantae | 9350 | 1.14 |
| 18S | Unassigned | 3737679 | 29.9 |
|  | Metazoa | 1088146 | 8.70 |
|  | Protist | 5070100 | 40.55 |
|  | Fungi | 1564356 | 12.51 |
|  | Plantae | 1042360 | 8.34 |

**Table S6:**
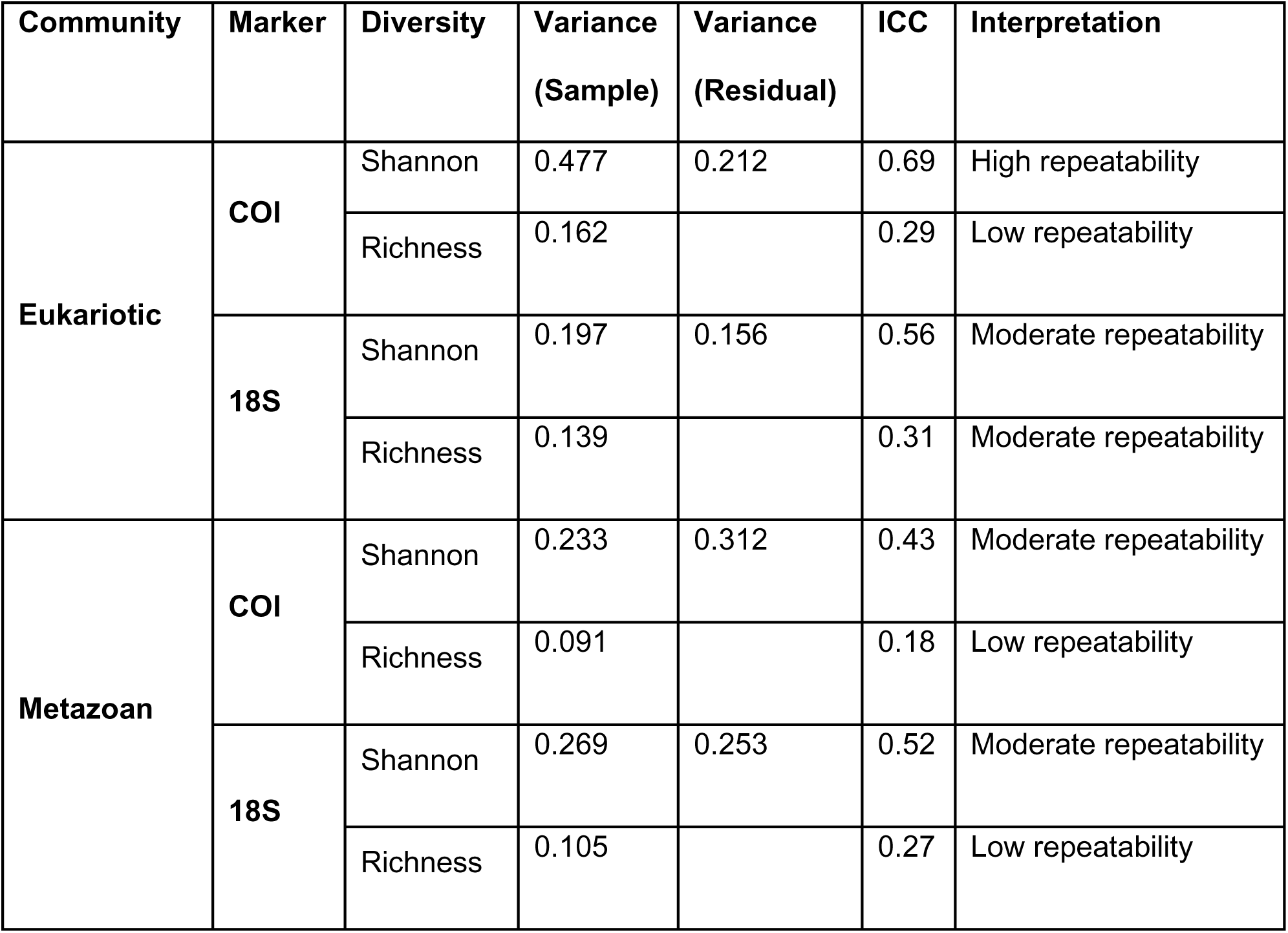
Variance components and intraclass correlation coefficients (ICC) from mixed-effects models evaluating repeatability among PCR replicates for richness and Shannon diversity. Site and sediment age group were included as fixed effects, while sample identity (Site:Core_Rep:AgeGroup) was included as a random effect. Variance (Sample) corresponds to variance attributable to differences among samples. For Shannon diversity models (lmer), residual variance represents within-sample variability among PCR replicates. For richness models (glmer.nb), residual variance is not reported because negative binomial models do not estimate a separate residual variance component. ICC values were interpreted as follows: >0.6 (high repeatability, low relative technical variation), 0.3–0.6 (moderate repeatability, noticeable technical variation), and <0.3 (low repeatability, substantial technical variation).

**Table S7:**
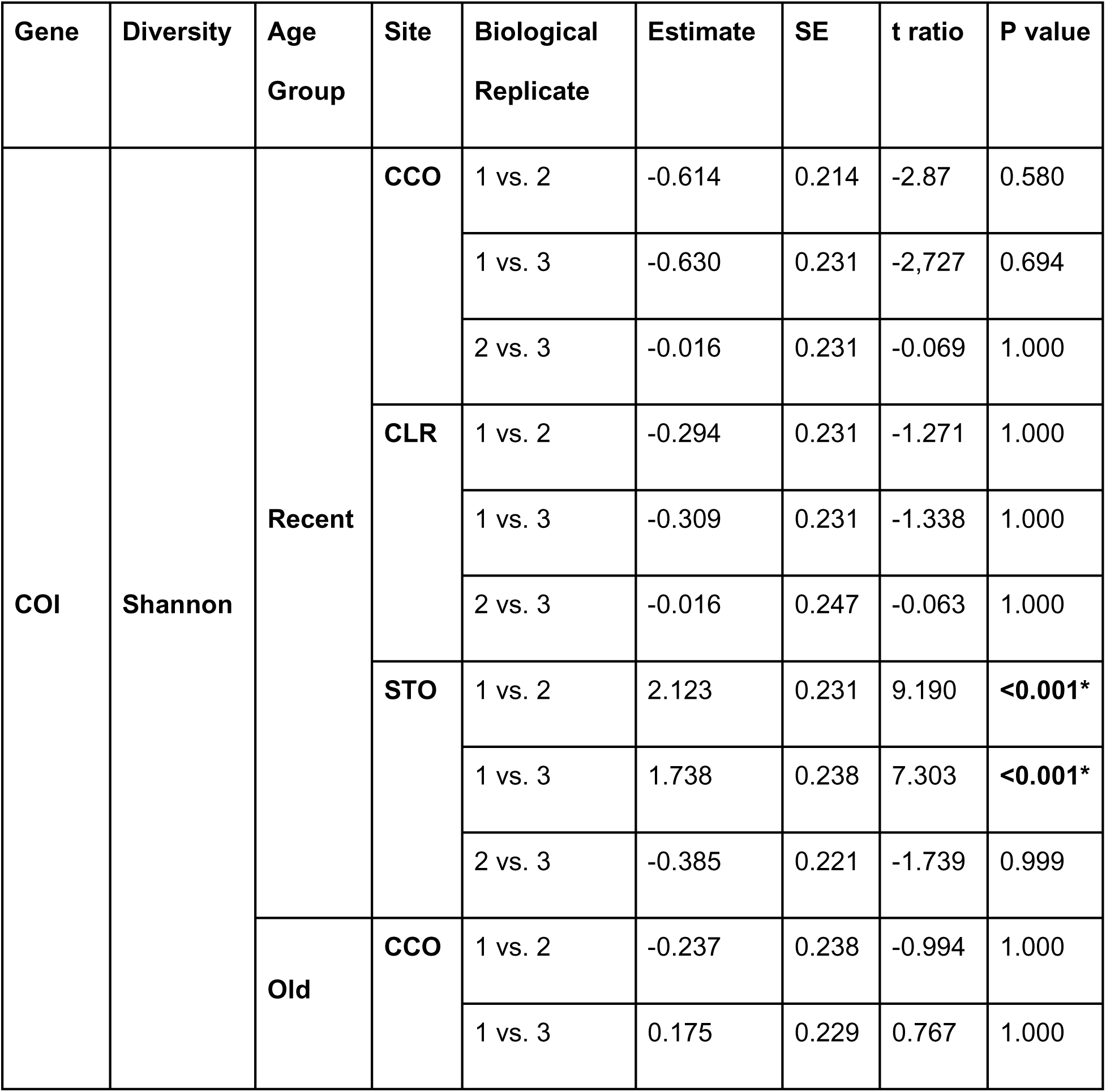
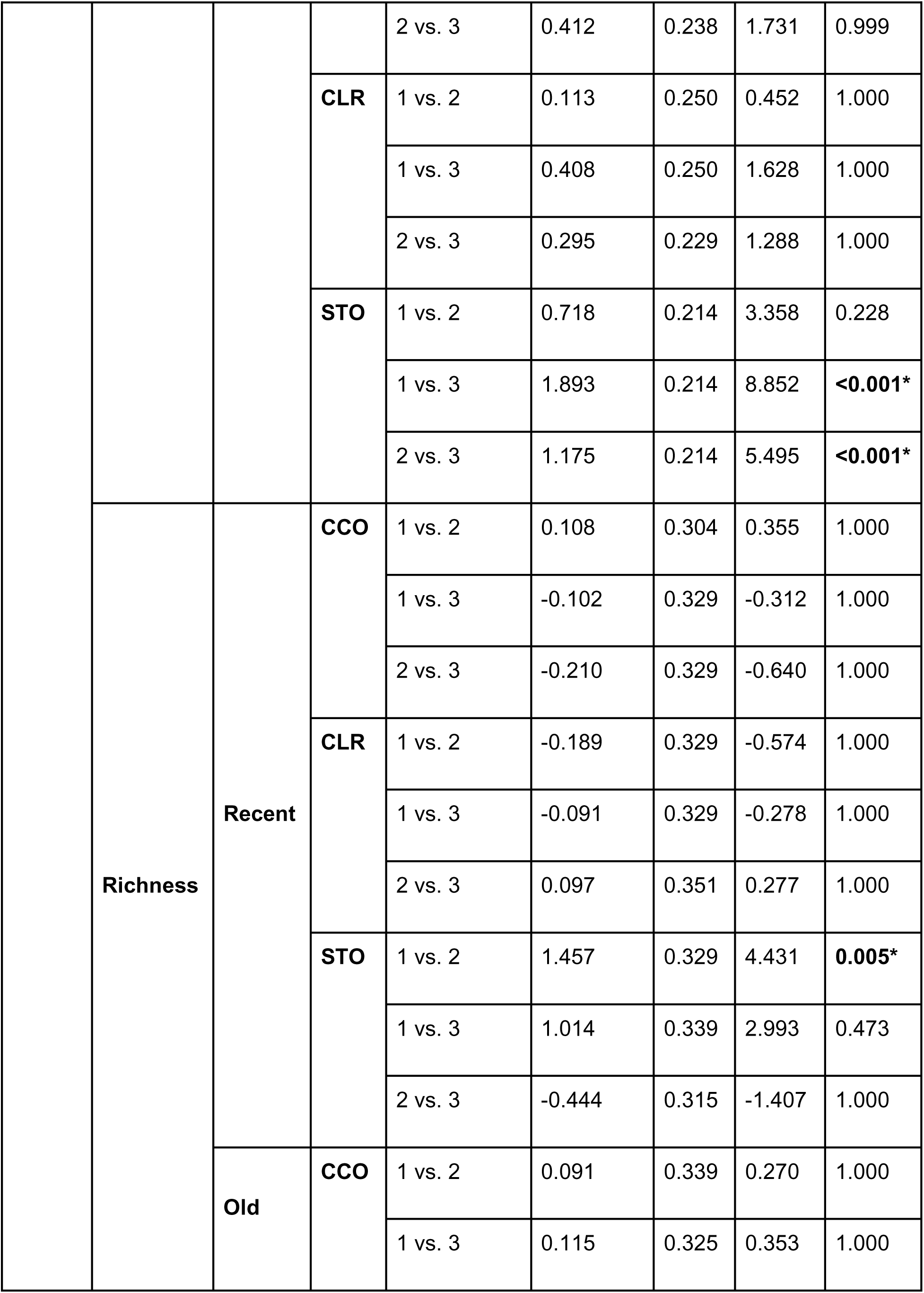
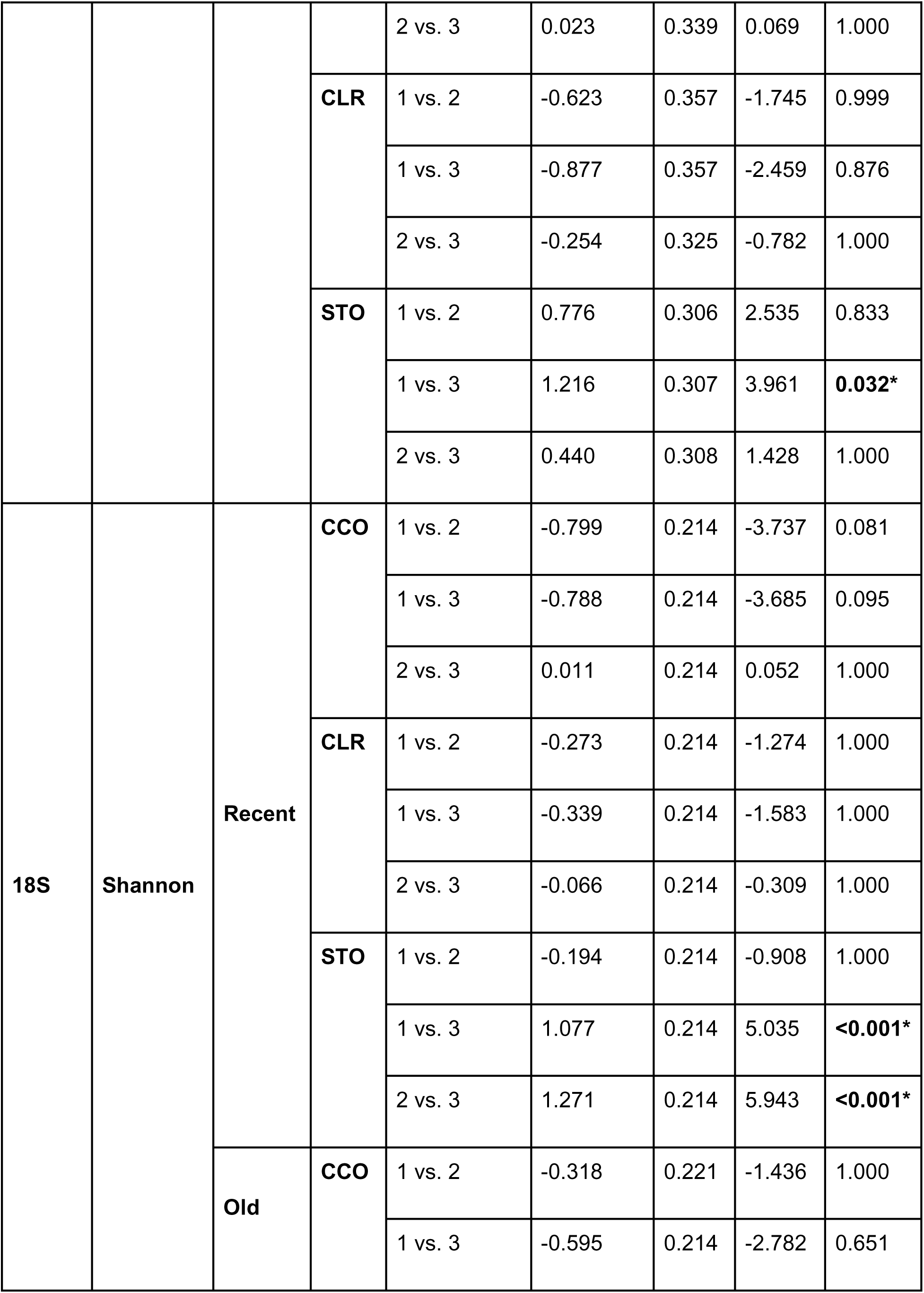
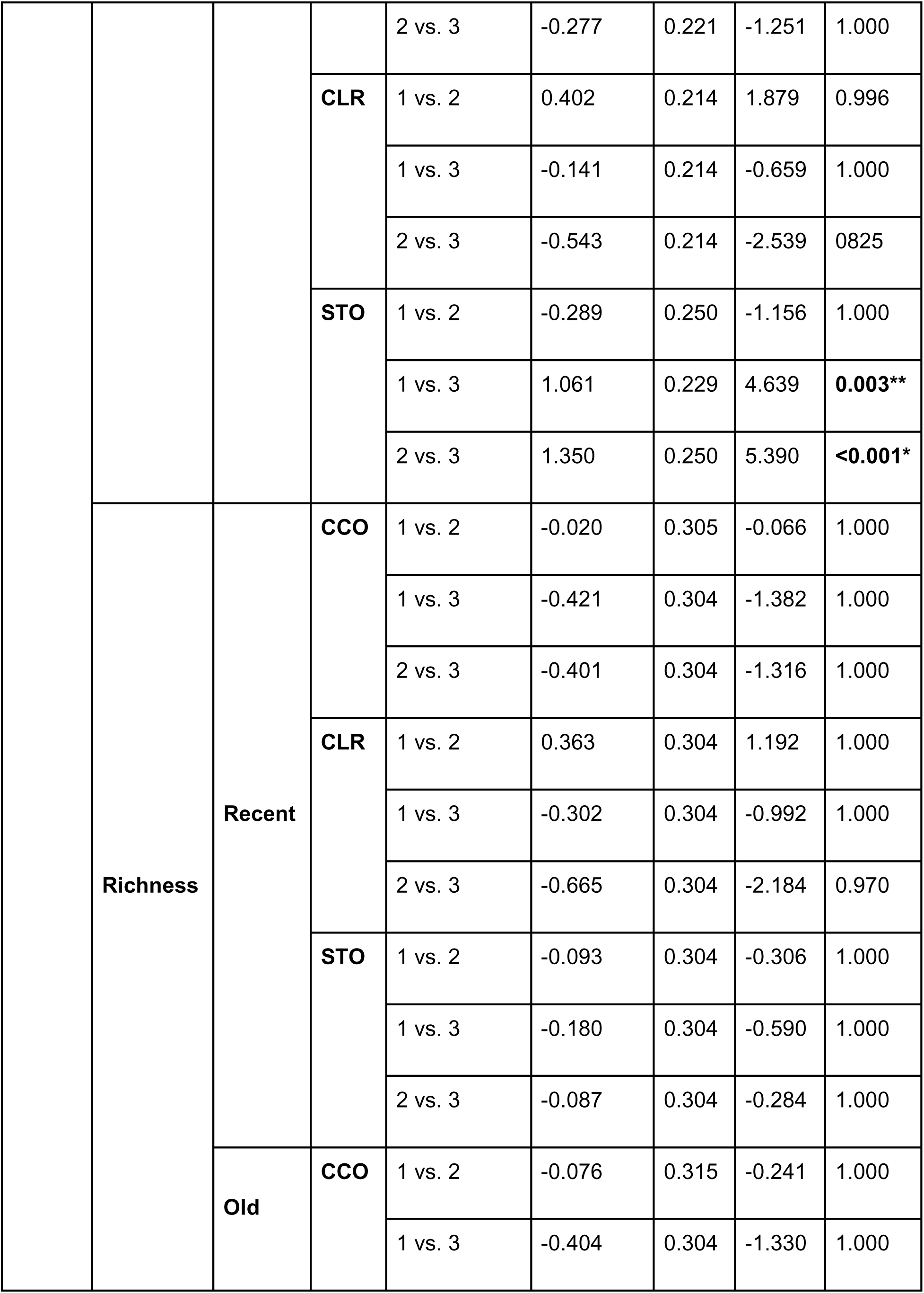
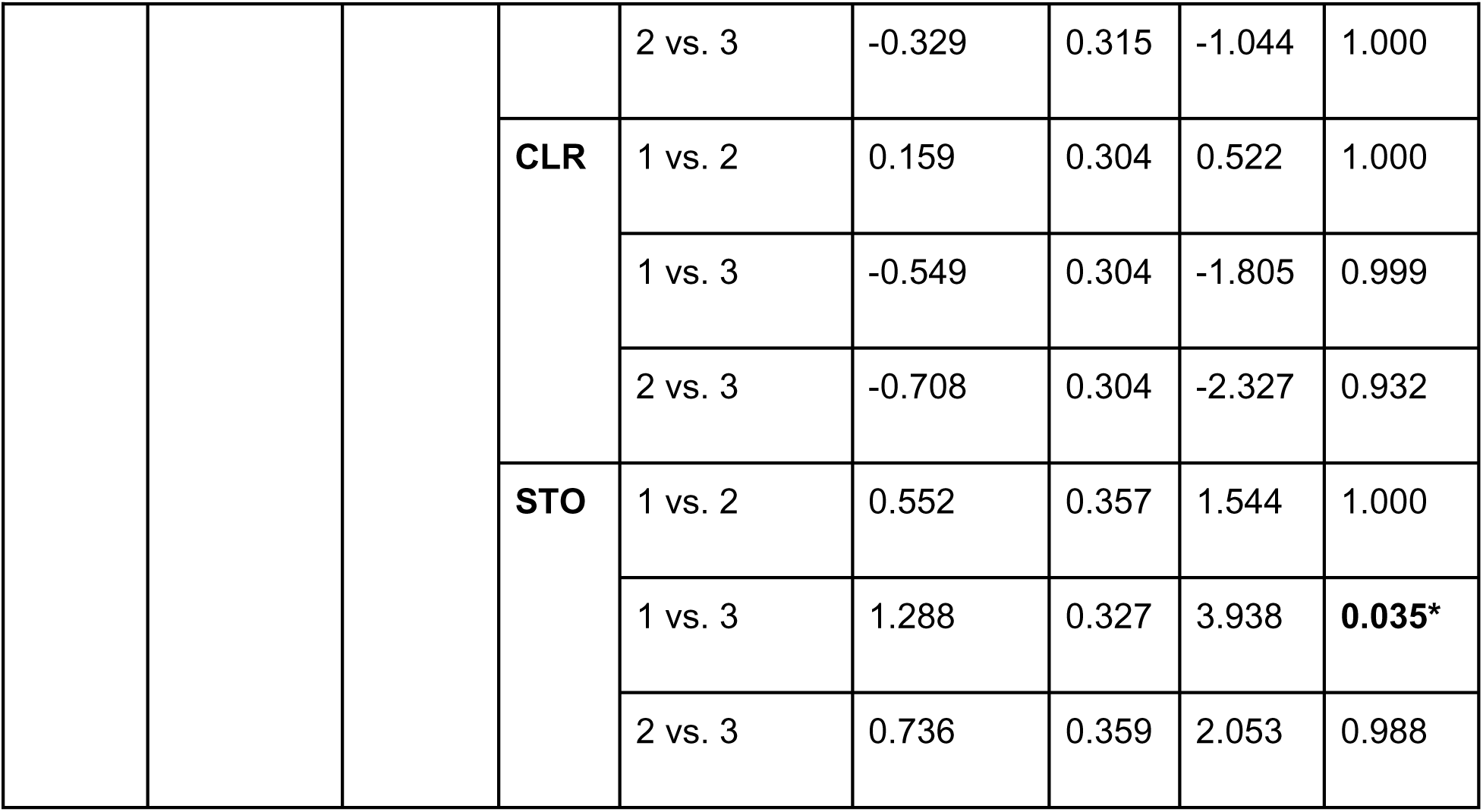
Effects of biological replication (core) on alpha diversity in the complete eukaryotic community. Pairwise comparisons among biological replicates (cores) were performed using linear (Shannon) and negative binomial (richness) models. Core identity (Core_Rep) was included as a fixed effect nested within each Marker × Site × AgeGroup combination. Estimates represent differences between cores (log scale for richness), with standard errors (SE), Wald statistics, and adjusted p-values (Tukey correction). Significant values (p < 0.05) indicate differences among biological replicates within the same marker, site, and sediment age group.

**Table S8:** Effects of biological replication (core) on alpha diversity in the metazoan community. Pairwise comparisons among biological replicates (cores) were performed using linear (Shannon) and negative binomial (richness) models. Core identity (Core_Rep) was included as a fixed effect nested within each Marker × Site × Age Group combination. Estimates represent differences between cores (log scale for richness), with standard errors (SE), Wald statistics, and adjusted p-values (Tukey correction). Significant values (p < 0.05) indicate differences among biological replicates within the same marker, site, and sediment age group.

| Gene | Diversity | Age Group | Site | Biological Replicate | Estimate | SE | t ratio | P value |
| --- | --- | --- | --- | --- | --- | --- | --- | --- |
| COI | Shannon | Recent | CCO | 1 vs. 2 | 0.234 | 0.264 | 0.888 | 1.000 |
|  |  |  |  | 1 vs. 3 | -0.745 | 0.285 | -2.614 | 0.776 |
|  |  |  |  | 2 vs. 3 | -0.979 | 0.285 | -3.436 | 0.188 |
|  |  |  | CLR | 1 vs. 2 | -0.072 | 0.285 | -0.251 | 1.000 |
|  |  |  |  | 1 vs. 3 | -0.146 | 0.285 | -0.513 | 1.000 |
|  |  |  |  | 2 vs. 3 | -0.075 | 0.304 | -0.245 | 1.000 |
|  |  |  | STO | 1 vs. 2 | 0.844 | 0.285 | 2.964 | 0.503 |
|  |  |  |  | 1 vs. 3 | 0.760 | 0.293 | 2.589 | 0.793 |
|  |  |  |  | 2 vs. 3 | -0.085 | 0.273 | -0.310 | 1.000 |
|  |  | Old | CCO | 1 vs. 2 | 0.034 | 0.293 | 0.117 | 1.000 |
|  |  |  |  | 1 vs. 3 | 0.417 | 0.282 | 1.481 | 1.000 |
|  |  |  |  | 2 vs. 3 | 0.383 | 0.293 | 1.306 | 1.000 |
|  |  |  | **CLR** | 1 vs. 2 | 0.026 | 0.309 | 0.085 | 1.000 |
|  |  |  |  | 1 vs. 3 | -0.848 | 0.309 | -2.744 | 0.681 |
|  |  |  |  | 2 vs. 3 | -0.874 | 0.282 | -3.099 | 0.397 |
|  |  |  | **STO** | 1 vs. 2 | 0.075 | 0.264 | 0.286 | 1.000 |
|  |  |  |  | 1 vs. 3 | 0.212 | 0.264 | 0.804 | 1.000 |
|  |  |  |  | 2 vs. 3 | 0.137 | 0.264 | 0.519 | 1.000 |
|  | **Richness** | **Recent** | **CCO** | 1 vs. 2 | 1.226 | 0.353 | 3.473 | 0.157 |
|  |  |  |  | 1 vs. 3 | 0.419 | 0.342 | 1.223 | 1.000 |
|  |  |  |  | 2 vs. 3 | -0.808 | 0.387 | -2.089 | 0.984 |
|  |  |  | **CLR** | 1 vs. 2 | -0.493 | 0.330 | -1.495 | 1.000 |
|  |  |  |  | 1 vs. 3 | 0.018 | 0.344 | 0.051 | 1.000 |
|  |  |  |  | 2 vs. 3 | 0.511 | 0.355 | 1.439 | 1.000 |
|  |  |  | **STO** | 1 vs. 2 | 0.856 | 0.411 | 2.081 | 0.985 |
|  |  |  |  | 1 vs. 3 | 1.157 | 0.454 | 2.551 | 0.824 |
|  |  |  |  | 2 vs. 3 | 0.302 | 0.469 | 0.643 | 1.000 |
|  |  | **Old** | **CCO** | 1 vs. 2 | -0.116 | 0.346 | -0.337 | 1.000 |
|  |  |  |  | 1 vs. 3 | 0.228 | 0.342 | 0.666 | 1.000 |
|  |  |  |  | 2 vs. 3 | 0.344 | 0.352 | 0.977 | 1.000 |
|  |  |  | **CLR** | 1 vs. 2 | -0.159 | 0.362 | -0.412 | 1.000 |
|  |  |  |  | 1 vs. 3 | -0.429 | 0.356 | -1.204 | 1.000 |
|  |  |  |  | 2 vs. 3 | -0.280 | 0.320 | -0.875 | 1.000 |
|  |  |  | **STO** | 1 vs. 2 | 0.118 | 0.550 | 0.214 | 1.000 |
|  |  |  |  | 1 vs. 3 | 0.405 | 0.586 | 0.692 | 1.000 |
|  |  |  |  | 2 vs. 3 | 0.288 | 0.598 | 0.481 | 1.000 |
| **18S** | **Shannon** | **Recent** | **CCO** | 1 vs. 2 | -0.109 | 0.264 | -0.413 | 1.000 |
|  |  |  |  | 1 vs. 3 | 0.020 | 0.264 | 0.075 | 1.000 |
|  |  |  |  | 2 vs. 3 | 0.129 | 0.264 | 0.488 | 1.000 |
|  |  |  | **CLR** | 1 vs. 2 | -0.784 | 0.264 | -2.975 | 0.494 |
|  |  |  |  | 1 vs. 3 | -0.533 | 0.264 | -2.021 | 0.988 |
|  |  |  |  | 2 vs. 3 | 0.252 | 0.264 | 0.954 | 1.000 |
|  |  |  | **STO** | 1 vs. 2 | 1.195 | 0.264 | 4.532 | **0.005\*** |
|  |  |  |  | 1 vs. 3 | 2.035 | 0.264 | 7.715 | **<0.001\*** |
|  |  |  |  | 2 vs. 3 | 0.840 | 0.264 | 3.184 | 0.336 |
|  |  | **Old** | **CCO** | 1 vs. 2 | 0.329 | 0.273 | 1.204 | 1.000 |
|  |  |  |  | 1 vs. 3 | 0.074 | 0.264 | 0.280 | 1.000 |
|  |  |  |  | 2 vs. 3 | -0.255 | 0.273 | -0.934 | 1.000 |
|  |  |  | **CLR** | 1 vs. 2 | 0.541 | 0.264 | 2.050 | 0.986 |
|  |  |  |  | 1 vs. 3 | 0.258 | 0.264 | 0.978 | 1.000 |
|  |  |  |  | 2 vs. 3 | -0.283 | 0.264 | -1.072 | 1.000 |
|  |  |  | **STO** | 1 vs. 2 | 0.763 | 0.309 | 2.470 | 0.863 |
|  |  |  |  | 1 vs. 3 | 1.141 | 0.282 | 4.048 | **0.029\*** |
|  |  |  |  | 2 vs. 3 | 0.378 | 0.309 | 1.225 | 1.000 |
|  | **Richness** | **Recent** | **CCO** | 1 vs. 2 | 0.098 | 0.263 | 0.371 | 1.000 |
|  |  |  |  | 1 vs. 3 | -0.007 | 0.263 | -0.025 | 1.000 |
|  |  |  |  | 2 vs. 3 | -0.104 | 0.263 | -0.396 | 1.000 |
|  |  |  | **CLR** | 1 vs. 2 | 0.491 | 0.263 | 1.867 | 0.998 |
|  |  |  |  | 1 vs. 3 | -0.136 | 0.261 | -0.520 | 1.000 |
|  |  |  |  | 2 vs. 3 | -0.627 | 0.263 | -2.386 | 0.910 |
|  |  |  | **STO** | 1 vs. 2 | 1.191 | 0.275 | 4.338 | **0.007\*** |
|  |  |  |  | 1 vs. 3 | 0.271 | 0.267 | 1.016 | 1.000 |
|  |  |  |  | 2 vs. 3 | -0.921 | 0.276 | -3.336 | 0.226 |
|  |  | **Old** | **CCO** | 1 vs. 2 | 0.162 | 0.280 | 0.578 | 1.000 |
|  |  |  |  | 1 vs. 3 | -0.129 | 0.269 | -0.480 | 1.000 |
|  |  |  |  | 2 vs. 3 | -0.291 | 0.279 | -1.042 | 1.000 |
|  |  |  | **CLR** | 1 vs. 2 | 0.274 | 0.264 | 1.039 | 1.000 |
|  |  |  |  | 1 vs. 3 | -0.345 | 0.262 | -1.319 | 1.000 |
|  |  |  |  | 2 vs. 3 | -0.619 | 0.263 | -2.357 | 0.922 |
|  |  |  | **STO** | 1 vs. 2 | 1.200 | 0.336 | 3.571 | 0.118 |
|  |  |  |  | 1 vs. 3 | 1.169 | 0.304 | 3.850 | **0.048\*** |
|  |  |  |  | 2 vs. 3 | -0.031 | 0.349 | -0.090 | 1.000 |

**Table S9:** Effects of site on alpha diversity in the complete eukaryotic community. Pairwise comparisons among sites were performed using mixed-effects models including Site, Marker, and Age Group as fixed effects, with biological replicates (cores) included as a random effect. Estimates represent differences between sites within each Marker × Age Group combination, with standard errors (SE), z-values, and adjusted p-values (Tukey correction).

| Gene | Diversity | Age Group | Site Intercept | Estimate | SE | t ratio | P value |
| --- | --- | --- | --- | --- | --- | --- | --- |
| COI | Shannon | Recent | CCO-CLR | -0.595 | 0.388 | -1.533 | 0.328 |
|  |  |  | CCO-STO | 0.513 | 0.387 | 1.324 | 0.423 |
|  |  |  | CLR-STO | 1.107 | 0.389 | 2.849 | 0.052 |
|  |  | Old | CCO-CLR | 0.753 | 0.390 | 1.930 | 0.193 |
|  |  |  | CCO-STO | 2.059 | 0.387 | 5.326 | **0.002\*** |
|  |  |  | CLR-STO | 1.306 | 0.387 | 3.370 | **0.025\*** |
|  | Richness | Recent | CCO-CLR | -0.203 | 0.267 | -0.760 | 0.727 |
|  |  |  | CCO-STO | 0.682 | 0.266 | 2.564 | **0.028\*** |
|  |  |  | CLR-STO | 0.885 | 0.270 | 3.281 | **0.003\*** |
|  |  | Old | CCO-CLR | 0.576 | 0.274 | 2.104 | 0.089 |
|  |  |  | CCO-STO | 2.256 | 0.265 | 8.509 | **< 0.001\*** |
|  |  |  | CLR-STO | 1.680 | 0.268 | 6.270 | **< 0.001\*** |
| 18S | Shannon | Recent | CCO-CLR | -0.772 | 0.384 | -2.011 | 0.176 |
|  |  |  | CCO-STO | -0.089 | 0.384 | -0.231 | 0.971 |
|  |  |  | CLR-STO | 0.683 | 0.384 | 1.779 | 0.240 |
|  |  | **Old** | CCO-CLR | -1.006 | 0.384 | -2.616 | 0.075 |
|  |  |  | CCO-STO | -0.708 | 0.388 | -1.825 | 0.224 |
|  |  |  | CLR-STO | 0.298 | 0.387 | 0.770 | 0.732 |
|  | **Richness** | **Recent** | CCO-CLR | -0.502 | 0.258 | -1.950 | 0.125 |
|  |  |  | CCO-STO | -0.121 | 0.258 | -0.470 | 0.886 |
|  |  |  | CLR-STO | 0.381 | 0.259 | 1.473 | 0.304 |
|  |  | **Old** | CCO-CLR | -0.437 | 0.274 | -1.590 | 0.210 |
|  |  |  | CCO-STO | 1.330 | 0.269 | 4.953 | **< 0.001\*** |
|  |  |  | CLR-STO | 1.768 | 0.267 | 6.616 | **< 0.001\*** |

**Table S10:** Effects of site on alpha diversity in the metazoan community. Pairwise comparisons among sites were performed using mixed-effects models including Site, Marker, and Age Group as fixed effects, with biological replicates (cores) included as a random effect. Estimates represent differences between sites within each Marker × Age Group combination, with standard errors (SE), z-values, and adjusted p-values (Tukey correction).

| Gene | Diversity | Age Group | Site Intercept | Estimate | SE | t ratio | P value |
| --- | --- | --- | --- | --- | --- | --- | --- |
| COI | Shannon | Recent | CCO-CLR | 0.591 | 0.308 | 1.921 | 0.180 |
|  |  |  | CCO-STO | 0.019 | 0.306 | 0.061 | 0.998 |
|  |  |  | CLR-STO | -0.573 | 0.309 | -1.854 | 0.198 |
|  |  | Old | CCO-CLR | -0.012 | 0.312 | -0.040 | 0.999 |
|  |  |  | CCO-STO | 1.156 | 0.306 | 3.783 | **0.009\*** |
|  |  |  | CLR-STO | 1.169 | 0.307 | 3.805 | **0.008\*** |
|  | Richness | Recent | CCO-CLR | -0.375 | 0.281 | -1.335 | 0.376 |
|  |  |  | CCO-STO | 0.799 | 0.301 | 2.657 | **0.022\*** |
|  |  |  | CLR-STO | 1.174 | 0.300 | 3.910 | **<0.001\*** |
|  |  | Old | CCO-CLR | -0.286 | 0.282 | -1.015 | 0.568 |
|  |  |  | CCO-STO | 2.087 | 0.340 | 6.143 | **<0.001\*** |
|  |  |  | CLR-STO | 2.373 | 0.339 | 7.000 | **<0.001\*** |
| 18S | Shannon | Recent | CCO-CLR | -0.390 | 0.301 | -1.295 | 0.430 |
|  |  |  | CCO-STO | 0.032 | 0.301 | 0.107 | 0.994 |
|  |  |  | CLR-STO | 0.422 | 0.301 | 1.403 | 0.377 |
|  |  | Old | CCO-CLR | -0.059 | 0.302 | -0.196 | 0.979 |
|  |  |  | CCO-STO | 1.039 | 0.308 | 3.374 | **0.016\*** |
|  |  |  | CLR-STO | 1.099 | 0.307 | 3.578 | **0.012\*** |
|  | Richness | Recent | CCO-CLR | -0.242 | 0.250 | -0.971 | 0.595 |
|  |  |  | CCO-STO | 0.705 | 0.252 | 2.803 | **0.014\*** |
|  |  |  | CLR-STO | 0.947 | 0.251 | 3.769 | **<0.001\*** |
|  |  | Old | CCO-CLR | -0.793 | 0.252 | -3.146 | **0.005\*** |
|  |  |  | CCO-STO | 0.785 | 0.265 | 2.962 | **0.009\*** |
|  |  |  | CLR-STO | 1.578 | 0.263 | 6.007 | **< 0.001\*** |

**Table S11:** Mean variance explained by fixed (site, sediment age group) and random effects, alongside model performance metrics (AUC and Tjur’s R²), across all jSDM models. Mean variance partitioning and model performance metrics for hierarchical models of species communities (jSDM), including site, sediment age group, and random effects.

| Community | Marker | Site | Age Group | Random:<br>Site | Random:<br>Core | Mean<br>AUC | Mean<br>Tjur's $R^2$ |
| --- | --- | --- | --- | --- | --- | --- | --- |
| Eukariotic | COI | 0.669 | 0.213 | 0.060 | 0.059 | 0.941 | 0.234 |
|  | 18S | 0.766 | 0.140 | 0.049 | 0.045 | 0.929 | 0.277 |
| Metazoa | COI | 0.630 | 0.283 | 0.046 | 0.040 | 0.959 | 0.263 |
|  | 18S | 0.740 | 0.164 | 0.047 | 0.049 | 0.935 | 0.220 |

**Table S12:** Results of PERMANOVA (*adonis2*) for all Eukaryotic community and Metazoan community separated between COI and 18S community composition based on Bray–Curtis dissimilarities. The effects of site, sediment age group and biological replicates were tested using marginal permutation tests (999 permutations). R² values represent the proportion of variance explained by each factor, while the residual indicates unexplained variation in community composition not accounted for by the model. Significance codes: *** p < 0.001; ** p < 0.01; * p < 0.05; ns, not significant.

| Community | Marker | Factor | df | R<sup>2</sup> | F | p-value |
| --- | --- | --- | --- | --- | --- | --- |
| Eukaryotic | COI | Biological Replicates | 2 | 0.032 | 0.764 | 0.712 |
|  |  | Site | 2 | 0.272 | 3.285 | 0.001 |
|  |  | Age group | 1 | 0.157 | 3.797 | 0.001 |
|  |  | Residual /Unexplained variation | 11 | 0.539 |  |  |
|  | 18S | Biological Replicates | 2 | 0.029 | 0.825 | 0.516 |
|  |  | Site | 2 | 0.345 | 4.977 | 0.001 |
|  |  | Age group | 1 | 0.177 | 5.114 | 0.002 |
|  |  | Residual /Unexplained variation | 11 | 0.451 |  |  |
| Metazoan | COI | Biological Replicates | 2 | 0.043 | 0.808 | 0.650 |
|  |  | Site | 2 | 0.147 | 1.367 | 0.120 |
|  |  | Age group | 1 | 0.116 | 2.170 | 0.022 |
|  |  | Residual /Unexplained variation | 11 | 0.697 |  |  |
|  | 18S | Biological Replicates | 2 | 0.046 | 0.976 | 0.442 |
|  |  | Site | 2 | 0.203 | 3.127 | 0.001 |
|  |  | Age group | 1 | 0.133 | 2.785 | 0.001 |
|  |  | Residual /Unexplained variation | 11 | 0.619 |  |  |

**Table S13:**
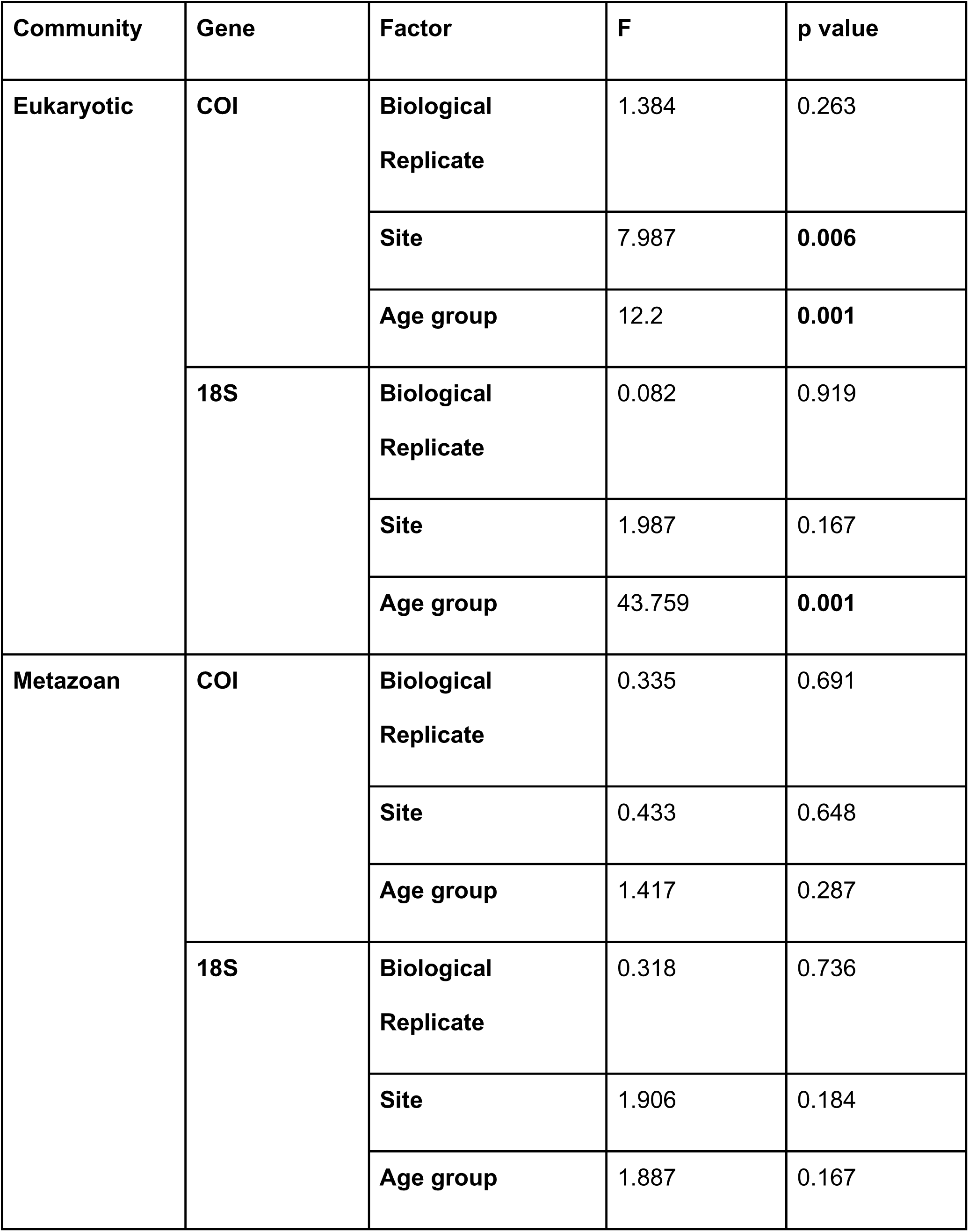
Permutation tests for homogeneity of multivariate dispersion (*betadisper*) for eukaryotic and metazoan datasets. The effects of site, sediment age group and biological replicates were tested using marginal permutation tests (999 permutations).

## Supplementary figures

**Fig. S1:**
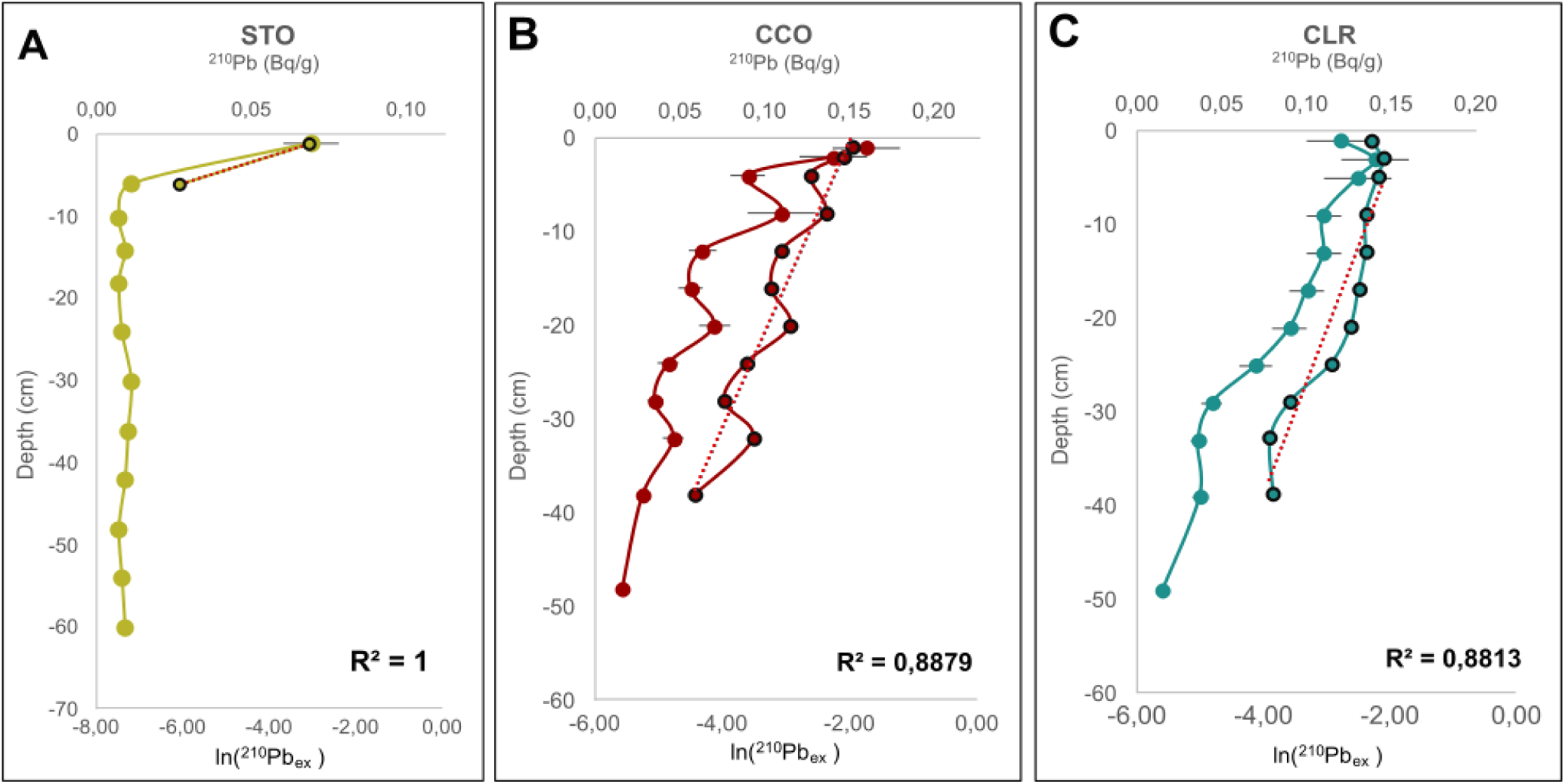
^210^Pb activity-depth profiles for sediment cores from (A) Santoña (STO), (B) Cicero (CCO), and (C) Colindres (CLR). ^210^Pb activities are shown against the upper x-axis, while the natural logarithm of excess or unsupported ^210^Pb is shown against the lower x-axis. Points with black outlines represent ln-transformed excess ^210^Pb values used to estimate sediment accumulation rates using the “Simple” Model (Robbins, 1978, dotted regression lines). Error bars represent analytical uncertainty. In the STO core, excess ^210^Pb activity is confined to the upper ∼10 cm, indicating a highly condensed sediment sequence.

**Fig. S2:**
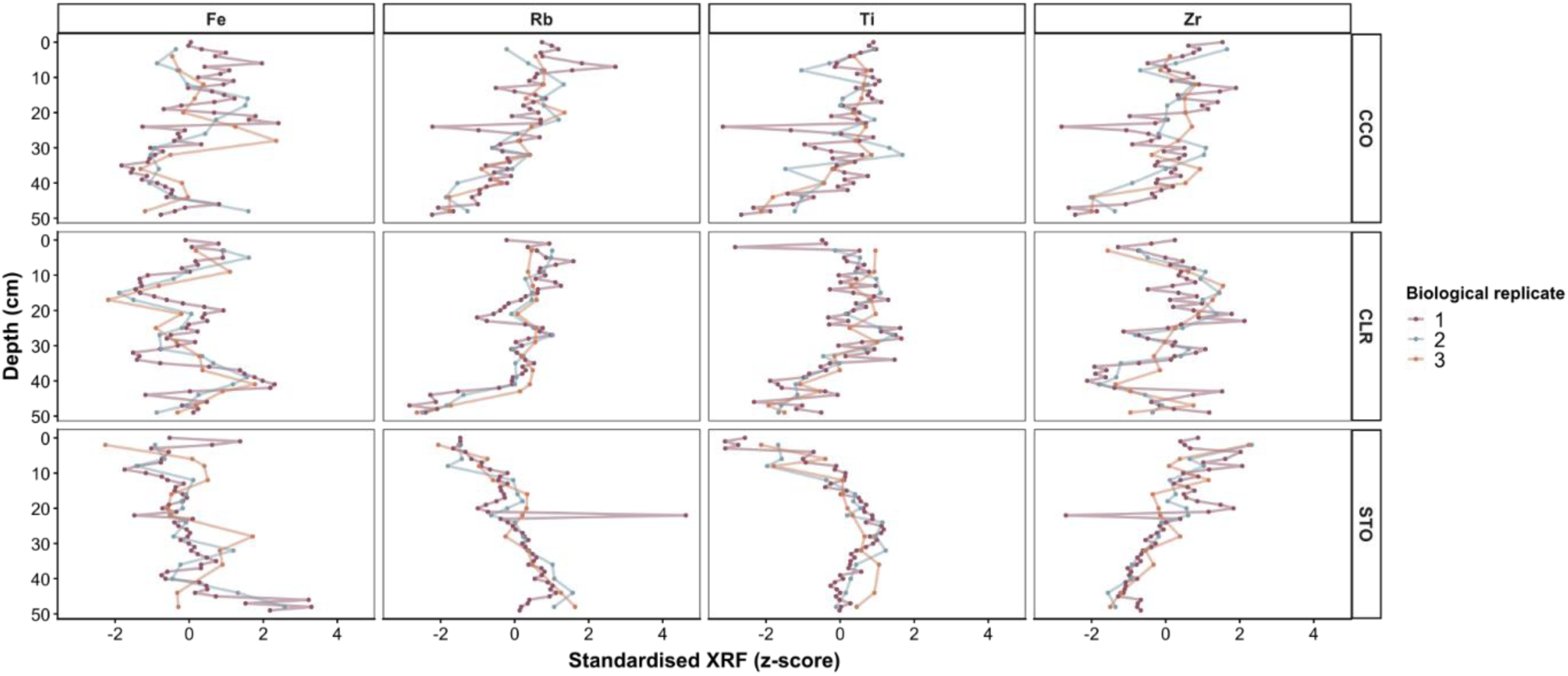
Standardised geochemical XRF profiles across biological replicates (cores) at sites CCO and CLR. Elemental concentrations (Fe, Rb, Ti and Zr) are shown as depth-standardised z-scores to facilitate comparison among cores. Lines represent individual biological replicates and illustrate closely matching stratigraphic trends across spatially separated cores within each site.

**Fig. S3:**
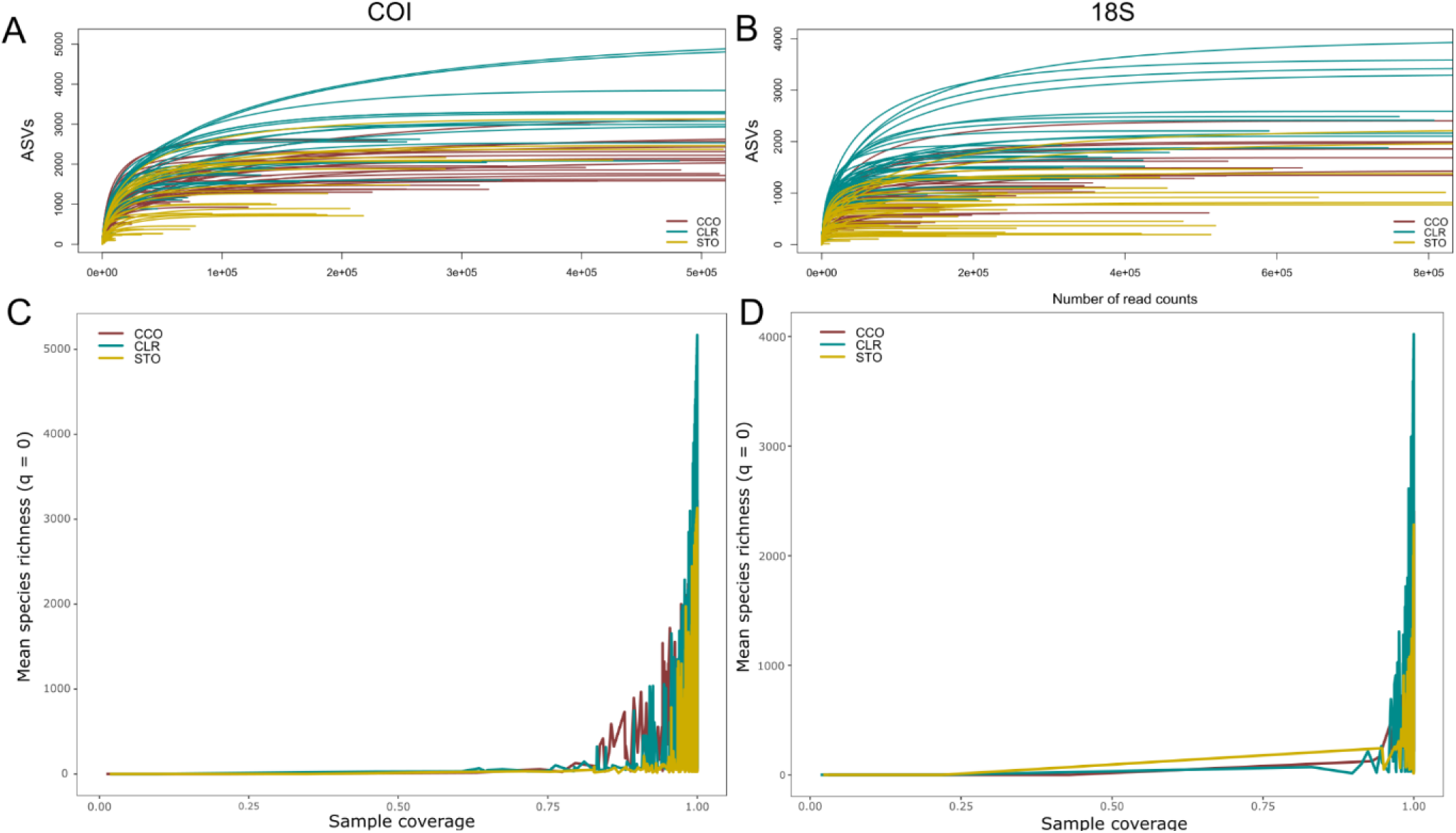
Rarefaction and coverage-based diversity estimates for metabarcoding-based *seda*DNA datasets. Panels (A) and (B) show sample-size-based rarefaction and extrapolation curves for the COI and 18S markers, respectively. Solid lines represent interpolation and dashed lines extrapolation, with shaded areas indicating 95% confidence intervals. Panels (C) and (D) show coverage-based rarefaction and extrapolation curves derived from iNEXT for COI and 18S, respectively, illustrating sample completeness across sites.

**Fig. S4:**
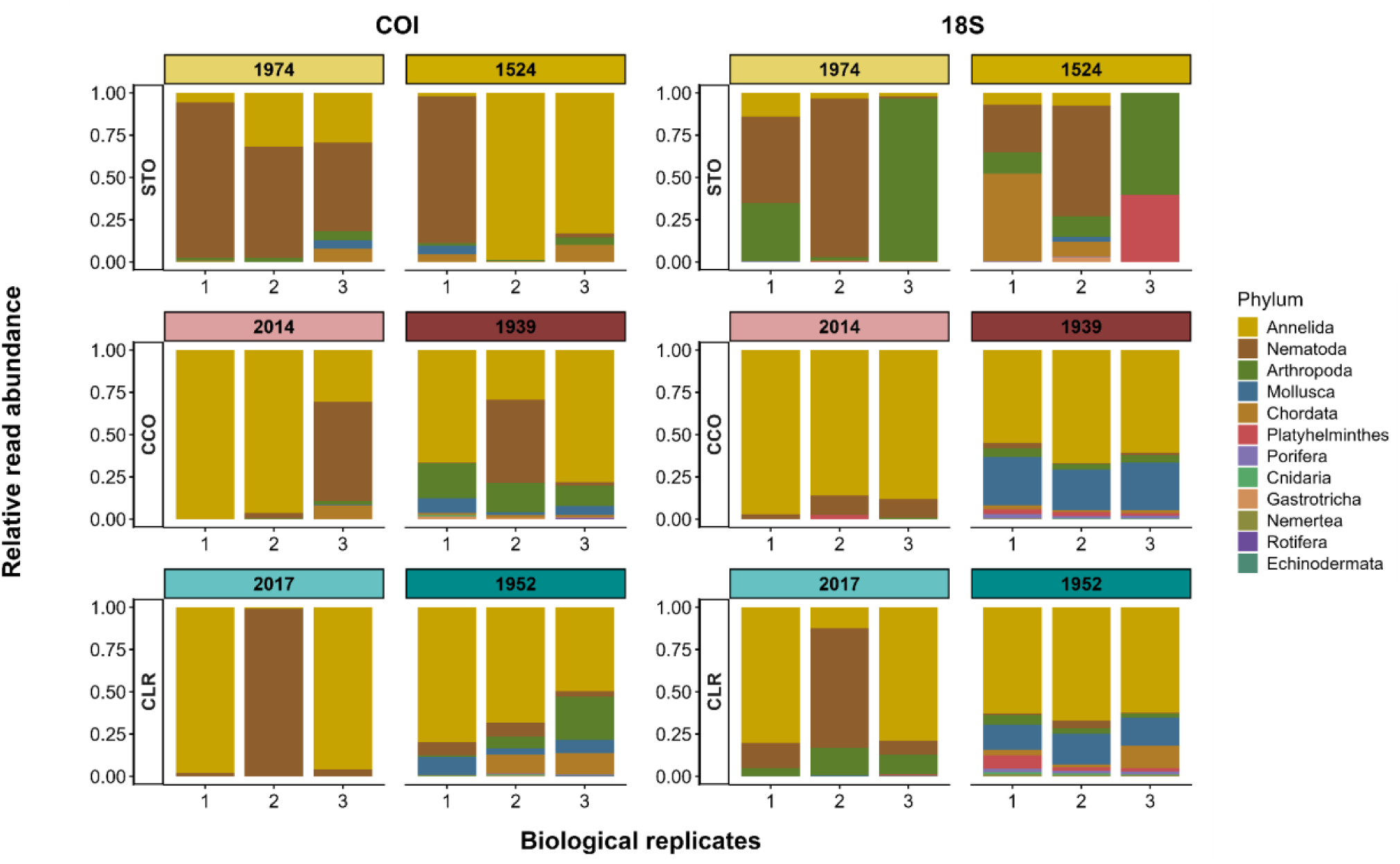
Relative abundance of metazoan phyla detected across the three intertidal sites (STO, CCO and CLR) based on metabarcoding-based *seda*DNA. Bars represent the mean relative abundance of amplicon sequence variants (ASVs) per core after collapsing eight technical PCR replicates. Facets indicate genetic markers (COI or 18S) and sediment age group.

**Fig. S5:**
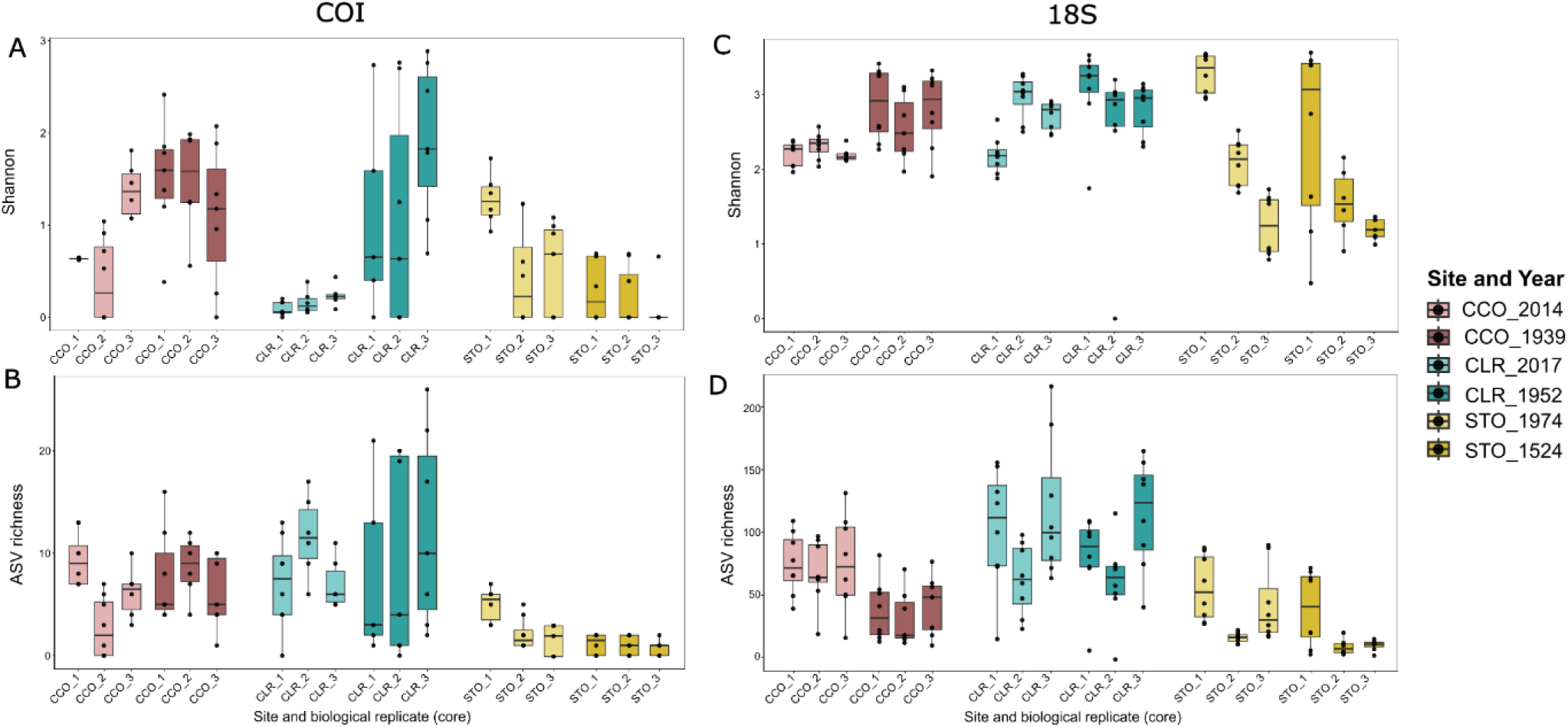
Boxplots show variation in (A) Shannon diversity index for COI and (C) for 18S, and (B) ASV richness for COI and (D) for 18S, among biological replicates (cores) across sites and sediment age group for all the metazoan community. Each box represents the distribution of technical replicates for a given biological replicate and sediment age group, overlaid as individual points. Colours indicate site and sediment age group, with lighter shades representing recent sediments and darker shades representing deeper (“old”) horizons.

**Fig. S6:**
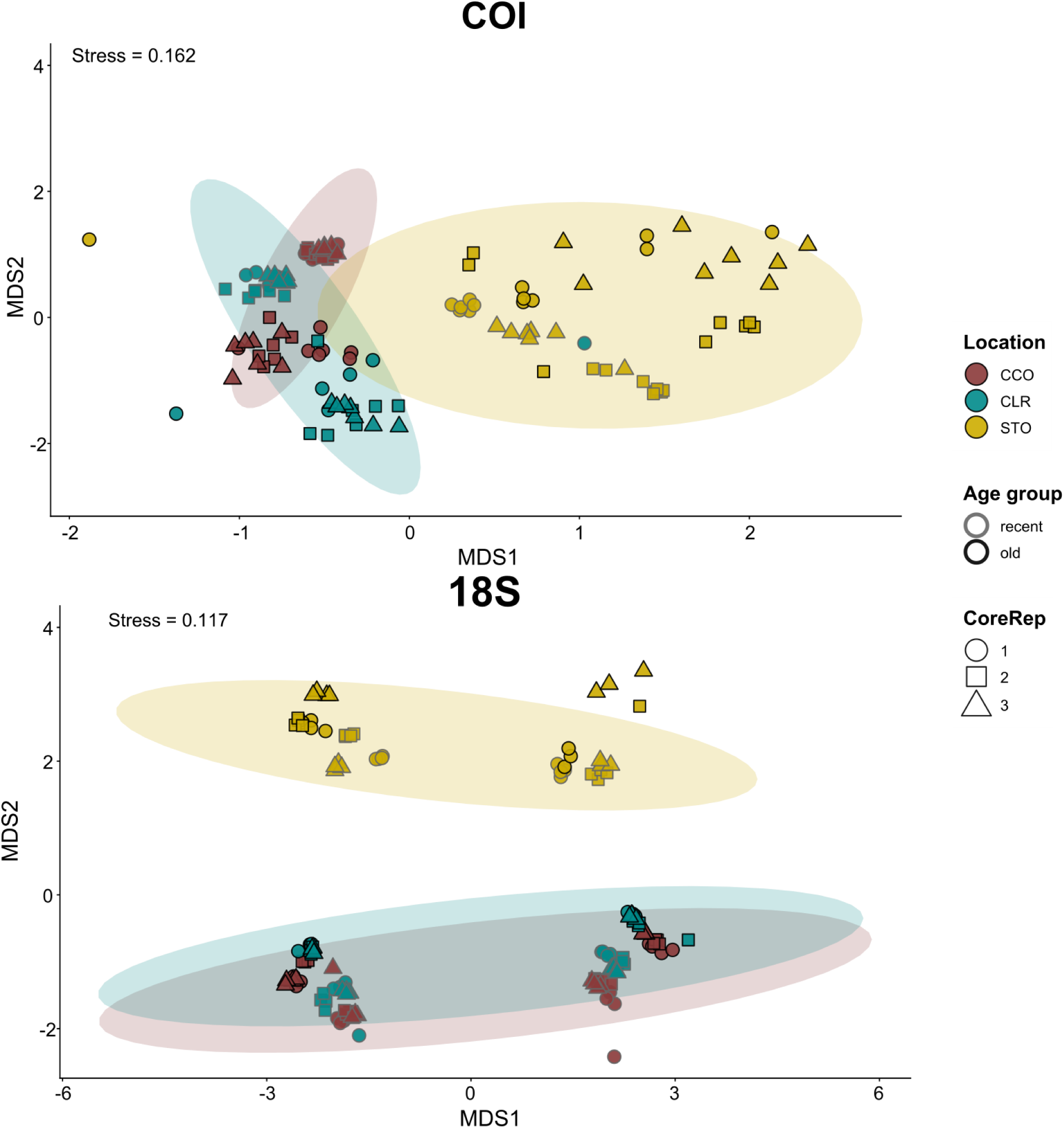
NMDS ordination including individual PCR replicates (technical replicates) showing increased within-sample dispersion compared to analyses based on averaged community profiles. The plot shows axes 1 and 2 of the two-dimensional NMDS solution based on Bray– Curtis dissimilarities of ASV relative abundances (stress value indicated in the panel). Points are coloured by site (CCO, CLR, STO) and sediment age group (recent vs. old), and shaped according to biological replicate (CoreRep) while shaded ellipses delimit the dispersion of biological replicates (cores) within each site.

**Fig. S7:**
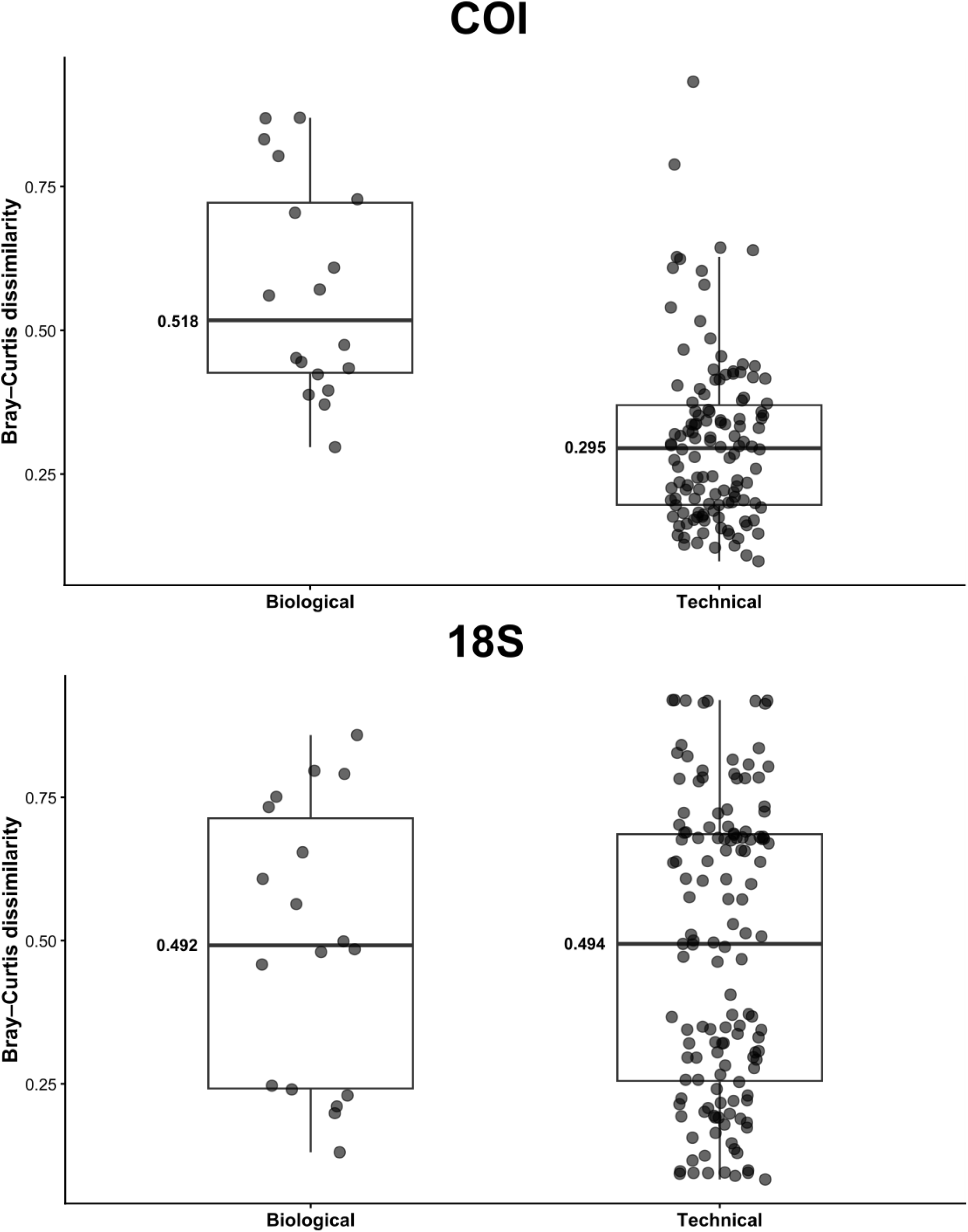
Comparison of Bray–Curtis dissimilarities between technical and biological replicates for each marker (COI and 18S). Distributions are shown as boxplots, with the median indicated by a bold horizontal line within each box.

**Fig. S8:**
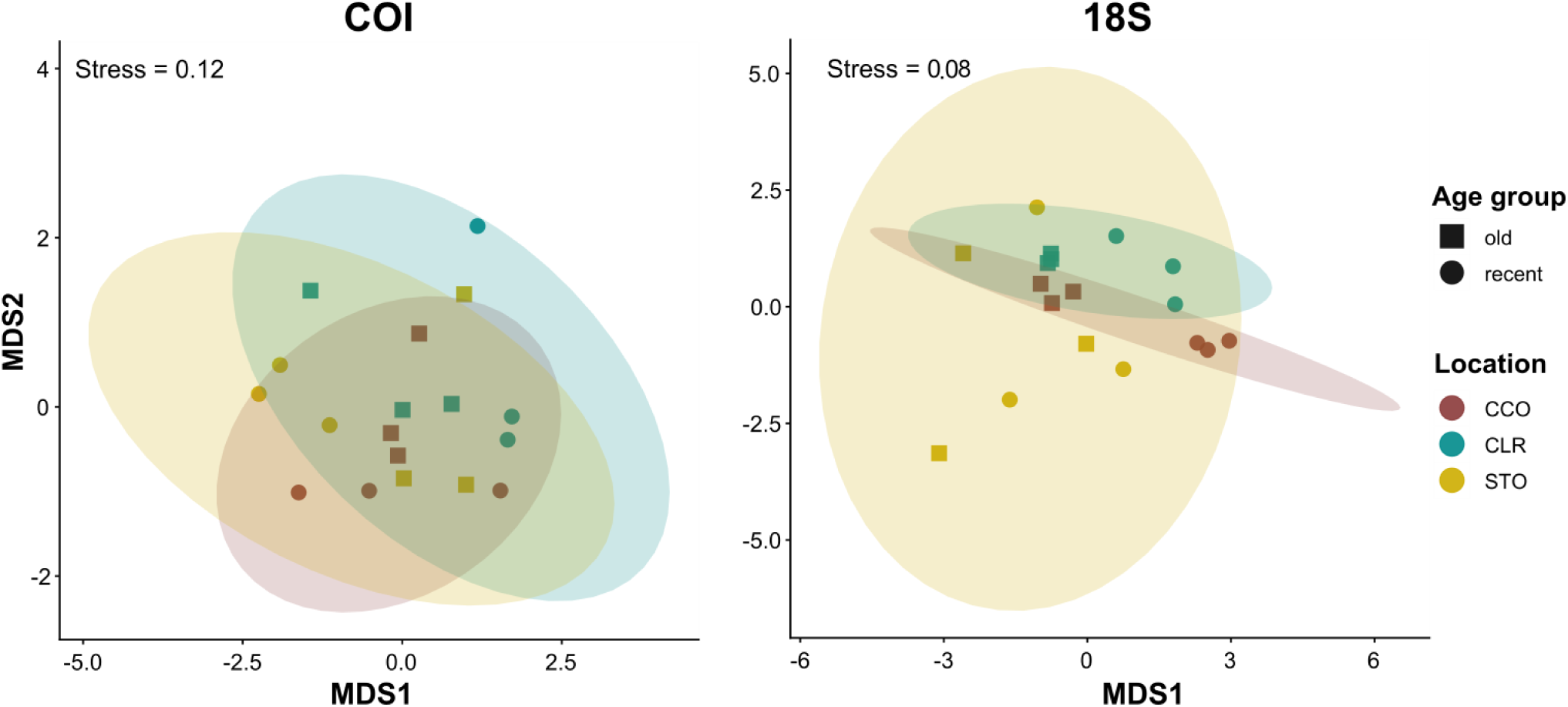
Non-metric multidimensional scaling (NMDS) ordination of COI- and 18S-based metabarcoding-based *seda*DNA Metazoan community composition across the three intertidal sites. The plot shows axes 1 and 2 of the two-dimensional NMDS solution based on Bray– Curtis dissimilarities of ASV relative abundances (stress value indicated in the panel). Points are coloured by site (CCO, CLR, STO) and shaped according to sediment age group (recent vs. old), while shaded ellipses delimit the dispersion of biological replicates (cores) within each site.

**Fig. S9:**
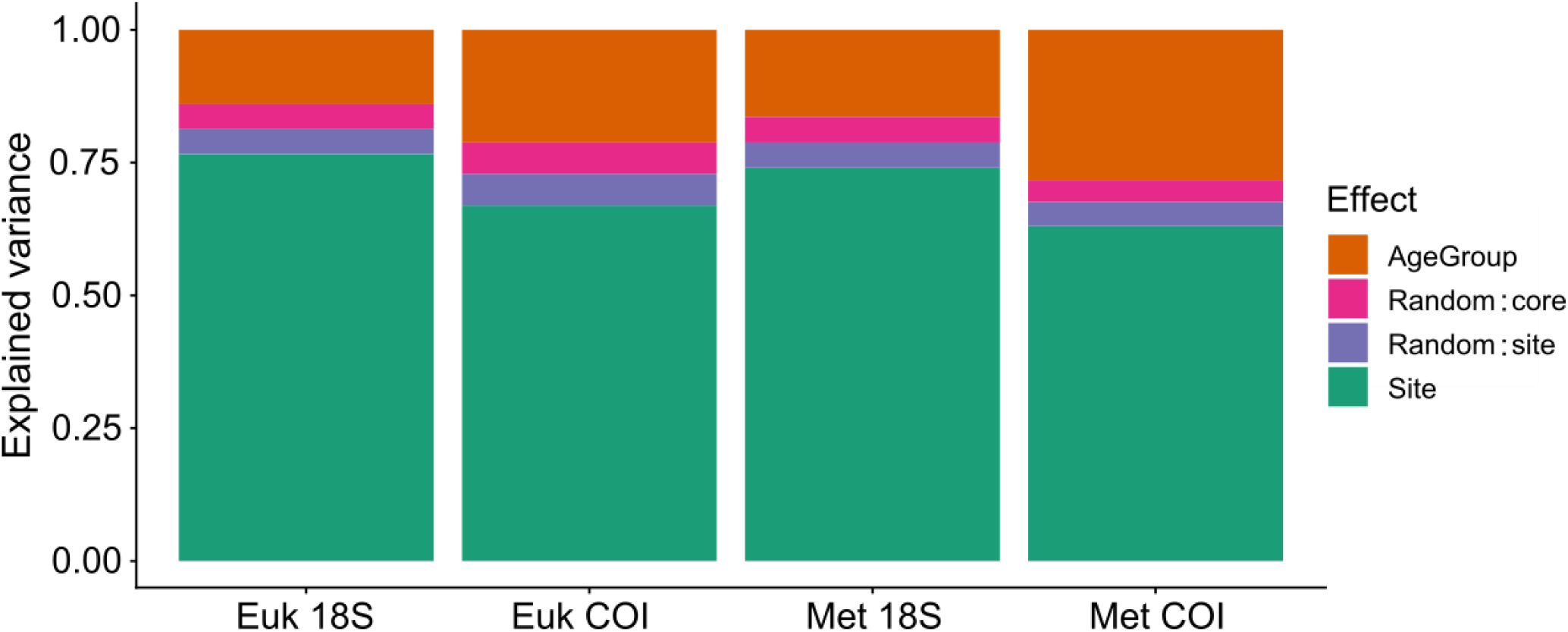
Relative contribution of environmental and spatial factors to community structure across datasets. Stacked barplots showing the mean proportion of variance explained by site, sediment age group, and random effects (site and core) across all taxa for each dataset (Eukaryota and Metazoa; COI and 18S markers), based on hierarchical modelling of species communities (jSDM). Values represent posterior means of variance partitioning.

**Fig. S10:**
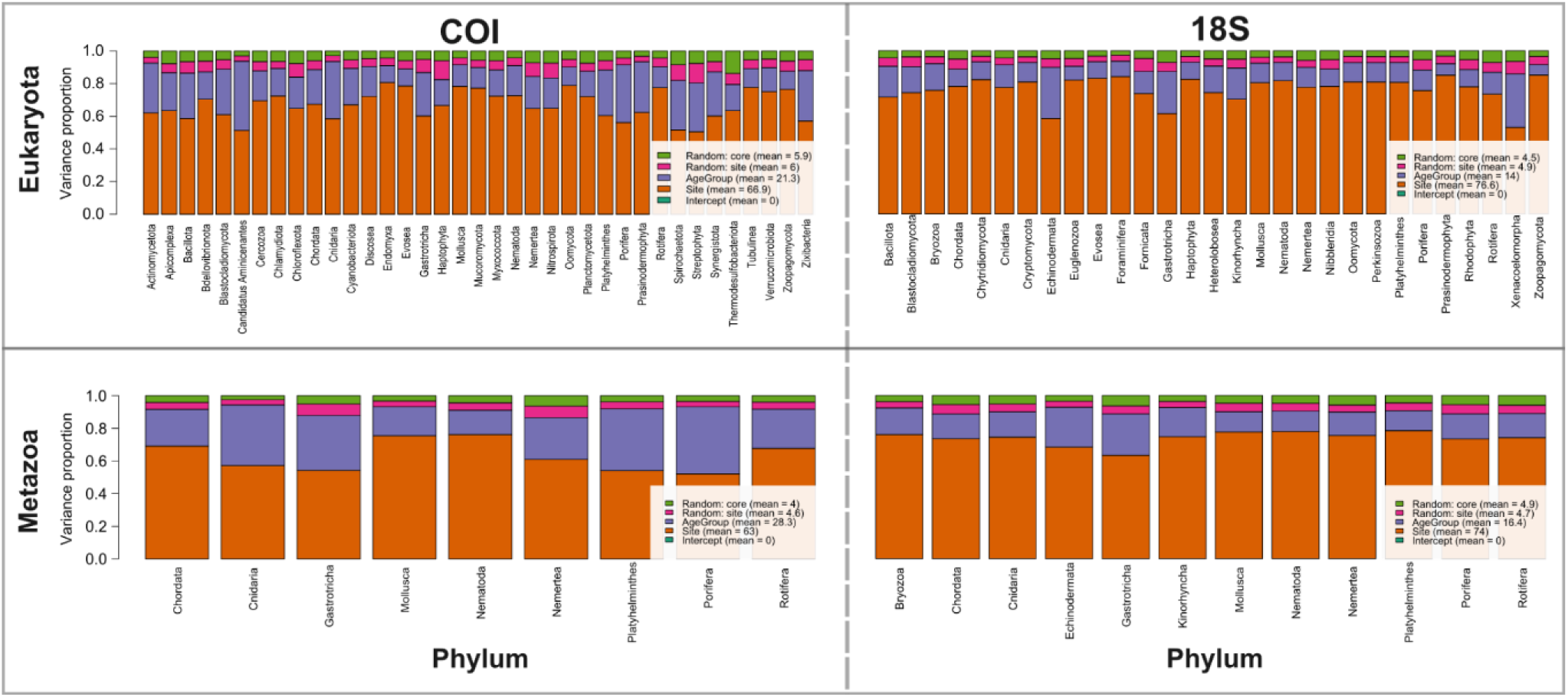
Variance partitioning at the phylum level across datasets. Stacked barplots showing the proportion of variance explained by site, sediment age group, and random effects (site and core) for each phylum across all datasets (Eukaryota and Metazoa; COI and 18S markers), based on hierarchical modelling of species communities (jSDM). Each bar represents an individual phylum. Values represent posterior means of variance partitioning.

**Fig. S11:**
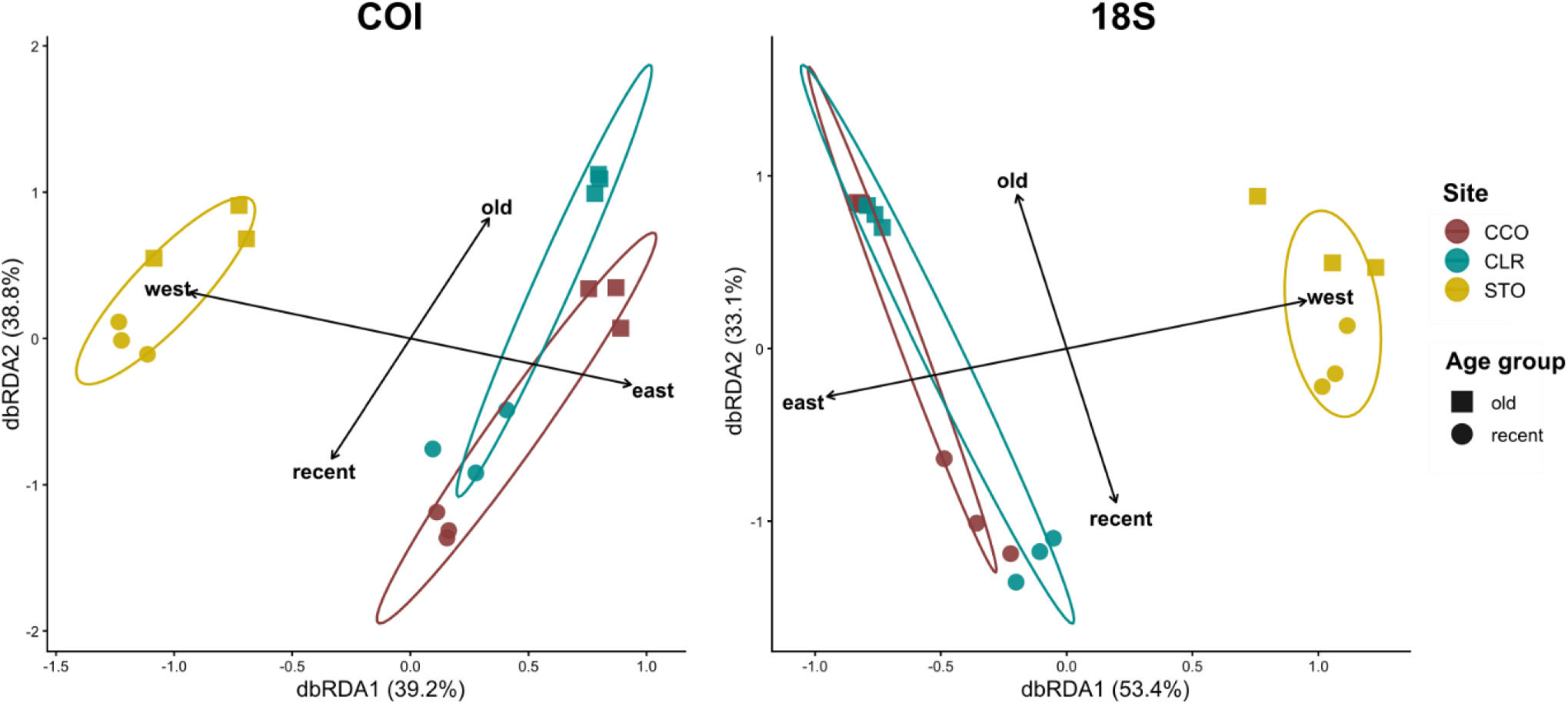
Distance-based redundancy analysis (db-RDA) of metabarcoding-based *seda*DNA community composition based on Bray–Curtis dissimilarities. Points represent individual samples coloured by site (CCO, CLR, STO) and shaped by sediment age group (recent vs. old). Ellipses show the 68% dispersion of samples within each site. The ordination is constrained by site and sediment age group, and the first two canonical axes (dbRDA1 and dbRDA2) are shown with their percentage of constrained variation explained. Arrows indicate fitted environmental contrasts derived from the metadata: “recent” versus “old” sediments (4– 5 cm vs. 45–50 cm) and “west” versus “east” region (STO vs. CCO/CLR).

**Fig. S12:**
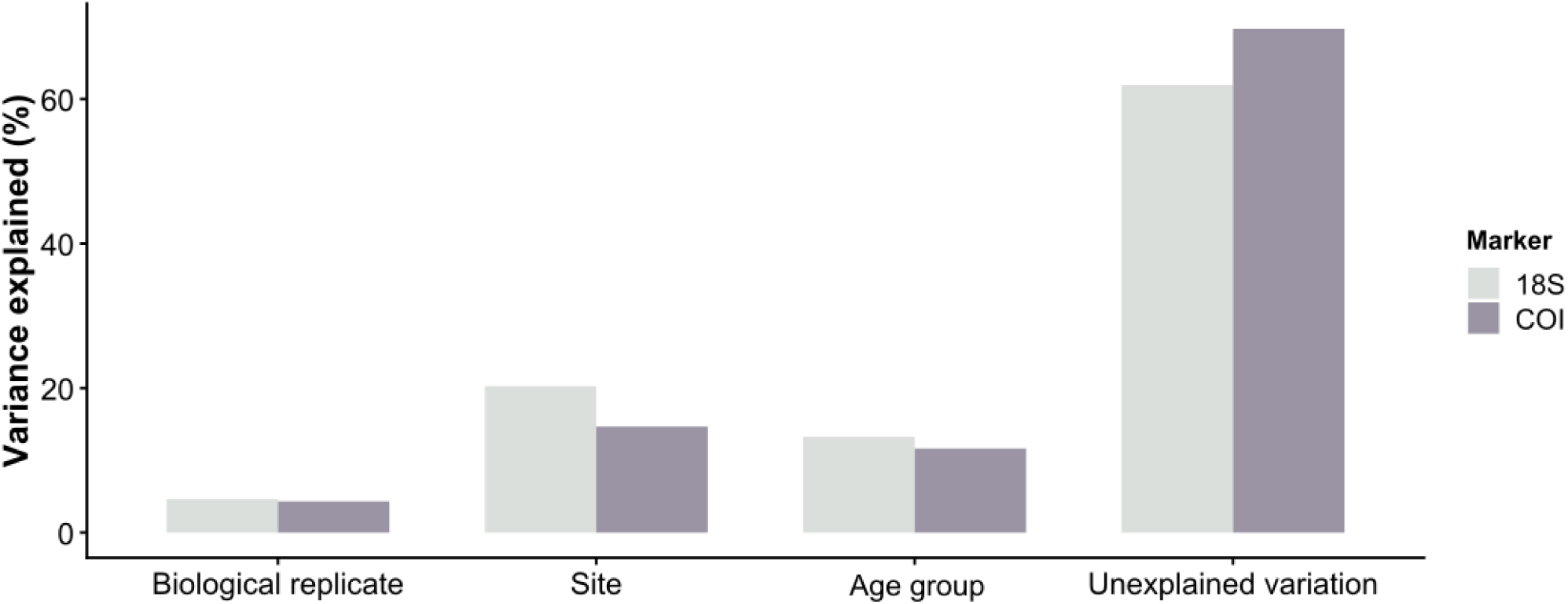
Bar plots showing marginal R² values from PERMANOVA models (adonis2, 999 permutations) illustrating the relative contribution of site, sediment age group, and biological replication to Bray–Curtis metazoan community dissimilarities for COI and 18S datasets. Marginal R² values represent the unique contribution of each factor after accounting for the others and should be interpreted comparatively rather than as additive proportions of explained variance.

**Fig. S13:**
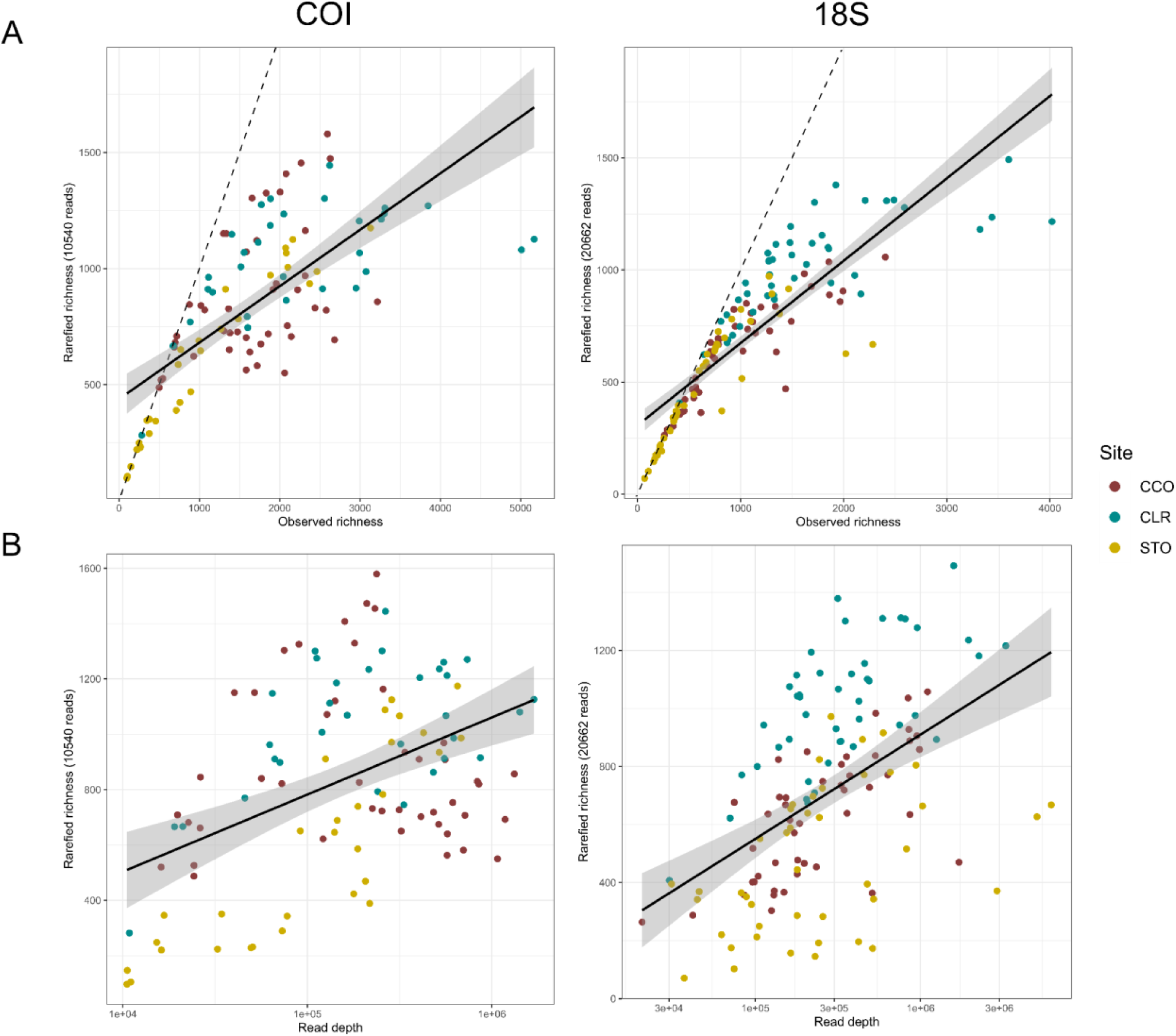
Sensitivity analysis evaluating the effect of sequencing-depth standardisation on ASV richness estimates. (A) Relationship between observed and rarefied richness for COI and 18S PCR replicates after sample-size rarefaction (10,540 reads for COI and 20,662 reads for 18S). (B) Relationship between rarefied richness and sequencing depth.

