## Supplementary figures and images for "Getting to the core of the matter – Assessing the role of replication in metabarcoding-based *seda*DNA"

### Figure S1

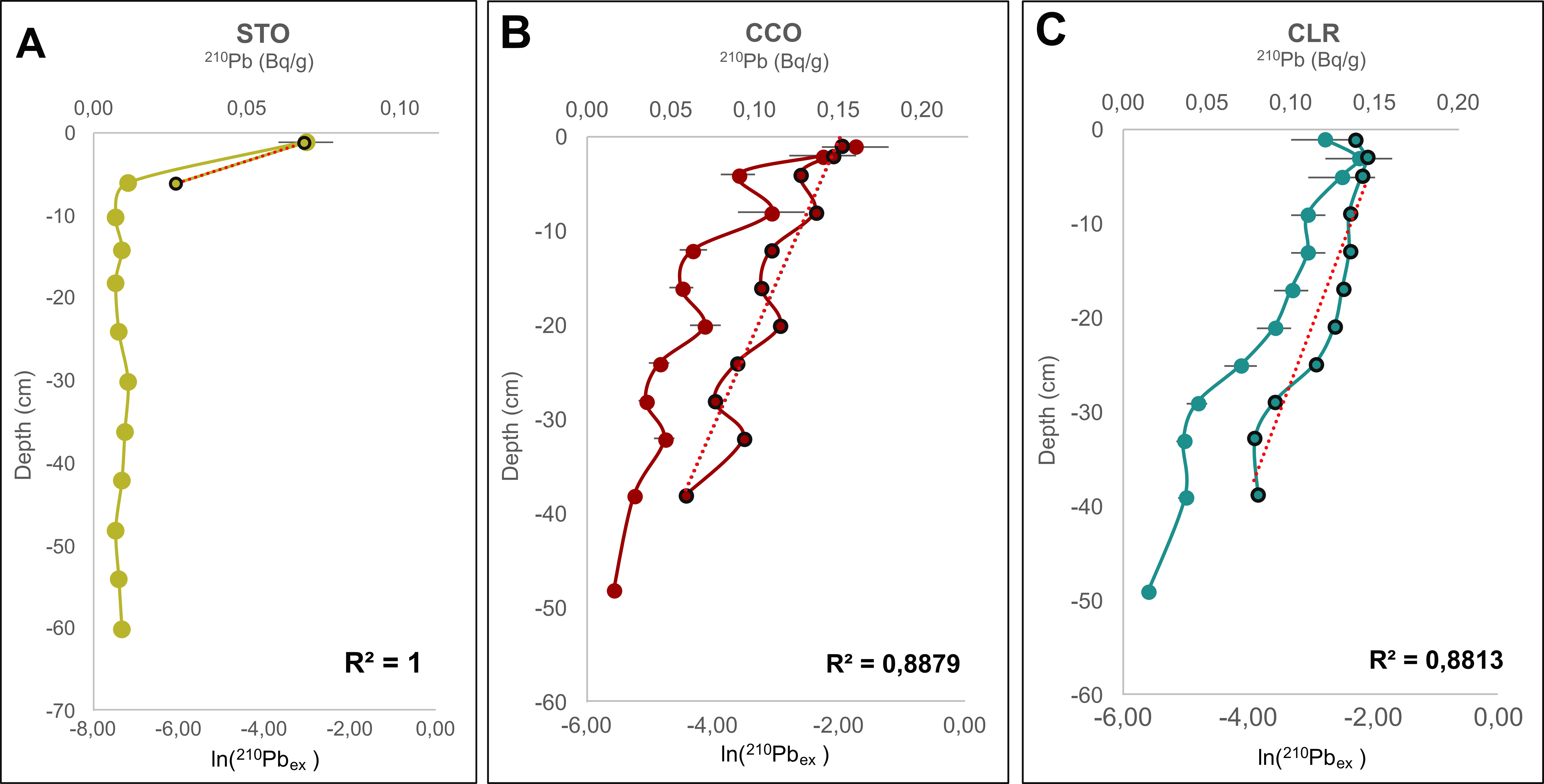

### Figure S2

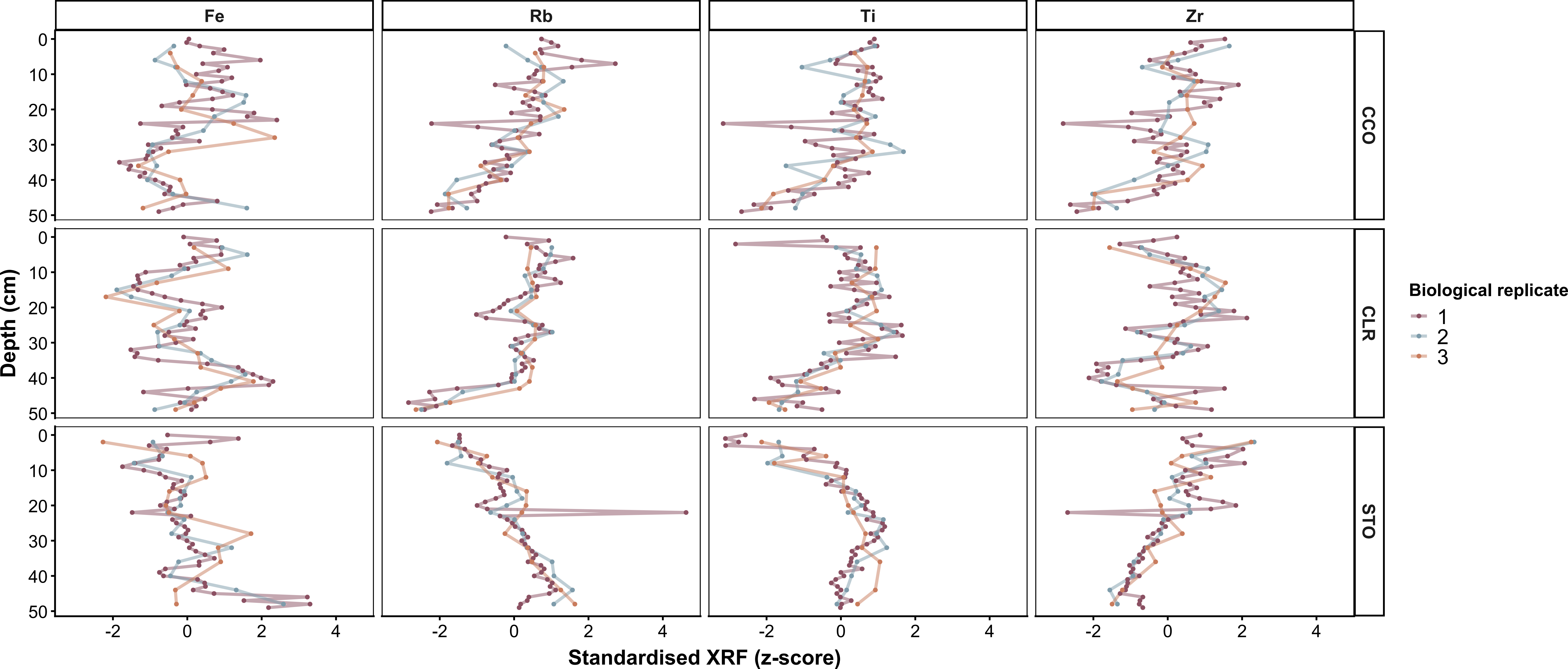

### Figure S3

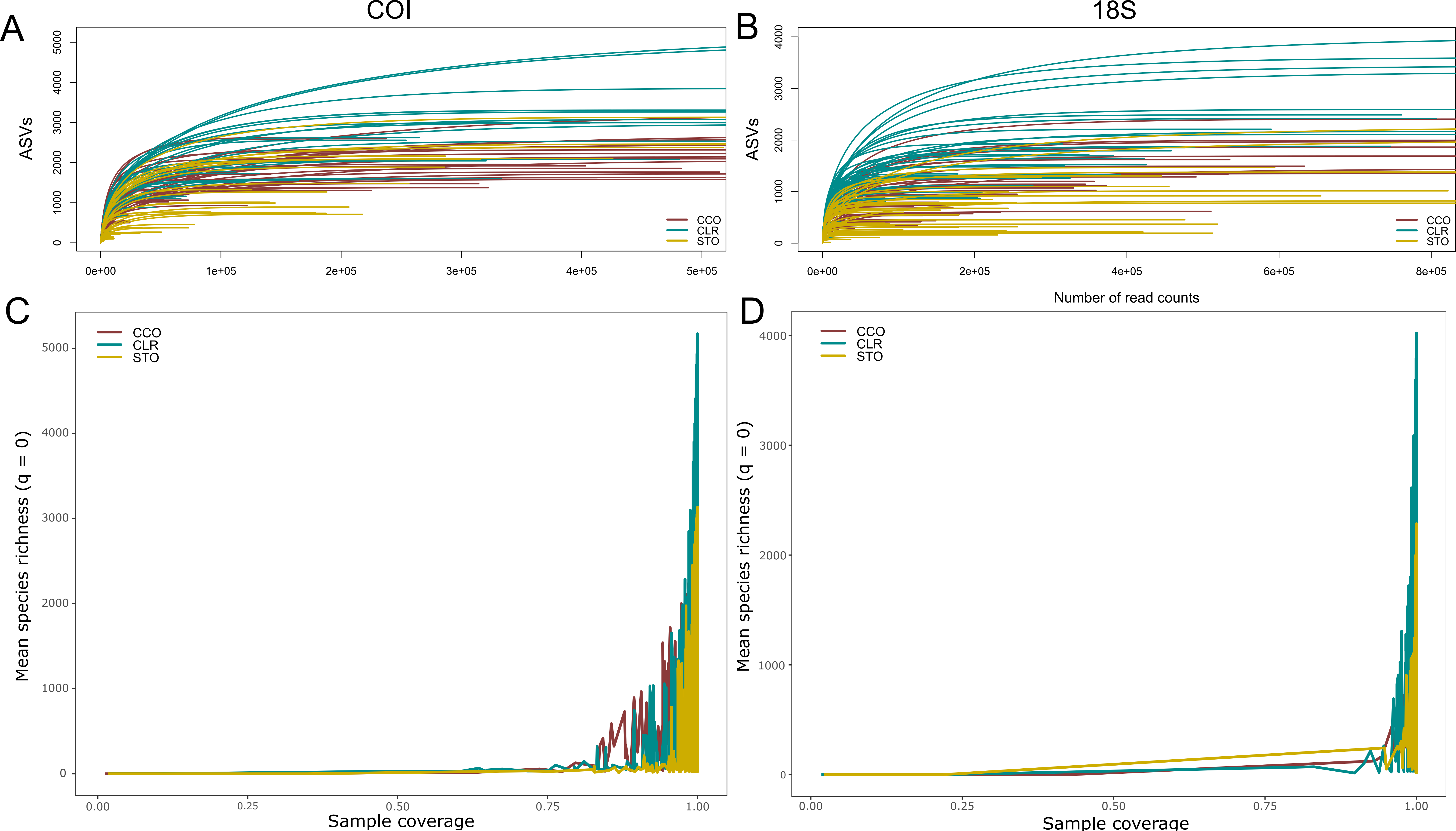

### Figure S4

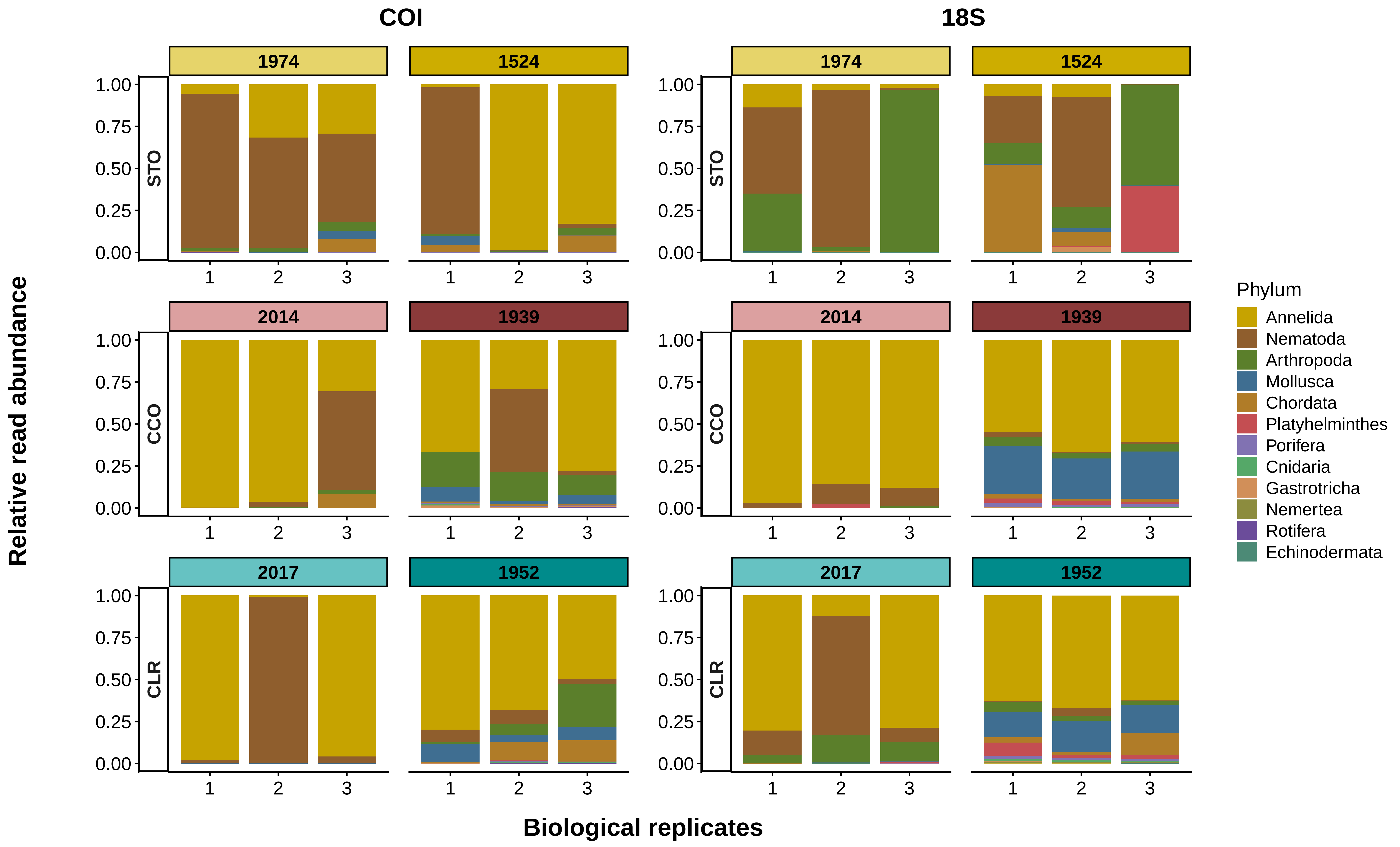

### Figure S5

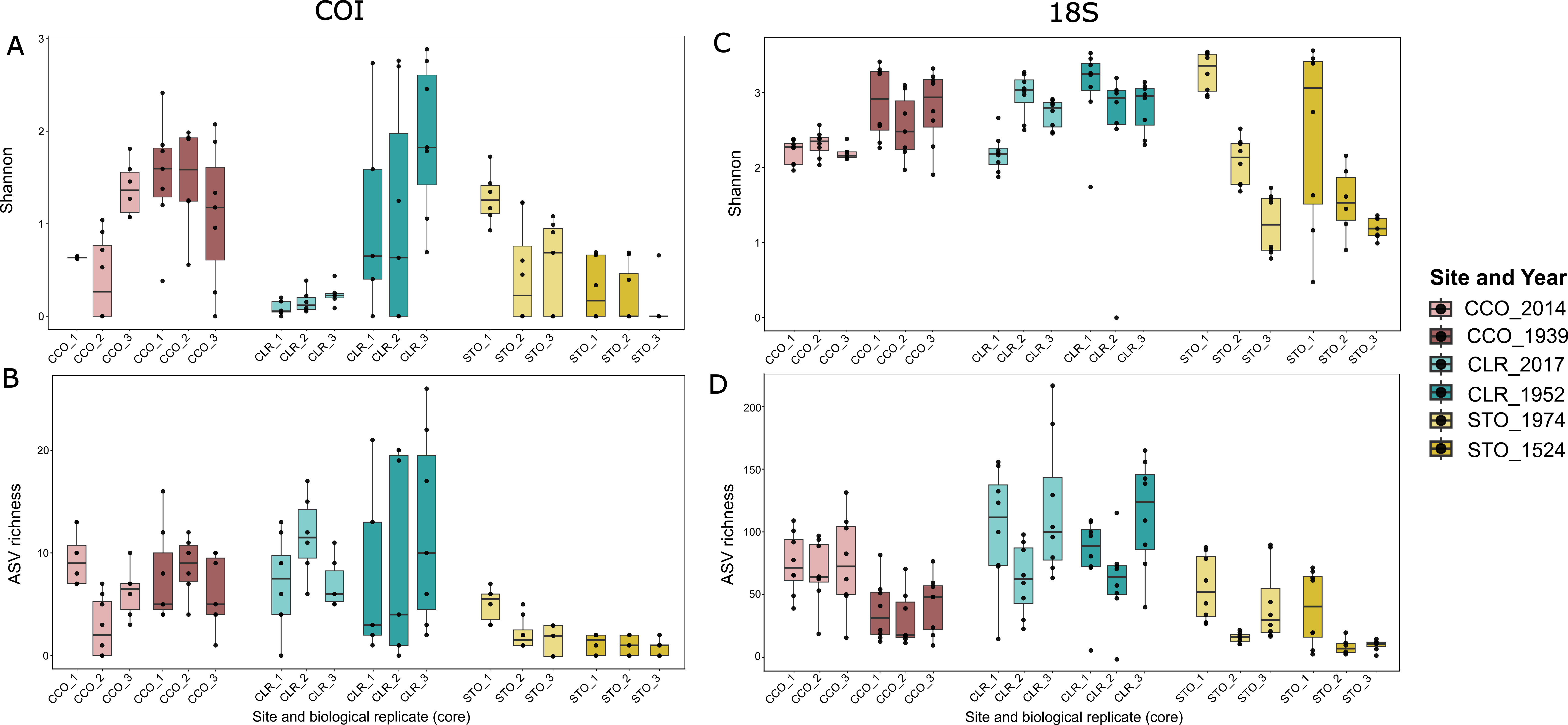

### Figure S6

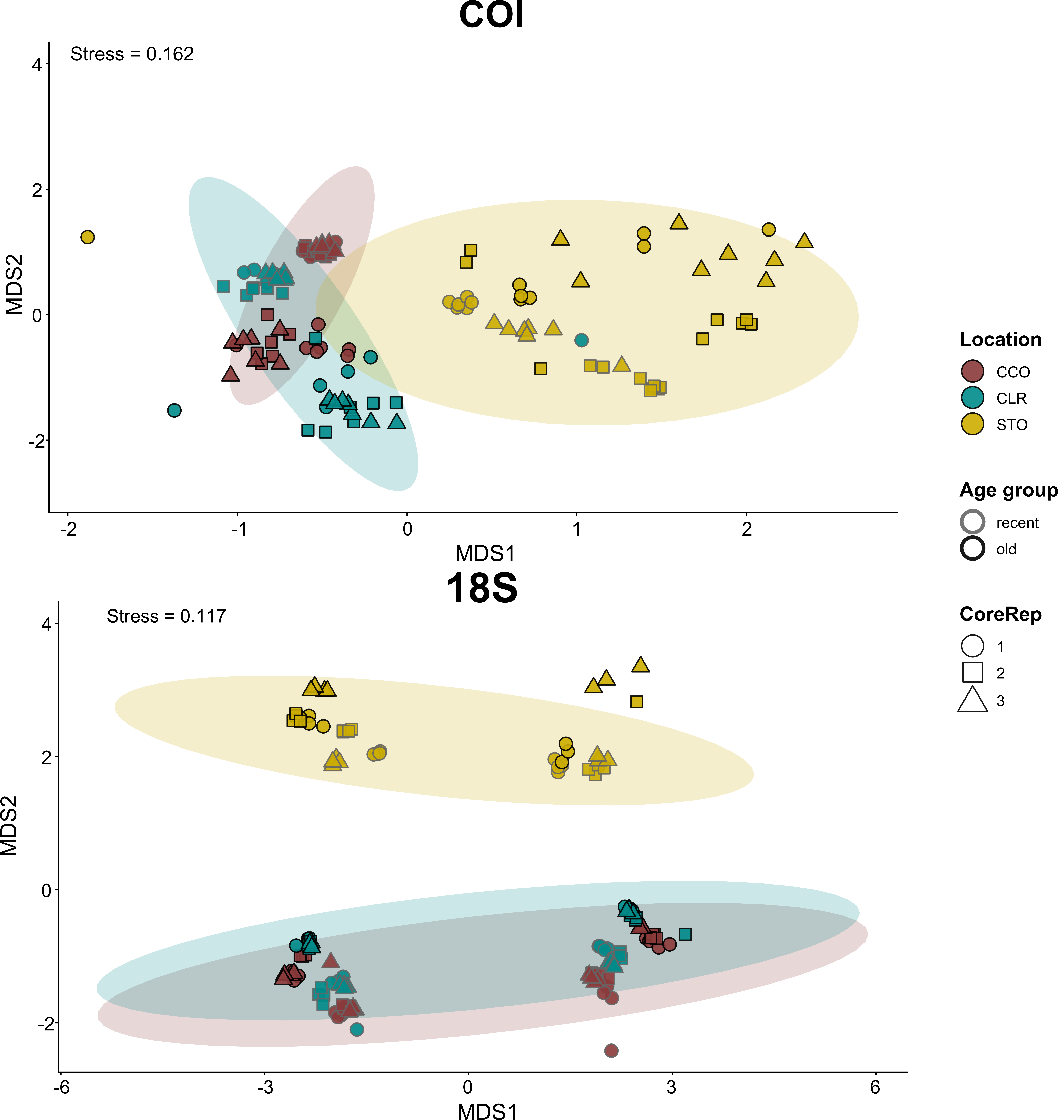

### Figure S7

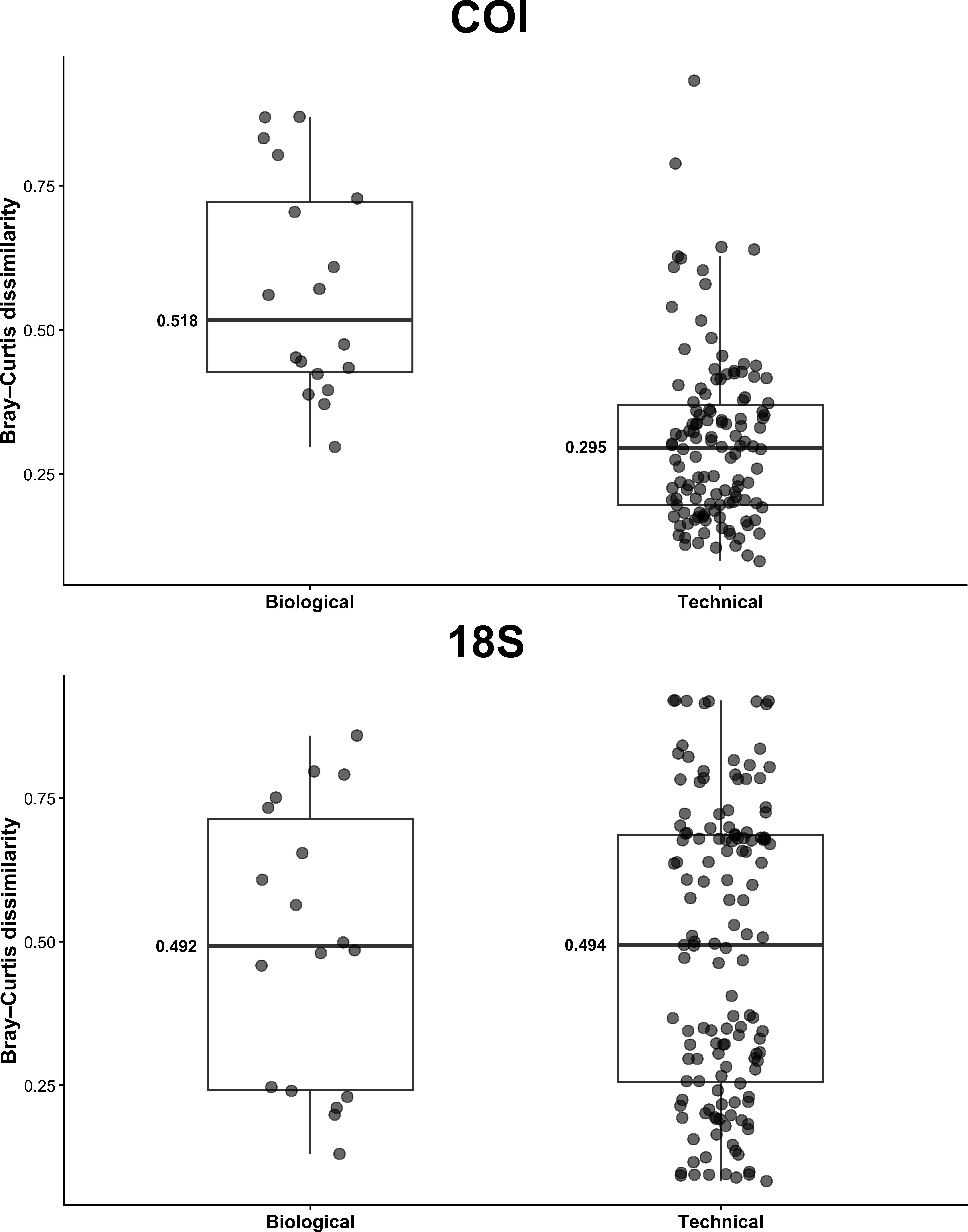

### Figure S8

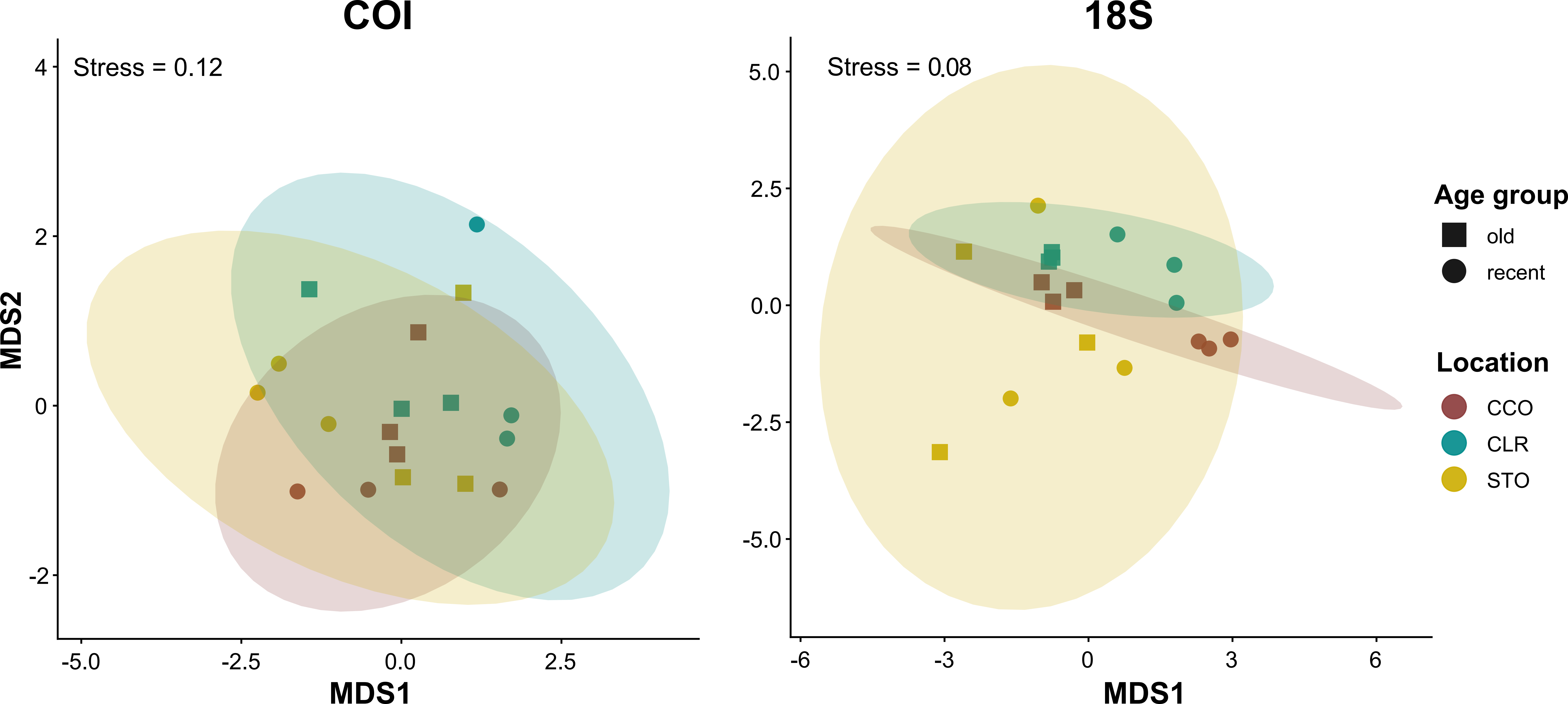

### Figure S9

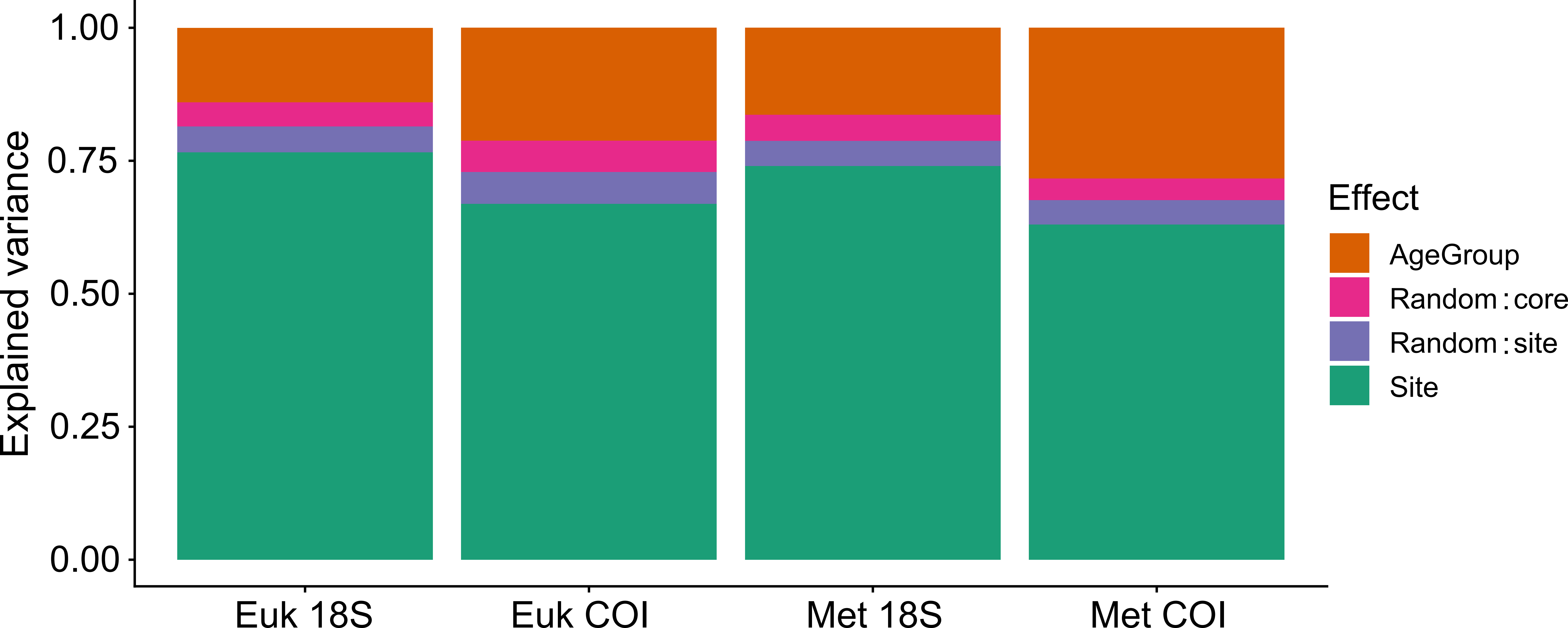

### Figure S10

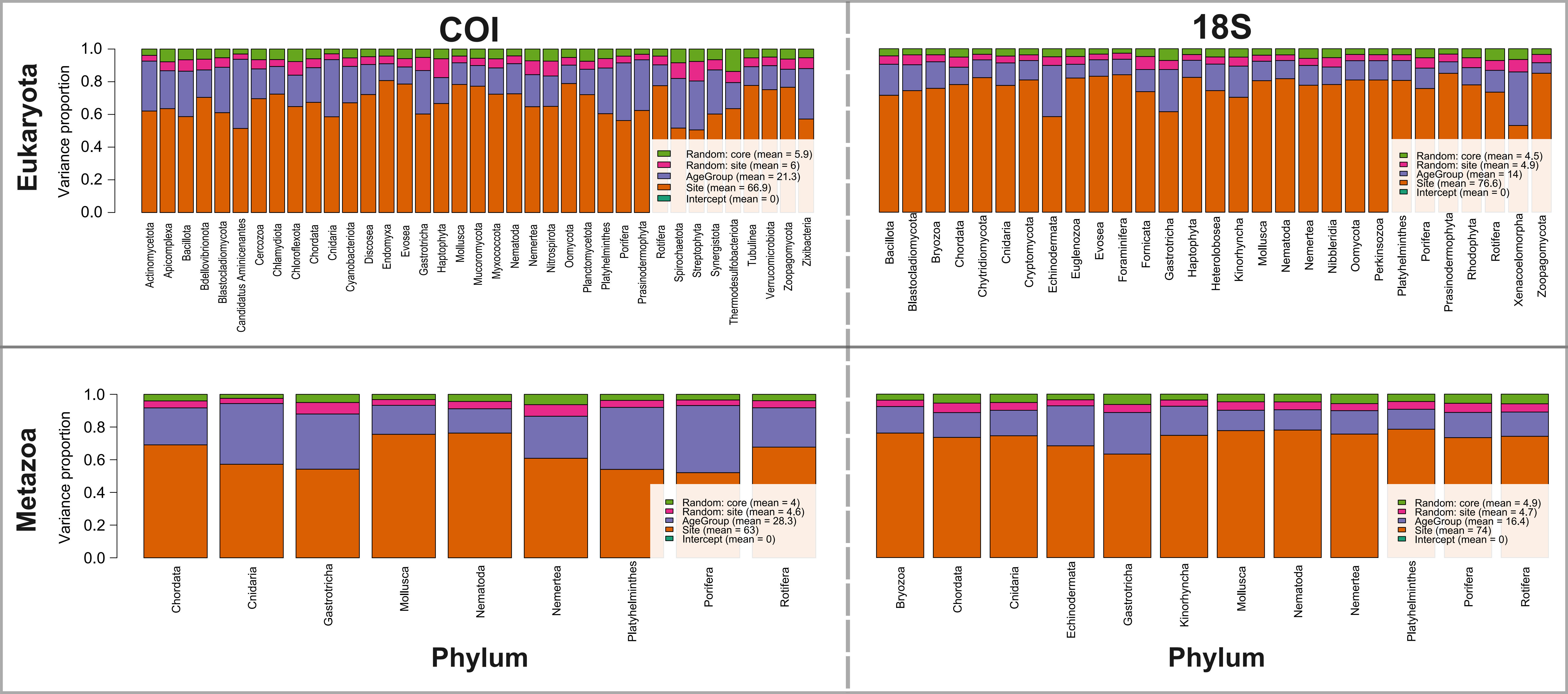

### Figure S11

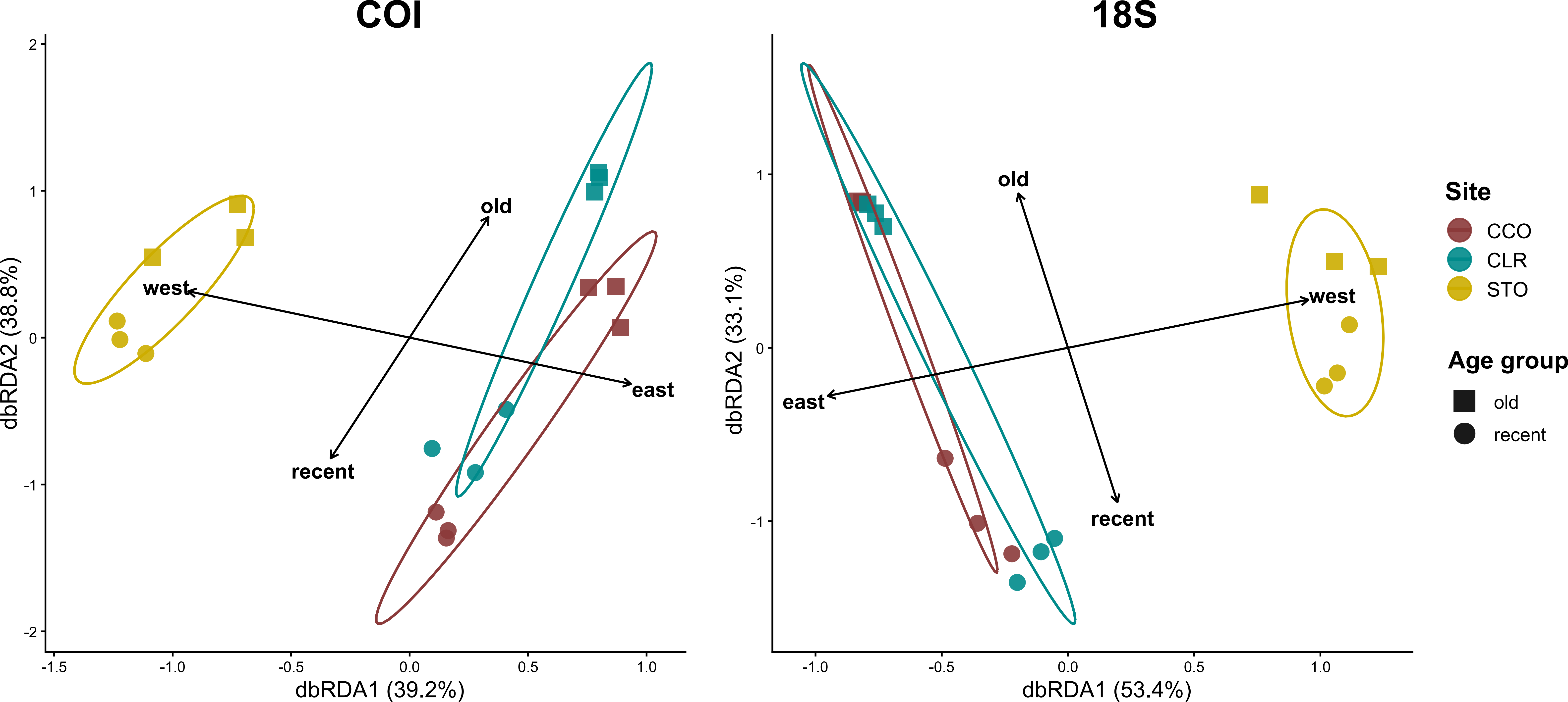

### Figure S12

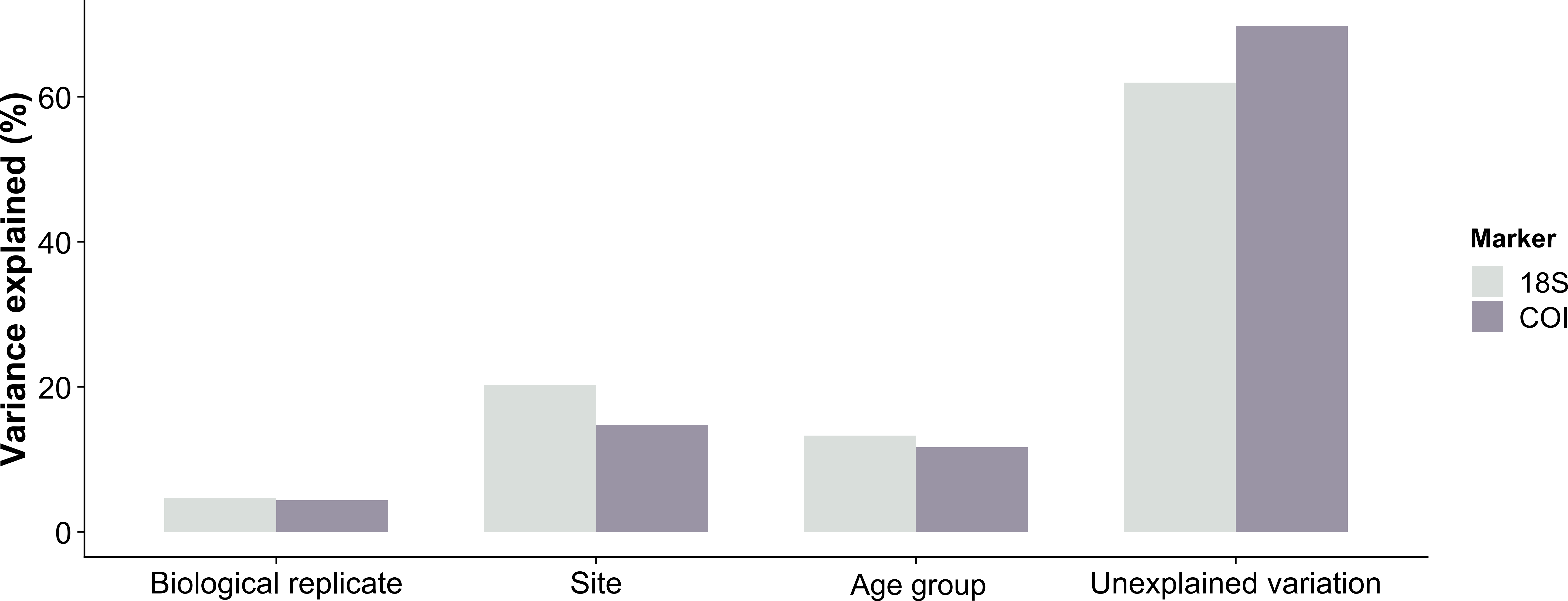

### Figure S13

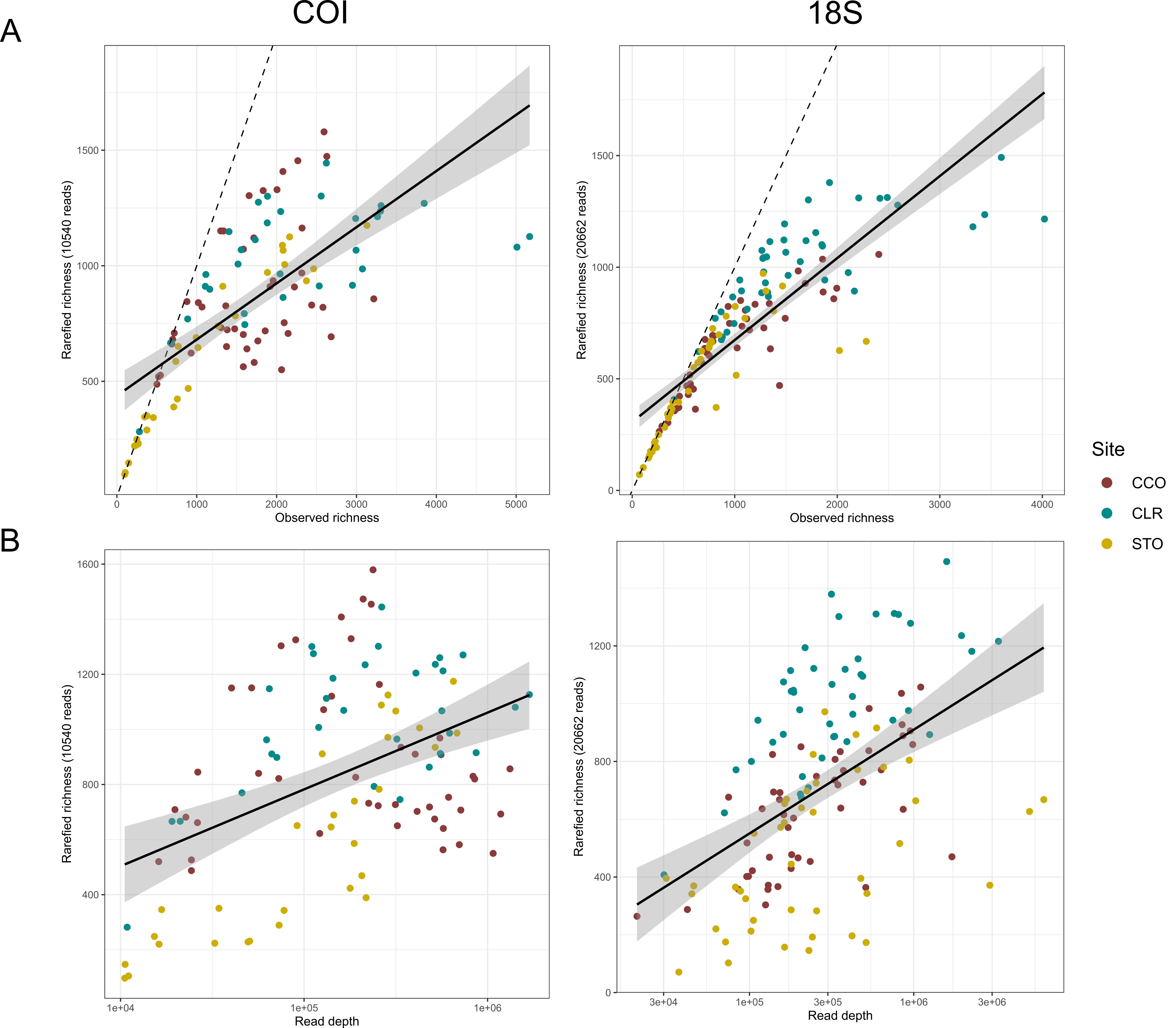
